# Trehalose-based coacervates for local bioactive protein delivery to the central nervous system

**DOI:** 10.1101/2023.10.05.561124

**Authors:** Laboni F Hassan, Riya Sen, Timothy M O’Shea

## Abstract

Therapeutic outcomes of local biomolecule delivery to the central nervous system (CNS) using bulk biomaterials are limited by inadequate drug loading, neuropil disruption, and severe foreign body responses. Effective CNS delivery requires addressing these issues and developing well-tolerated, highly-loaded carriers that are dispersible within local neural parenchyma. Here, we synthesized biodegradable trehalose-based polyelectrolyte oligomers using facile A2:B3:AR thiol-ene Michael addition reactions that form complex coacervates upon mixing of oppositely charged oligomers. Coacervates permit high concentration loading and controlled release of bioactive growth factors, enzymes, and antibodies, with modular formulation parameters that confer tunable release kinetics. Coacervates are cytocompatible with cultured neural cells *in vitro* and can be formulated to either direct intracellular protein delivery or sequester media containing proteins and remain extracellular. Coacervates serve as effective vehicles for precisely delivering biomolecules, including bioactive neurotrophins, to the mouse striatum following intraparenchymal injection. These results support the use of trehalose-based coacervates as part of therapeutic protein delivery strategies for CNS disorders.

## Introduction

Local delivery of bioactive proteins is being explored as part of experimental treatments for various central nervous system (CNS) disorders [1, 2]. In CNS injuries, such as spinal cord injury (SCI) and ischemic stroke, local delivery of neurotrophins including Nerve growth factor (NGF), Glial cell line-derived neurotrophic factor (GDNF), Brain-derived neurotrophic factor (BDNF), and Ciliary neurotrophic factor (CNTF) have been used to chemoattract regrowing axons across large complete lesions, guide connectivity of regrowing circuits, or recruit newly proliferated glia and neuroblasts to repair sites of tissue damage [3–10]. In models of neurodegeneration such as Parkinson’s and Alzheimer’s disease, local delivery of neurotrophins has helped arrest disease progression by improving neuron survival and preserving neural circuit function [11, 12], leading to their investigation in clinical trials [13]. To aid neuroplasticity after CNS injury, local delivery of other protein therapies such as extracellular matrix-modifying enzymes has been studied to augment relay circuit formation [14, 15]. Ultimately, as our understanding of the pathophysiology of CNS disorders continues to evolve, new protein-based therapeutic candidates will continue to be identified.

Despite their promise, clinical translation of protein-based therapies continues to be hampered by molecular delivery challenges that require new bioengineering solutions. Protein-based therapies do not readily cross the blood brain barrier, so obtaining therapeutic levels in the CNS requires high systemic doses that can be toxic [16] or have substantial off-target effects [17]. While local delivery of proteins to the CNS via high dose bolus intraparenchymal injections or indwelling catheter systems may overcome systemic toxicity issues, a lack of control over concentration and spatial distribution of delivered protein can cause local neurotoxicity [18, 19]. Additionally, rapid protein aggregation and proteolytic degradation can limit temporal stability *in vivo* [20]. Injectable or implantable biomaterial-based drug delivery systems can improve upon the spatiotemporal control of protein release to optimize therapeutic effects while minimizing off-target neurotoxicity [21].

Using biomaterials to deliver protein therapeutics has led to improved outcomes in preclinical models of stroke [9, 22], SCI [3, 4], and Parkinson’s disease [23]. However, the fragile nature of these protein-based therapies means that they are particularly susceptible to denaturation during encapsulation into biomaterials as well as during prolonged delivery from biomaterials *in vivo*. Practically, this means that most protein delivered from biomaterials has limited bioactivity beyond the first few days. Foreign body responses (FBRs) that are provoked by bulk biomaterial carriers when introduced into the CNS can also dramatically alter delivery outcomes [24]. Biomaterials that evoke a severe FBR in the CNS cause infiltration of macrophages and microglia and formation of a non-neural cell compartment, thus mediating preferential delivery to these cells rather than to neurons, and in practice, has resulted in reduced neurotrophin efficacy after stroke [24]. Additionally, since the CNS is a constrained anatomical site, implantation of bulk biomaterial carriers, even those amenable to minimally invasive injection, can displace considerable volumes of neural tissue causing CNS damage and dysfunction [24]. If the therapeutic potential of local protein delivery for CNS disorders is to be realized, new strategies are needed to: 1) improve loading and long-term stability of proteins within biomaterials, and 2) minimize the neural tissue disruption caused by biomaterials.

To improve the long-term stability and bioactive delivery of proteins, we and others have incorporated multivalent presentation of trehalose into polymers and hydrogels [25–28]. Trehalose is a non-reducing disaccharide that is produced by yeast and other extremophile organisms to effectively stabilize intracellular proteins, nucleic acids, and cell membranes under extreme environments [29]. The unique strength, spatial orientation, and symmetry of hydrogen bonding afforded by trehalose confers enhanced protein stabilization compared to other excipients [30]. Trehalose has been applied as a cryo-preservative agent in commercial production of cell-based therapies[31] and is included on the “generally regard as safe” (GRAS) list by the FDA. While we have recently incorporated trehalose into poly(trehalose-co-guanosine) glycopolymer hydrogels to alter the natural responses of glial cells after CNS injury [32], trehalose-based biomaterials have yet to be tested as part of local protein delivery systems applied to the CNS.

Given that bulk biomaterials such as hydrogels and scaffolds displace significant volumes of neural tissue upon implantation, different approaches are needed to develop a delivery system that ensures high protein loading capacity and retains protein bioactivity while also being capable of local dispersion into volumes of neural tissue. While polymer nanoparticles can be used to deliver protein therapies [33] and may be readily dispersible in neural tissue [34], it is challenging to prepare nanoparticles with sufficiently high protein concentrations needed for therapy, and formulation methods using harsh organic solvents can denature particularly fragile proteins [35, 36]. In this work, we developed and tested an alternative method for dispersible protein delivery to CNS tissue by using a new trehalose-based complex coacervate system. Complex coacervates are small liquid droplets formed as a result of liquid–liquid phase separation evoked by ionic complexation of polyelectrolytes [37]. These self-assembled structures can form homogeneous spherical droplets on the nanometer to micron scale [38], permit very high protein encapsulation [39–41], and may stabilize proteins directly, leading to enhanced bioactivity [42, 43]. These properties make complex coacervates promising candidates for local CNS drug delivery applications. However, most current coacervate technologies incorporate linear polyelectrolytes that are either non-biodegradable or that have minimal control over electrostatic interactions, which can hamper protein release from these systems.

Here, we hypothesized that biodegradable multivalent trehalose-based complex coacervates: (i) could be used to load, stabilize, and release high concentrations of proteins; (ii) would be readily dispersible within CNS tissue by non-invasive intraparenchymal injection, causing minimal disruption to healthy neural tissue; and (iii) would evoke a minimal FBR such that proteins can be delivered predominately to neurons and glia rather than to recruited peripherally derived immune cells. To test these hypotheses, we synthesized branched polyelectrolyte trehalose-based oligomers by thiol-ene Michael addition, investigated the formulation conditions needed to prepare stable coacervates, evaluated protein encapsulation and bioactivity stabilization capacity of coacervates *in vitro,* assessed how coacervates can be formulated to permit intracellular protein delivery to neural cells *in vitro*, and tested coacervate-based delivery of reporter molecules and bioactive neurotrophins *in vivo* in the healthy mouse striatum. Our findings demonstrate the feasibility of using trehalose-based coacervates for enhanced local protein delivery to the CNS.

## Results

### Multifunctional anionic and cationic trehalose-based branched oligomers can be synthesized by Thiol-ene Michael addition

Multivalent presentation of trehalose within covalently crosslinked hydrogels effectively stabilizes biomacromolecules during controlled release [27]. Here, we sought to repurpose similar chemistry to generate complex coacervates to enable bioactive protein delivery to volumes of neural tissue with minimal tissue disruption that is otherwise associated with injection of bulk biomaterials[24]. In our previous trehalose-based hydrogel work, we synthesized trehalose diacrylate through regiospecific acylation of 6 or 6′ primary hydroxyl sites by vinyl acrylate in acetone using Candida antarctica Lipase B (CALB)-immobilized on acrylic resin as an enzymatic catalyst [27]. This reaction achieves diacrylate functionalization of trehalose in a single reaction without requiring protection/deprotection steps. However, vinyl acrylate is expensive and is currently difficult to source. We refined the trehalose diacrylate synthesis methodology to increase reaction yields and improve affordability by making the following adjustments: i) substituting vinyl acrylate with 2,2,2-Trifluoroethyl acrylate (TFEA) (TFEA is ∼7% the cost of vinyl acrylate per gram); ii) using anhydrous trehalose instead of trehalose dihydrate; and iii) using 2-methyl-2-butanol (2M2B) instead of acetone as a reaction solvent (Figure 1A). Whereas transesterification with vinyl acrylate affords an unstable enol leaving group upon acylation of trehalose, effectively making the reaction irreversible [44], the use of TFEA introduces the possibility of the reverse competing hydrolysis reaction taking place during trehalose acylation. However, with TFEA, the three electron withdrawing fluorine groups reduce the nucleophilicity of the 2,2,2-trifluoroethanol leaving group such that we readily derived the trehalose diacrylate product (Figure 1A, S1) [45]. Solubilizing anhydrous trehalose in DMSO before combining with 2M2B also increased solubility of trehalose in the organic media and improved the overall yield of trehalose diacrylate. As before, trehalose diacrylate was readily purified by silica flash chromatography to yield the pure product (Figure 1A, S1) and required the addition of 4-methoxyphenol (MEHQ) to prevent free-radical homopolymerization during work-up [27].

**Figure 1.**
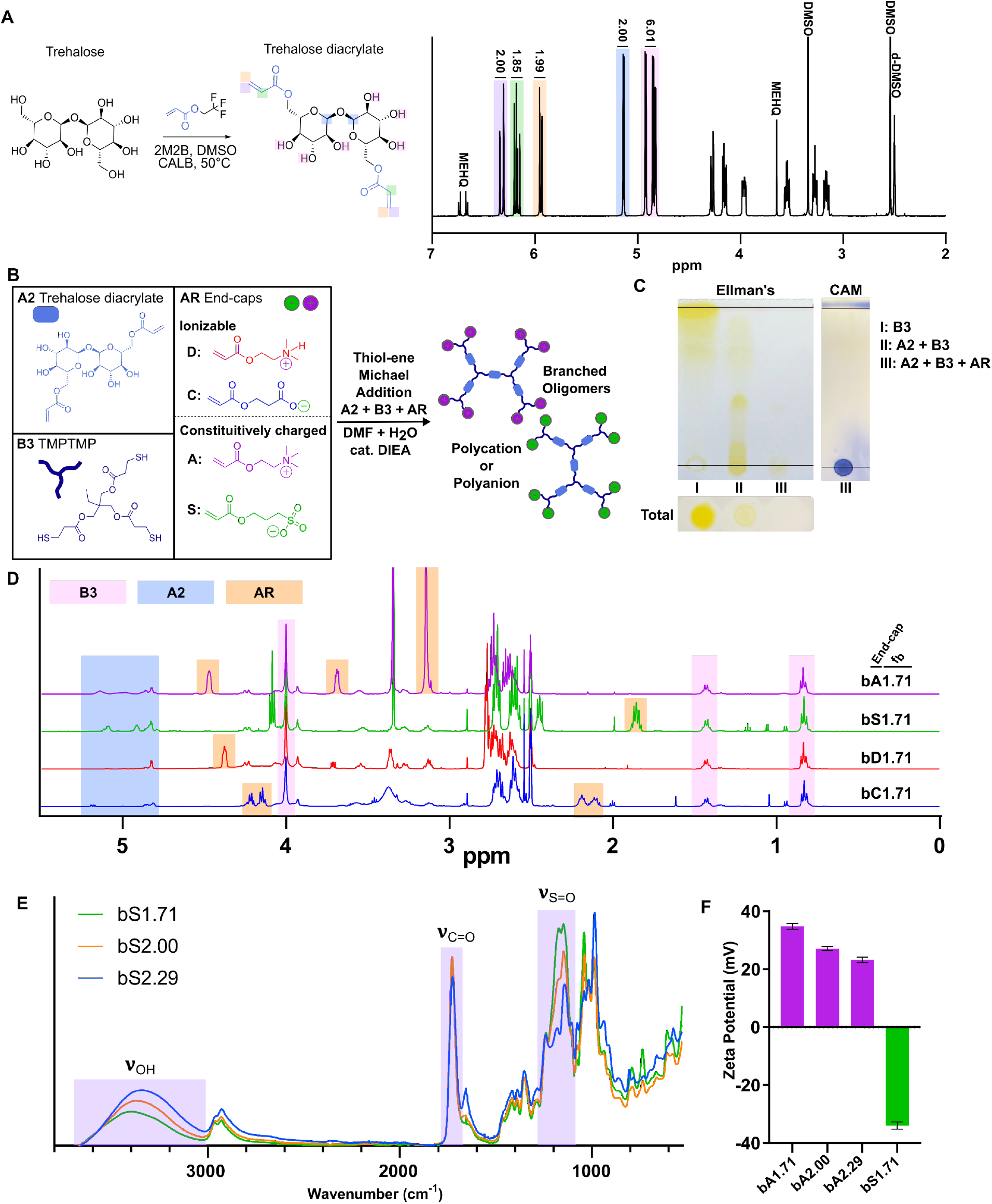
Synthesis and characterization of trehalose-based oligomers. A) Synthesis scheme for enzymatically catalyzed acylation reaction to prepare trehalose diacrylate and corresponding ^1^H-NMR with characteristic peaks and associated integrations labeled. B) Synthesis scheme for base-catalyzed A2+B3+AR thiol-ene Michael addition reactions to form trehalose-based oligomers. A: [2-(Acryloyloxy)ethyl] trimethylammonium chloride, S: 3-Sulfopropyl acrylate, D: 2-(Dimethylamino)ethyl acrylate, C: 2-Carboxyethyl acrylate. C) TLC with Ellman’s and CAM stains indicating progression of thiol-ene reactions after addition of each monomer. D) ^1^H-NMR of oligomers of the same branching functionality (f_b_) with each of 4 end-caps with characteristic peaks highlighted. E) FTIR of oligomers at 3 different f_b_ prepared with S end-cap with characteristic IR stretches of select chemical bonds highlighted. F) Zeta potential measurements of constitutively positive bA oligomers at 3 different f_b_ and a constitutively negative bS oligomer. Graph shows mean ± SEM with n=6 cycles per group.

Next, we synthesized trehalose-based branched oligomers using an A2+B3+AR reaction scheme [46] involving base-catalyzed thiol-ene Michael addition step-growth between trehalose diacrylate (A2), trimethylolpropane tris(3-mercaptopropionate) (TMPTMP) (B3), and various charged monoacrylate end-capping species (AR) (Figure 1B). We investigated 4 different end-cap species: i) 2-carboxyethyl acrylate (C), and ii) 2-(dimethylamino) ethyl acrylate (D) which are ionizable monomers with functional groups that have pKa values within two to three units of physiological pH making them partially charged species under such conditions; as well as iii) 3-sulfopropyl acrylate (S) and iv) 2-acryloxyethyl trimethyl ammonium chloride (A) which are constitutively charged monomers (Figure 1B). Branched oligomers functionalized with C or S end-caps are polyanions while D or A end-cap functionalization generates polycation oligomers. Nomenclature for oligomers followed the following convention: “b” for branched; “C”, “D”, “S”, or “A”, to denote end-cap functionality; and a numerical value corresponding to the intended branching functionality (f_b_) of the synthesis (Figure S2). Branched oligomer synthesis was performed in one-pot at a concentration of 100 mg/ml in a DMF/water mixed solvent with N,N-Diisopropylethylamine (DIEA) as a catalytic base to initiate the reaction (Figure 1B). Combining end-capping monoacrylates (AR) at the onset of the reaction effectively avoids crosslinking and gelation of A2+B3 systems at high conversion[47]. Irrespective of the end-cap species used, oligomer reactions proceeded with complete consumption of thiol and acrylate groups, as monitored by TLC developed with Ellman’s and CAM reagents (Figure 1C) as well as by NMR (Figure 1D, S3-4). We first prepared oligomers with each end-cap at a fixed stoichiometric ratio of 0.5:1:2 (A2:B3:AR) for a f_b_ of 1.71. Adjusting the reaction stoichiometry to 1:1:1 and 1.375:1:0.25 (A2:B3:AR), to increase the f_b_ to 2.00 and 2.29 respectively, was associated with increased trehalose content and a reduction in the proportion of end-caps on the oligomer as measured by FTIR and NMR (Figure 1E, S4,S5). Number average molecular weights (Mn) for oligomers estimated by NMR ranged from approximately 2 kDa for f_b_ of 1.71 to 15 kDa for f_b_ of 2.29 with the oligomer molecular weights altered minimally by the different end-cap species (Figure S6). MALDI-TOF analysis of ionizable D end-capped oligomers showed higher intensities of molecular weight oligomeric species at 4 and 5kDa as the f_b_ increased from 1.71 to 2.29 and a concurrent reduction in the intensity of lower molecular weight species in the 1-2 kDa range (Figure S7). The f_b_ of 2.29 was determined to be the upper limit of oligomer branching permitted without inducing insoluble crosslinked networks, but significant intramolecular cyclization likely still occurs at this functionalization [46]. Consistent with chemical characterization of branching properties, constitutively charged oligomers with A or S end-caps had positive and negative zeta potentials respectively that were directly proportional to the stoichiometrically controlled density of end-cap species on the oligomer (Figure 1F). To purify oligomers, reaction solutions were concentrated *in vacuo* and acidified or alkalized in aqueous solutions depending on the end-cap to dissolve the oligomer in the aqueous phase, followed by liquid-liquid extraction with ethyl acetate to remove DIEA, DMF, and any low molecular weight or minimally end-capped oligomers. Yield after purification and lyophilization was typically 60-90%. These data show that branched trehalose oligomers of different charges could be readily synthesized at high scale using A2+B3+AR thiol-ene Michael addition.

### Aqueous solubility of trehalose-based branched oligomers is altered by salt & pH

Complex coacervates are formed through liquid–liquid phase separation caused by association of two oppositely charged polyelectrolytes[48]. Formation of coacervates is conditional on each polyelectrolyte maintaining aqueous solubility prior to mixing, but polyelectrolyte solubility can be dramatically altered by salt and pH conditions. Therefore, we next evaluated how salt concentration and pH of aqueous buffers affected the solubility of trehalose-based branched oligomers with different f_b_ and end-cap functionalization. All oligomers with f_b_ ranging from 1.71-2.29 irrespective of end-cap functionalization could be readily dissolved in unbuffered water at 500 mg/ml. To test salt effects on solubility of ionizable versus constitutively charged oligomers, we screened differently branched oligomers with either A or D cationic end-caps under NaCl concentrations ranging from 0-500 mM. Solubility was screened using spectrophotometric turbidity measurements with absorbance values greater than 0.5 units defined as insoluble (Figure 2A). At 10 mg/ml, highly branched (f_b_ = 2.29) oligomers were insoluble after addition of only a small amount of salt, indicating that oligomers with minimal charged end-caps, high branching, and possible intramolecular cyclization were readily affected by salt screening. Oligomers with f_b_ = 1.71 remained soluble up to salt concentrations of 500 mM with no differences detected between the two different cationic end-caps. All oligomers with f_b_ = 2.00 were soluble up to physiological salt concentrations (∼150 mM) irrespective of chemical composition of the end-cap (Figure 2D).

**Figure 2.**
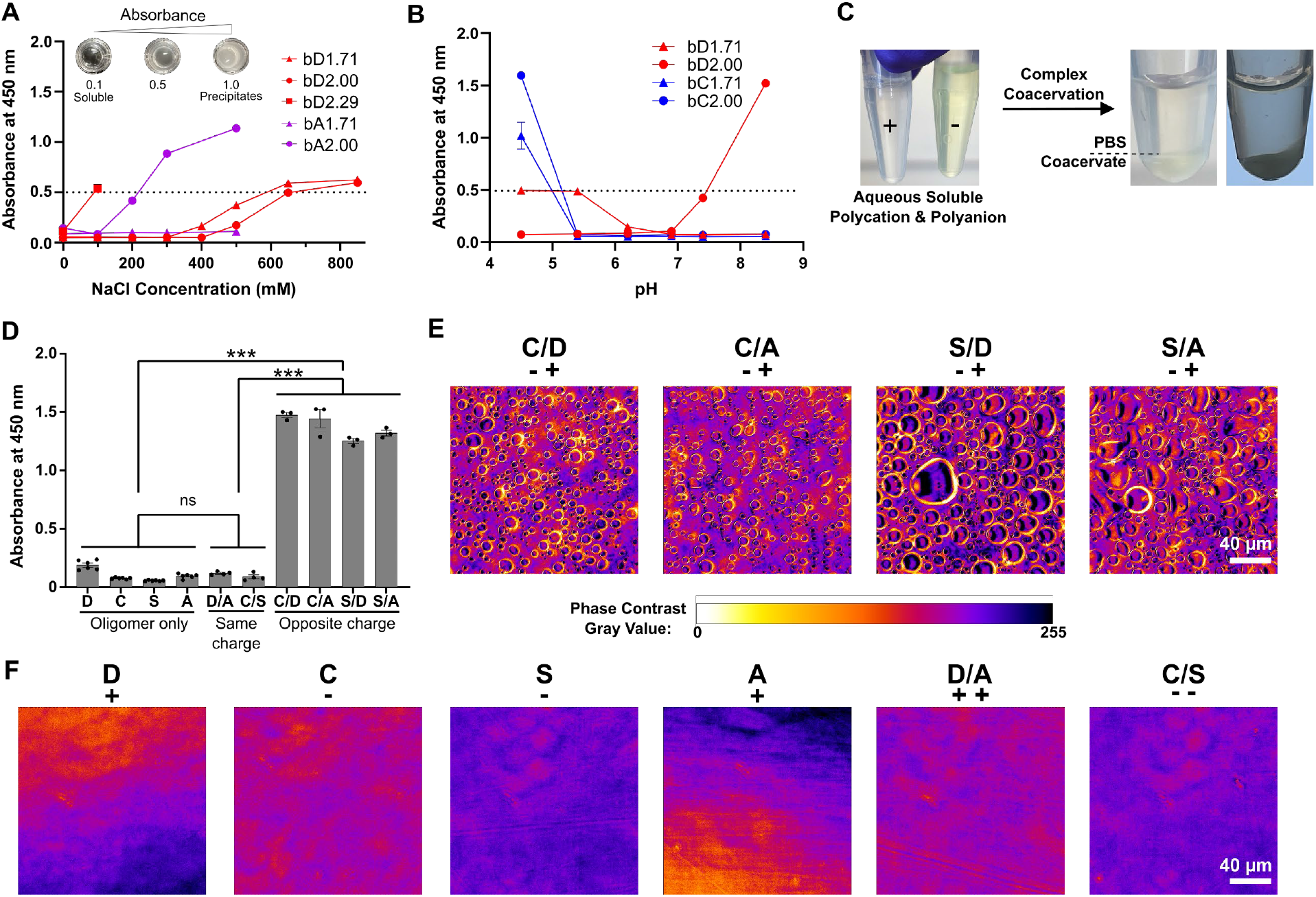
Solubility and coacervate formation of trehalose-based oligomers. A) Solubility of oligomers (10 mg/ml) under increasing NaCl concentrations determined by spectrophotometric turbidity measurements. B) pH dependent changes in solubility of ionizable oligomers (10 mg/ml). C) Photographic images demonstrating the formation of coacervates from mixing of soluble oligomers by liquid-liquid phase separation. D) Solubility of branching functionality (f_b_) = 2.00 oligomers (10 mg/ml) and all combinations of end-caps at 1:1 molar ratios in isotonic phosphate buffer (pH 6). Not significant (ns) and *** P < 0.0001, one-way ANOVA with Tukey’s multiple comparison test. E) Pseudo-colored phase contrast images of coacervates formed from each combination of polycationic and polyanionic oligomers at f_b_ = 2.00. Oligomers are abbreviated by end-cap. F) Pseudo-colored phase contrast images of soluble oligomers and same-charged combinations of oligomers at f_b_ = 2.00. Oligomers are abbreviated by end-cap. (A, B) Threshold Abs450 = 0.5 set at point of visible turbidity. All graphs show mean ± SEM with n=3-6.

For pH experiments, isotonic pH adjustable buffers were used to derive solutions with pH buffering capacity ranging from 4.5-8.5. Polycationic and polyanionic oligomers with f_b_ up to 2.00 were solubilized in water at 500 mg/ml and diluted into different pH buffers at a final concentration of 10 mg/ml. Oligomers with constitutively charged A and S end-caps displayed no pH dependent solubility changes as expected (Figure S8). D end-capped oligomers with f_b_ = 2.00 (bD2.00) were fully soluble under mildly acidic conditions (pH <6.5) where the end-cap tertiary amines are essentially fully protonated (Figure 2B). bD2.00 displayed an apparent pKa of 7.99 in aqueous buffers, and at pH values greater than this apparent pKa, the oligomer started to become insoluble (Figure 2B, S9). With lower f_b_, bD1.71 readily maintained solubility across the tested pH range, and was attributed to a higher net charge due to more end-cap species on the oligomer, rather than an alteration to the effective pKa. bC2.00 was soluble under physiologic pH and mildly basic conditions, but this oligomer formed insoluble precipitates under acidic (pH<5) conditions (Figure 2B). Just like for D end-capped oligomers, C end-capped oligomers with lower branching and higher end-cap functionality showed a more extended pH range in which they were soluble (Figure 2B). These data show that branched trehalose-based oligomers require a minimum effective charge by electrolytic dissociation of end-caps for oligomers to maintain aqueous solubility and that charge neutralization can provoke immiscibility of highly branched trehalose-based oligomers. Such charge dependent solubility behaviors are important factors in driving phase separation and coacervation versus soluble complex formation upon interaction of oppositely charged polyelectrolytes.

### Trehalose-based coacervates form by mixing oppositely charged oligomers

Next, we screened for complex coacervate formation by mixing different combinations of oligomers with f_b_ = 2.00 in isotonic phosphate buffered saline at pH = 6.0. Oligomer f_b_ was chosen to maximize trehalose content on the oligomers to confer the greatest potential protein stabilizing effect in resulting coacervates while maintaining solubility in isotonic conditions and minimizing intramolecular cyclization (Figure S6) that occurs with highly branched oligomers. Buffer conditions were selected based on prior salt and pH solubility results to optimize charge density for bC and bD oligomers before mixing. As before, all individual oligomers were soluble as determined by spectrophotometric turbidity measurements (Figure 2C,D). Combining two different oligomers of the same charge type, namely bD2.00+bA2.00 or bC2.00+bS2.00, at a 1:1 molar ratio did not evoke phase separation or coacervation (Figure 2D,F). However, upon mixing any two oppositely charged oligomers under the same conditions, we observed rapid liquid-liquid phase separation that resulted in significantly increased turbidity and formation of coacervate droplets with sizes ranging from 5-50 µm (Figure 2D,E). While all 4 combinations of oppositely charged oligomers (bC2.00+bD2.00, bC2.00+bA2.00, bS2.00+bD2.00, bS2.00+bA2.00) formed coacervates with equivalent relative turbidity, there were some detectable variations in the size distribution and shape of the coacervates immediately after mixing (Figure 2E). Irrespective of these minor initial variations, all coacervates formulated at this concentration (10 mg/ml) coalesced into a bulk liquid phase within 30 minutes (Figure 2C). Based on these data and the understanding that the ionizable carboxylate and tertiary amine chemical functionalities are likely to be better tolerated *in vitro* and *in vivo*[49],[50], we opted to explore coacervates derived from oligomers end-capped with C and D functionality for the remainder of the studies reported here.

To map the formulation space and better understand factors that alter complex coacervate derivation using trehalose-based branched oligomers, we evaluated the effect of modifying oligomer concentration, charge ratio, salt concentration, and pH buffering conditions (Figure 3). At a 1:1 molar ratio of bC2.00 and bD2.00, a minimum concentration of 1 mg/ml of total oligomer was required to generate microscopically detectable coacervates, with the size of initially formed coacervate droplets increasing with higher oligomer concentrations up to 25 mg/ml (Figure 3A,B). Coacervate formation was minimally impacted by minor perturbations in the molar ratio of bC2.00 to bD2.00 on either side of 1:1 stoichiometric equivalency at a fixed total oligomer concentration of 10 mg/ml. With increasing divergence from stoichiometric equivalency, resultant coacervate sizes decreased by approximately 10-fold. At very high charge imbalances (e.g. > 16:1), coacervates did not form and a clear solution persisted upon mixing (Figure 3C,D). There was a significant effect of solution salt concentration on coacervate size with a near two-fold decrease in overall coacervate diameter being detected as buffer salt concentration was increased from 0 to 500 mM (Figure 3E,F). There was minimal effect of pH on resultant coacervates formulated within a pH range of 6 to 8 (Figure 3G,H). Coacervates formulated at high concentrations (>5 mg/ml) in PBS resulted in coalescence of small spherical droplets into a bulk contiguous liquid coacervate phase (Figure 3I). This bulk liquid coacervate phase could be readily resuspended and dispersed into smaller microdroplets of less than 20 µm in diameter by simply diluting with PBS (Figure 3I). The viscous, liquid-like properties of the coalesced bulk coacervate phase was confirmed by Fluorescence recovery after photobleaching (FRAP) studies (Figure S10). Dilute coacervate solutions (<1 mg/ml) that did not display microscopic sized coacervates formed stable nanodroplets that were detectable only by dynamic light scattering (DLS), with coacervate size decreasing proportionally with coacervate concentration from >1000 nm at 1 mg/ml to less than 500 nm at 0.0625 mg/ml (Figure 3J).

**Figure 3.**
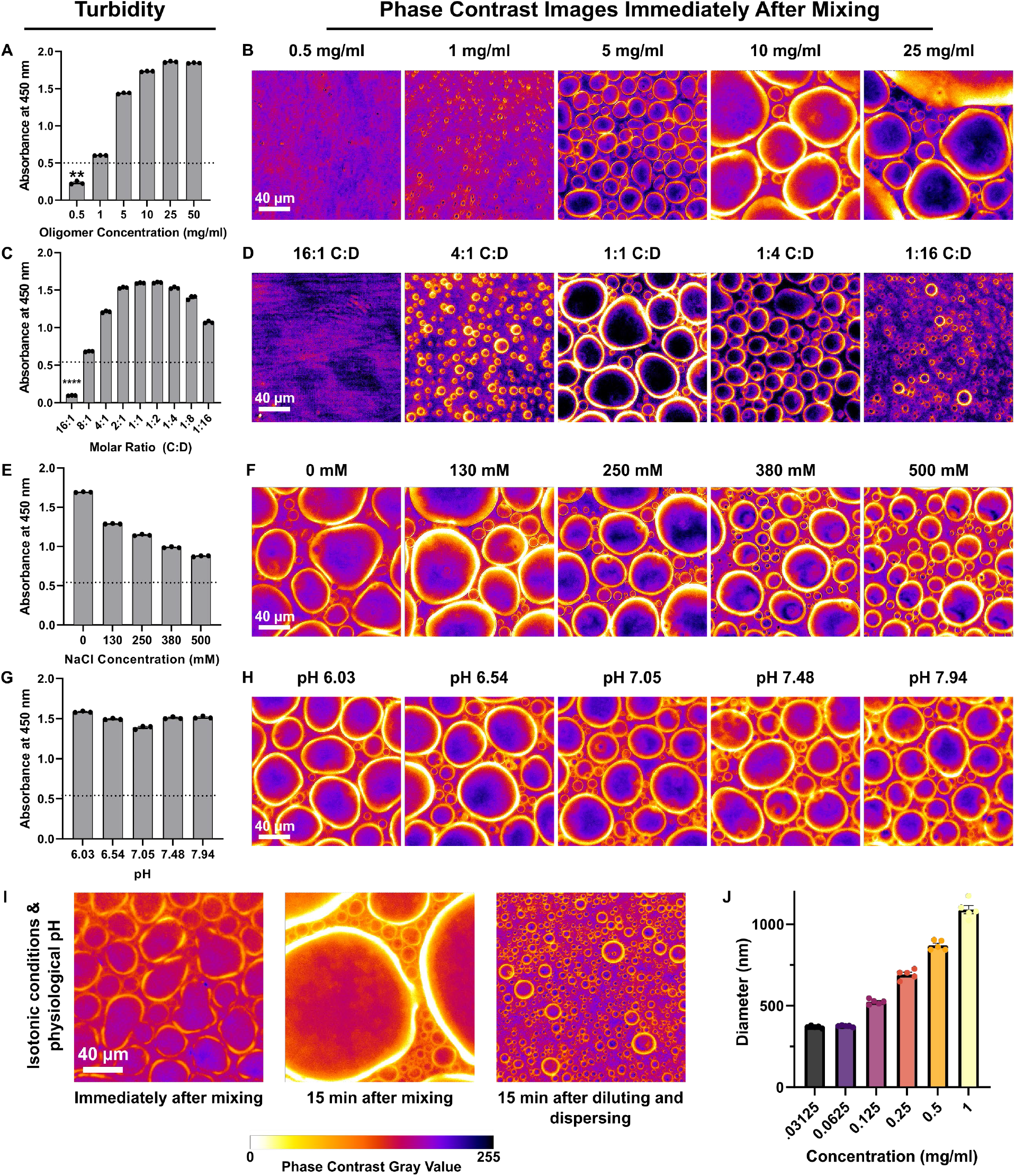
Effect of formulation factors on complex coacervate derivation with bC2.00 and bD2.00 oligomers. A) & B) Turbidity and corresponding phase contrast images for coacervates formed at oligomer concentrations ranging from 0.5-50 mg/ml in 20 mM phosphate buffer + 130 mM NaCl at pH 6 and 1:1 molar ratio. C) & D) Turbidity and corresponding phase contrast images for coacervates formed at C:D molar ratios ranging from 16:1 to 1:16 at 10 mg/ml in 20 mM phosphate buffer + 130 mM NaCl. C:D denotes molar ratio of bC2.00 to bD2.00. E) & F) Turbidity and corresponding phase contrast images of coacervates at NaCl concentrations ranging from 0-500 mM at 10 mg/ml 20 mM phosphate buffer at pH 6 and 1:1 molar ratio measured by absorbance at 450 nm. G) & H) Turbidity and corresponding phase contrast images of coacervates at different pH values ranging from pH 6-8 at 10 mg/ml in 20 mM phosphate buffer + 130 mM NaCl at 1:1 molar ratio. I) Phase contrast images of coacervates: upon formation at 25 mg/ml and 1:1 molar ratio in isotonic PBS at pH 7.4; after being left to coalesce for 15 minutes; and after being resuspended following a 2-fold dilution. J) Diameter of coacervates serially diluted from 1 mg/ml at 1:1 molar ratio in PBS as measured by DLS. Graph shows mean ± SEM with n=5. (A, C, E, G) Graphs show mean ± SEM with n=3 & Threshold Abs450 = 0.5 set at point of visible turbidity. (A, C) One sample t test below theoretical mean Abs450 = 0.5. ** P < 0.01, **** P < 0.0001.

These data show that trehalose-based oligomers form complex coacervates that coalesce into a bulk liquid phase upon mixing of oppositely charged oligomers and that coacervate formation occurs over a wide range of experimental conditions including under physiological salt and pH conditions. Although concentrated coacervates form bulk liquid phases quickly, this bulk phase can be readily dispersed into stable micro- and nano-droplets through simple mixing and dilution in biologically relevant buffers.

### Trehalose-based coacervates can be loaded with high concentrations of protein

To begin investigating protein incorporation into trehalose-based coacervates, we employed a readily available model protein, lysozyme (Lyz), to conduct initial studies. The size of Lyz (∼15 kDa) and its positive charge under physiologically buffered conditions (isoelectric point (pI) = 10-11) is similar to many neurotrophins, making it an appropriate surrogate for neurotrophins in our initial evaluations. In PBS, FITC conjugated Lyz (FITC-Lyz) was combined with bC2.00 and bD2.00 oligomer solutions at varied loading ratios (LR, oligomer mass/protein mass (mg/mg)) ranging from 25 to 100, with total protein concentration kept constant at 1 mg/ml. To enhance the protein loading potential of the positively charged FITC-Lyz, it was first combined with the soluble polyanion, bC2.00, prior to mixing with bD2.00. FITC-Lyz readily incorporated into coacervates irrespective of LR. Coacervate size at time of mixing was directly proportional to the LR, likely due to a paralleled increase in total oligomer concentration, such that coacervates with an average radius of 3.5 µm were detected with a LR of 25 while 20.3 µm radius coacervates were generated with a LR of 100 (Figure 4A,B). Coacervates formulated with protein at the lower LR of 25 persisted as stable micron-sized coacervates for several hours, whereas the higher LR samples coalesced into a bulk liquid phase quickly, within minutes. To quantify protein encapsulation efficiency, the coacervate liquid phase was isolated from the solution by centrifugation and the depletion of protein from the solution was characterized by fluorometry. Concentrating factor, which was defined as the fold-increase in detected concentration of the protein in the coacervate phase relative to its solution starting concentration, was computed by accounting for the total volume of coacervate formed and the total protein encapsulated. Increasing the oligomer solution concentration at a fixed LR (Figure 4C) or increasing the LR at a fixed protein solution concentration (Figure 4D) both served to increase the encapsulation efficiency of FITC-Lyz. At a fixed LR of 10, encapsulation efficiency as a function of oligomer solution concentration (1-100 mg/ml) followed a one phase association relationship with a plateau encapsulation of approximately 91% (Figure 4C). By contrast, the concentrating factor parameter showed a local maximum value of 14 at a much lower 20 mg/ml oligomer concentration. At a fixed protein solution concentration of 3.6 mg/ml, LR of 35 or more achieved a plateaued encapsulation efficiency of 93-99.6%. At this fixed protein solution concentration, the concentrating factor peaked at a LR of 10, resulting in formation of a coacervate phase loaded with FITC-Lyz at nearly 69 mg/ml (Figure 4D). To evaluate how robustly different proteins could be loaded into coacervates, we assessed encapsulation efficiencies and concentrating factors for two other model proteins, FITC-labeled Bovine Serum Albumin (FITC-BSA) (pI = 4.5-5.0) and FITC-labeled Human IgG (pI = 6.5-9.5) all prepared at the same protein solution concentration of 4 mg/ml. Coacervate encapsulation efficiencies for these proteins were dependent on overall net protein charge in the formulation buffer (pH = 7.4 for FITC-IgG, pH = 6.5 for FITC-BSA), such that the least charged protein under these conditions, FITC-IgG, had the lowest encapsulation efficiency at 16% whereas the most charged protein, Lyz, was nearly completely partitioned into the coacervate phase with an encapsulation efficiency of >99% (Figure 4E). FITC-IgG could be loaded into coacervates with higher encapsulation efficiency at the same LR by lowering the protein solution concentration to 2 mg/ml (Figure S11A). Encapsulation efficiencies of FITC-IgG could also be enhanced at higher protein concentrations by two-fold by increasing the LR from 50 to 200, indicating that FITC-IgG had not reached its plateau encapsulation value within the formulations tested (Figure S11B). For coacervate formulations displaying high protein encapsulation efficiency, we consistently noted a concentrating effect of at least 2.6-fold (Figure 4E, S11).

**Figure 4.**
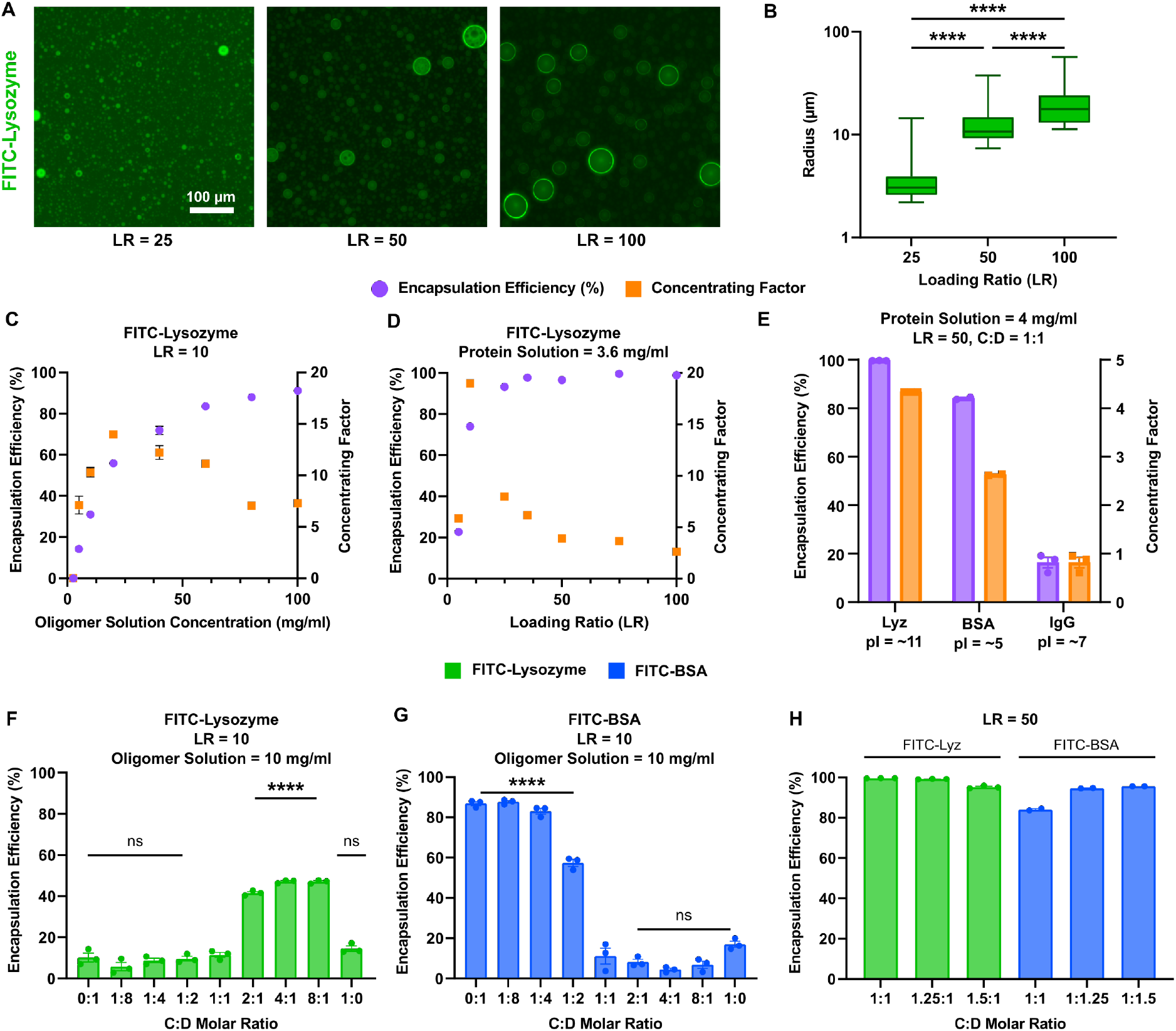
Protein-loading in trehalose-based coacervates. A) Fluorescence images of coacervates loaded with FITC-Lyz at loading ratios (LR) = 25, 50, 100 in PBS. FITC-Lyz concentration = 1 mg/ml. B) Size analysis of coacervates at LR = 25, 50, 100. Box & whiskers, min to max, midline denotes median. C) Encapsulation efficiency and concentrating factor of FITC-Lyz with increasing concentration of total oligomer in PBS. Constant LR = 10. D) Encapsulation efficiency and concentrating factor of FITC-Lyz with increasing LR in PBS. Constant protein concentration = 3.6 mg/ml. E) Encapsulation efficiency and concentrating factor of 3 different model protein solutions. FITC-Lyz, FITC-BSA, FITC-IgG coacervates formed with 4 mg/ml protein solutions at LR = 50, C:D = 1:1 in PBS. F) Encapsulation efficiency of FITC-Lyz at constant LR = 10 and oligomer concentration = 10 mg/ml at C:D ratios from 0:1 to 1:0. All groups compared to 1:1. G) Encapsulation efficiency of FITC-BSA at constant LR = 10 and oligomer concentration = 10 mg/ml at C:D ratios from 0:1 to 1:0. All groups compared to 1:1. H) Encapsulation efficiency of FITC-Lyz and FITC-BSA constant LR = 50 and oligomer concentration = 50 mg/ml with slight C:D molar excesses. (C-H) Graphs show mean ± SEM with n=2-3. All statistical tests are ordinary one-way ANOVA with Tukey’s multiple comparisons test. **** P < 0.0001.

We next explored how perturbing the molar ratio of polyanionic and polycationic oligomers altered protein encapsulation efficiency. To detect the relative contributions of molar ratio differences on protein encapsulation, we conducted studies using relatively dilute oligomer solutions (10 mg/ml) and a low LR of 10 which collectively promotes only a moderate encapsulation efficiency of ∼15-20% for both FITC-Lyz and FITC-BSA at stoichiometric equivalent ratios of the two oppositely charged bC2.00 and bD2.00 oligomers (Figure 4F,G). Formulations with excess polyanion, bC2.00, enhanced the relative encapsulation efficiency of cationic FITC-Lyz by approximately 4-fold, suggesting that charge imbalance can be an effective way to sequester more oppositely charged protein from the solution. However, for FITC-Lyz there was no significant increase in encapsulation detected beyond a molar ratio of imbalance 2:1 (C:D) (Figure 4F). Formulations prepared with excess polycationic bD2.00 oligomers did not result in a depletion of loaded FITC-Lyz compared to stoichiometric equivalency, suggesting an excess of like charges does not repel protein from the coacervate. For FITC-BSA, which has an acidic pI, preparing coacervates with an excess of polycationic oligomers also showed increased protein encapsulation (Figure 4G). However, in this instance, increasing the molar ratio imbalance up to 1:8 C:D did further increase the encapsulation of FITC-BSA compared to 1:2 C:D such that over 85% protein was partitioned into the coacervate phase. Interestingly, formulating FITC-BSA with just the polycationic bD2.00 oligomer alone was sufficient to induce coacervate formation, resulting in 87% FITC-BSA encapsulation. By contrast, bC2.00 alone with FITC-Lyz provoked formation of a minimal coacervate phase and a limited depletion of protein from the solution. These dissimilar outcomes can be readily attributed to differences in the immiscibility of the two oligomers in their neutralized states. Despite the significant contribution of oligomer charge imbalance in dilute solutions, at a higher LR of 50, the encapsulation efficiency was only minimally altered by perturbations in the molar ratio with all formulations having over 80% encapsulation efficiency at this LR and all but one formulation enabling 94-99.6% encapsulation (Figure 4H).

These data show that trehalose-based coacervates permit high encapsulation efficiency and concentration loading of a variety of different proteins including serum proteins, enzymes, and antibodies. Encapsulation efficiency and concentrating factor for proteins loaded into coacervates can be readily tuned on a protein-by-protein basis by optimizing formulation parameters such as the LR, oligomer and protein solution concentration, and molar imbalance of oppositely charged oligomers.

### Trehalose-based coacervates show temporally controlled bioactive protein release

We hypothesized that protein released from bulk coacervates could involve a combined contribution from several possible mechanisms including release via: (i) Fickian (first order) diffusion; (ii) oligomer ester degradation; (iii) bulk coacervate loss occurring through coacervate disassembly; or (iv) coacervate surface erosion leading to a steady surface blebbing of small coacervate droplets from the coacervate bulk. To determine the relative contributions of these potential mechanisms to overall protein release we started by investigating how coacervates degrade in the absence of loaded protein cargo. By selectively incorporating a small proportion of fluorescein labeling on bC2.00 using a fluorescein o-acrylate end-cap, we monitored mass of bulk coacervate material lost following daily replenishment of PBS supernatant. The rate of mass loss was maintained at essentially a constant rate (2-4 mg/day) across samples of increasing total mass such that 10 mg of coacervate sample was totally depleted within 4 days while 40 mg of sample took 10 days to completely resorb (Figure 5A). The approximate zero-order kinetics of coacervate mass loss suggested that surface erosion, rather than disassembly of charge interactions or ester hydrolysis, was the predominant mechanism of coacervate loss. Analysis of coacervate supernatants by DLS revealed the presence of nano- and micro-sized droplets in supernatants after each day of release, with >1 µm sized droplets on the first day of the incubation period followed by consistently sized droplets in the range of 250-650 nm detected thereafter (Figure 5B). Chemical analysis of the recovered coacervate supernatants by FTIR did not detect any emergent carboxylate groups in the C=O stretch at 1550-1610 cm^-1^ that would otherwise be present if there was substantial hydrolysis (Figure 5C). These data are consistent with the notion that coacervates are blebbing off from the coacervate bulk and diffusing into the supernatant as micro- and nano-sized droplets, suggesting that there is no disassembly of coacervate charge interactions and negligible oligomer hydrolysis up to at least 10 days of aqueous incubation.

**Figure 5.**
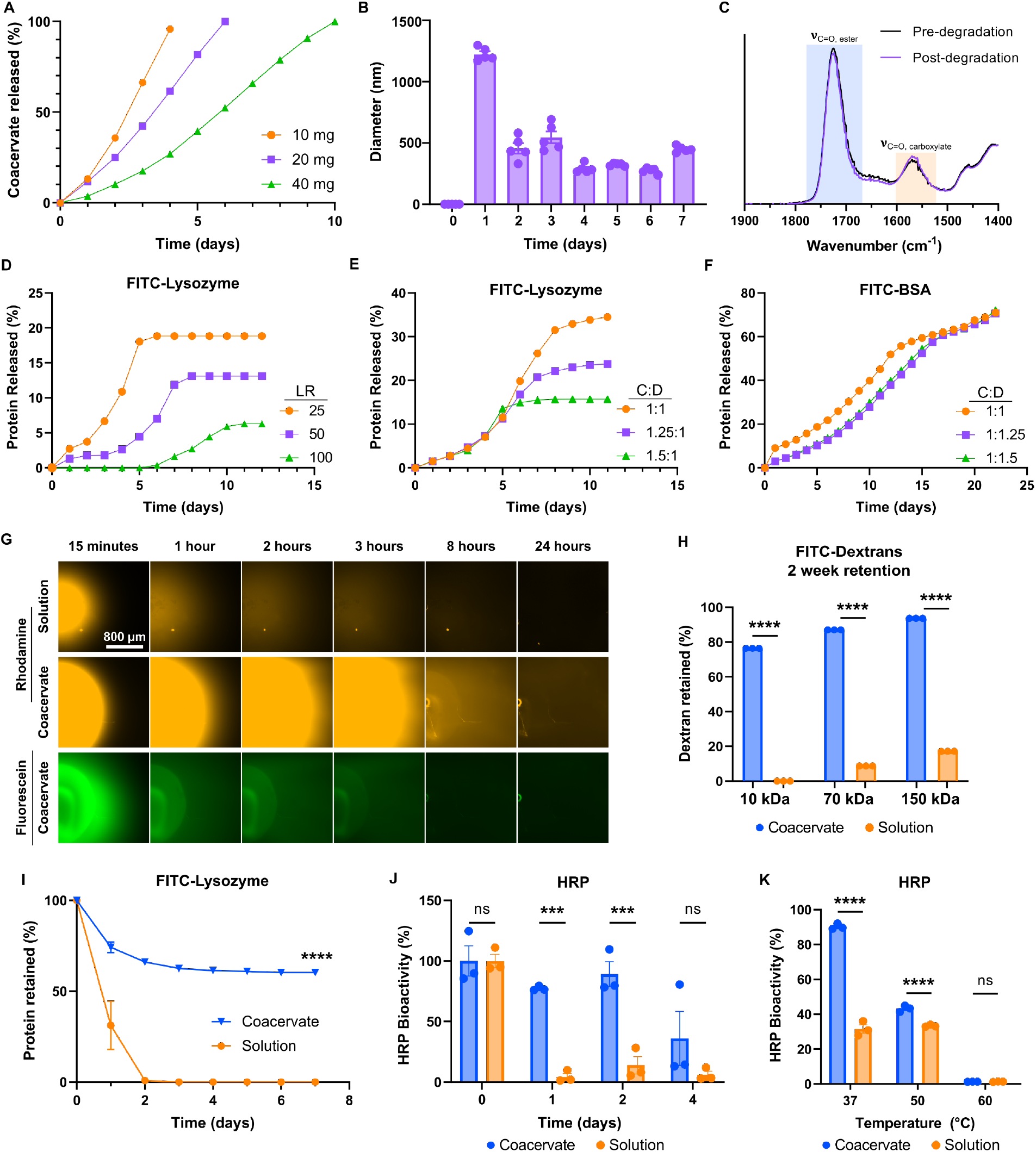
Temporally controlled bioactive protein release from coacervates. A) Percent mass released from bulk fluorescein-coacervates of different mass over 10 days. B) Diameter of particles detected in supernatant retrieved from incubated coacervates over 1 week at 37°C measured by DLS. Graph shows mean ± SEM with n=5 cycles. C) FTIR of coacervate material before and after release showing minimal change in the carboxylate stretch region. D) FITC-Lyz released from bulk coacervate at different LRs. LR 25 = 17.5 mg oligomer, LR 50 = 35 mg oligomer, LR 100 = 70 mg oligomer. E) FITC-Lyz released from bulk coacervate loaded with FITC-Lyz at different C:D ratios. LR = 50, oligomer mass = 40 mg. F) FITC-BSA released from bulk coacervate loaded with FITC-BSA at different C:D ratios. LR = 50, oligomer mass = 40 mg. G) Fluorescence images of rhodamine-dextran solution (top) and rhodamine-dextran loaded fluorescein-coacervates (middle, bottom) diffusing through 0.6 wt% agarose gel over 24 hours. H) Retention of FITC-dextrans in coacervate or solution at MW 10kDa, 70 kDa, 150 kDa in 0.6 wt% agarose gel. I) Retention profile of FITC-Lyz in coacervate or solution in 0.6 wt% agarose gel. J) Horseradish peroxidase (HRP) bioactivity measured over 4 days at 37°C in coacervate or solution. Measured with TMB substrate and absorbance at 450 nm. K) HRP bioactivity measured after 1-hour incubations in coacervate or solution at different temperatures. Measured with TMB substrate and absorbance at 450 nm. (A, D-F, H-K) Graphs shows mean ± SEM with n=3. (H-K) 2way ANOVA with Sidak’s multiple comparisons test. Not significant (ns), *** P < 0.001, **** P < 0.0001.

To evaluate protein release from coacervates we first incorporated FITC-Lyz into coacervates at different LRs with a fixed protein mass of 0.7 mg and oligomer C:D ratio of 1:1, such that there was essentially equivalent protein encapsulation across the different samples (93-99%) and evaluated protein release from bulk coacervates into PBS that was replaced daily. There was reduced initial burst and decreased total FITC-Lyz released from coacervates with increasing LR (Figure 5D). Surprisingly, the total amount of FITC-Lyz recovered from surface eroding coacervates was low across all samples. Approximately 20% of total FITC-Lyz was recovered from coacervates with a LR of 25 while we detected essentially no released FITC-Lyz for up to 5 days in coacervates with a LR of 100 and in this formulation only around 5% of total loaded FITC-Lyz was recovered (Figure 5D). For a fixed LR, mixing FITC-Lyz with the polyanion prior to coacervate formulation with the polycation provided a small but significant increase in encapsulation efficiency when compared to protein loading after coacervate formation but did not alter the overall percentage of protein recovered during *in vitro* release (Figure S12). Associated with the decreasing recovery of FITC-Lyz in coacervates with higher LR was the presence of water-insoluble precipitates in the supernatant that only emerged after the first 5 days of incubation when most of the coacervate bulk had eroded away (Figure S13). There was negligible additional release of FITC-Lyz from the solid precipitates for the 12 days of testing indicating that these precipitates were likely particularly stable solid protein-oligomer complexes (Figure 5D). From these results we inferred the coexistence of both liquid coacervate and solid precipitate phases when trehalose-based oligomers are formulated with Lyz, with the relative abundance of protein within the precipitate phase increasing with a greater excess of oligomer. Prior studies have demonstrated a preference towards solid precipitate formation when Lyz is combined with polyanions due to its particularly high isoelectric point and density of cationic charge at physiologic pH [51]. To confirm that the formation of solid precipitates with Lyz is due directly to polyanion interactions, we examined formulations with an increasing excess of polyanion species at a constant LR of 50 (Figure 5E). Although the extent of protein release during the first 5 days of coacervate erosion was essentially unchanged, a higher ratio of polyanion oligomers resulted in greater retention of FITC-Lyz in the precipitate phase, consistent with literature [51] (Figure 5E). We also detected co-existence of coacervate and precipitate phases in FITC-BSA formulations with the same LR. However, there was minimal effect of excess polycation on FITC-BSA recovery and there was continuous release of FITC-BSA from the precipitate phase during a 24-day testing period (Figure 5F). These data are consistent with FITC-BSA-oligomer solid precipitates being less stable compared to those formed with FITC-Lyz, owing to the lower density of charge on BSA at physiological pH. It was also notable that fluorescein-labeled coacervates formulated with either unlabeled Lyz or unlabeled BSA showed faster coacervate surface erosion compared to empty fluorescein-labeled coacervates, suggesting that the solid precipitate formation disrupts the stability of coalesced coacervates (Figure S14). Overall, these protein release data suggest that trehalose-based oligomers effectively prolong protein release via combined action of coacervate surface erosion followed by solid precipitate dissolution and that the net release profile is both protein and formulation dependent.

Intraparenchymal infusions of macromolecules or nanoparticles into the CNS results in extensive transport of these entities along perivascular spaces of blood vessels throughout neural tissue via convection-enhanced delivery (CED) that may ultimately hamper desired local therapeutic effects [52, 53]. To assess *in vitro* whether coacervates might mitigate the extent of CED and retain molecules within more focal tissue volumes, we evaluated transport of different molecular weight dextrans and proteins loaded into coacervates following injection into 0.6 wt% agarose gel phantom brains (Figure S15), which serve as effective *in vitro* systems for modeling CED[54]. We loaded non-ionic rhodamine labeled dextran (10 kDa) into fluorescein-labeled coacervates so that we could track both coacervate and dextran transport individually through the phantom (Figure 5G). Non-charged dextrans have minimal electrostatic interactions with oligomers thus mitigating the formation of a precipitate phase. There was notably slower transport of dextran loaded coacervates through agarose phantoms compared to the dextran solution over the course of 24 hours, with dextran retention correlating with coacervate retention in the agarose (Figure 5G). The co-dispersion of coacervates and dextrans illustrates that loaded cargo does not readily diffuse out of coacervates, but that coacervates remain as a protective carrier for cargo throughout dispersion through the agarose. Coacervate loading of non-charged dextrans was dependent primarily on molecular weight, with encapsulation efficiency increasing from 36% for 10 kDa to 60% for 70 kDa when loaded at 2 mg/ml with a LR of 50 (Figure S16). There was no difference in the encapsulation efficiency between 70 and 150 kDa suggesting that coacervates readily accommodate larger molecular weight cargo. Coacervates enhanced the retention of different molecular weight dextrans in agarose phantoms in a size dependent manner over a 2-week period *in vitro* with 76% of 10 kDa and 94% of 150 kDa dextran retained in the agarose by formulating with coacervates. By contrast, no 10 kDa dextran and only 17% of 150 kDa dextran was retained in agarose phantoms at 2 weeks when injected as a solution (Figure 5H, S16). Monitoring recovered FITC-Lyz injected into agarose phantoms also showed enhanced local retention using coacervates with greater than 60% of protein preserved in the agarose phantom after 1 week whereas FITC-Lyz solution was completely removed from the agarose after just 2 days (Figure 5I). Collectively, these data show that coacervates may be an effective way of minimizing CED effects to retain high local concentrations of molecular cargo, including proteins and non-charged polysaccharides, in narrow tissue zones *in vivo*.

To determine the capacity of trehalose-based coacervates to maintain the bioactivity of loaded proteins, we loaded horseradish peroxidase (HRP) into coacervates and monitored time-, temperature-, and lyophilization-dependent effects on enzymatic activity compared to HRP solutions. HRP is a suitable model protein for evaluating the effects of trehalose-based coacervates on protein stability and function since any conformational or structural perturbations to HRP cause loss of enzymatic activity, and dilute solutions of HRP become inactive within 24 hours[27]. Formulating HRP into coacervates did not alter the bioactivity of the enzyme, and coacervates preserved HRP enzymatic activity for up to 4 days longer at 37°C *in vitro* compared to the HRP solution (Figure 5J). Formulating HRP into coacervates provided moderate protection against denaturing thermal stressors, such that higher preserved enzymatic activity was maintained at 50°C, although the stabilizing effect was lost at temperatures at or above 60°C (Figure 5K). Trehalose-based coacervates also enabled lossless recovery of bioactive HRP following lyophilization, which was an improvement on the recovery from protein solution only formulations (Figure S17). These data show that coacervates are effective carriers for maintaining long-term functional bioactivity of proteins during controlled release.

### Trehalose-based coacervates are cytocompatible and have formulation dependent interactions with neural cells *in vitro*

To test the cytocompatibility of trehalose-based coacervates with neural cells, we exposed proliferating mouse neural progenitor cells (NPC) and quiescent mouse astrocyte cultures to coacervates *in vitro* (Figure 6, S18, S19). Formulated neutral coacervates, as well as their individual constituents, bC2.00 and bD2.00, were tolerated well by both NPC (Figure 6A, S18) and quiescent astrocytes (Figure S19) at all concentrations tested up to 1 mg/ml with cell viability being not significantly different to that of untreated controls. In a separate assay, the number of viable NPC detected after 24 hours was directly proportional to the treated coacervate concentration with the highest 1 mg/ml concentration promoting a 30% increase in the number of detected NPC compared to the lower 0.03125 mg/ml concentration, with a significant linear correlation of cell viability to log_2_(coacervate concentration) (Figure S18). NPC proliferation rates are exquisitely sensitive to the concentrations of EGF and FGF growth factors used in the cell culture media[55]. We surmised then that the above results could be due to coacervates sequestering EGF/FGF from the media resulting in an amplified proliferation effect due to coacervate mediated presentation of higher local concentrations of growth factors at cell membranes. To test this hypothesis, we next formulated coacervates at high concentration in PBS and then diluted them into growth factor-supplemented cell culture media containing the standard 100 ng/ml concentration of EGF and FGF growth factors (100 EF media) to obtain final coacervate concentrations ranging from 2 to 10 mg/ml. Upon coacervate mixing, the solution was immediately centrifuged to pellet the coacervates and enable the coacervate-exposed media to be decanted and recovered. NPC grown in coacervate-exposed media showed a coacervate concentration-dependent reduction in total number of NPC after 4 days of incubation, consistent with these media having reduced concentrations of growth factors, due to sequestration of growth factors from the media by coacervates (Figure 6B). We then took the coacervate bulk that had been exposed to 100 EF supplemented media, mixed it into un-supplemented (0 EF) media, and applied it to NPC cultures. Incorporating these coacervates into the media promoted a modest but significant increase in total cell number after 4 days of culture compared to cells grown without growth factors, which indicates that coacervates effectively presented bioactive EGF/FGF that they had sequestered from the 100 EF media to NPC (Figure 6C). These data show that coacervates can serve to sequester growth factors and present them locally at cells to augment their effects.

**Figure 6.**
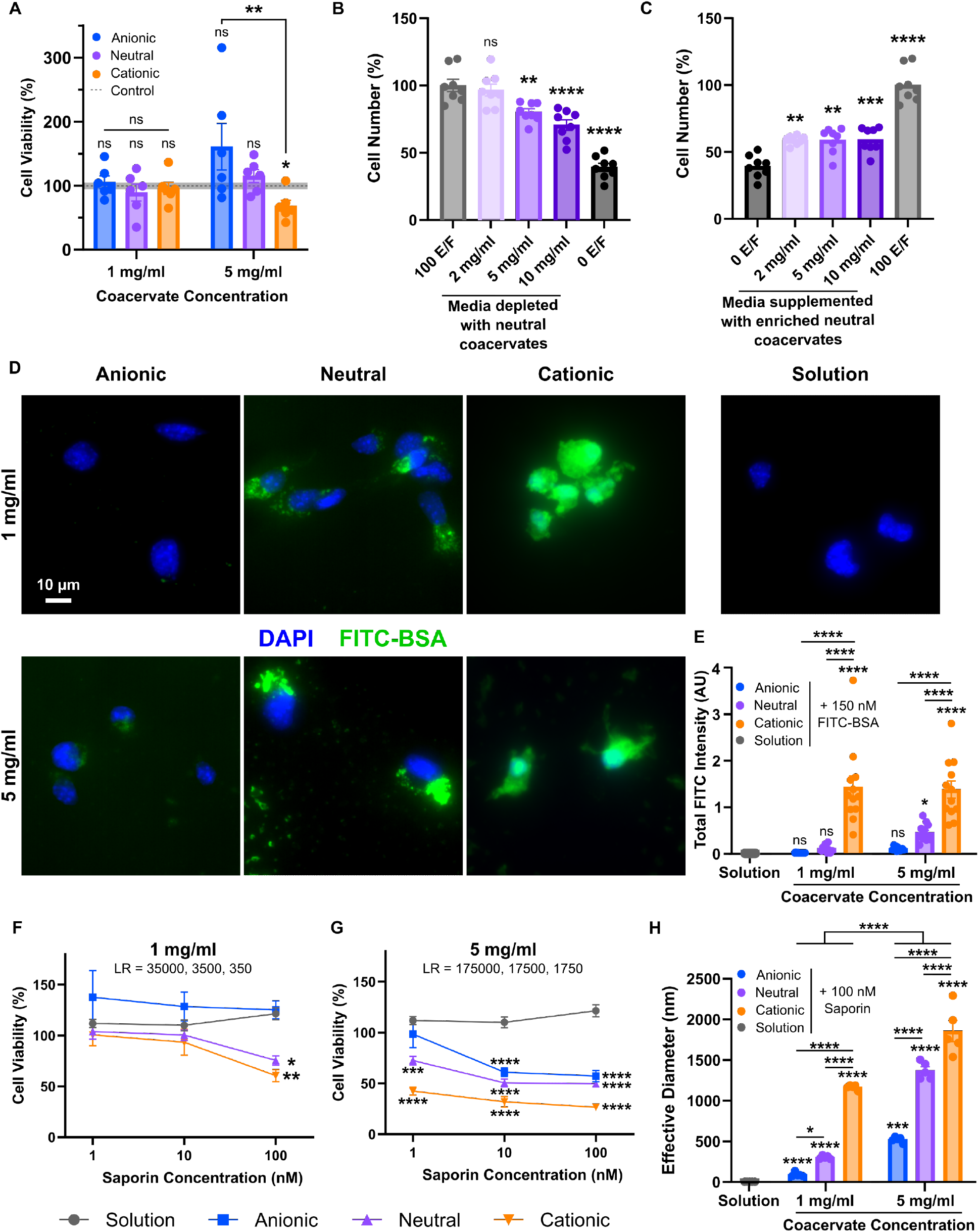
Coacervates are cytocompatible and can facilitate formulation-dependent intracellular or extracellular protein delivery *in vitro*. A) Viability of neural progenitor cells (NPC) with coacervates at different formulations (Anionic = 4:1, Neutral = 1:1, Cationic = 1:16 (C:D)) at 1 mg/ml and 5 mg/ml for 72 hours; measured by Calcein AM assay. Dotted line and shading represents mean ± SEM of control. B) NPC number for cells treated with neural expansion media either fully supplemented with 100 ng/ml EGF/FGF (100 E/F), with no added EGF/FGF (0 E/F), or with media partially depleted of supplements using neutral coacervates for 4 days; measured by Calcein AM assay. C) NPC number of cells treated with neural expansion media either fully supplemented with 100 ng/ml EGF/FGF (100 E/F), with no added EGF/FGF (0 E/F), or with media partially supplemented using neutral coacervates that depleted fully-supplemented media for 4 days; measured by Calcein AM assay. D) Images of NPC treated with FITC-BSA coacervates or solution at different formulations (Anionic = 4:1, Neutral = 1:1, Cationic = 1:16 (C:D)) at 1 mg/ml (LR = 100) and 5 mg/ml (LR = 500) for 72 hours. FITC-BSA solution was at 10 µg/ml in all formulations. Stained with DAPI. E) Quantification of NPC images of FITC-BSA intensity normalized to DAPI cell count. F) Viability of NPC treated with coacervates loaded with saporin at different formulations (Anionic = 4:1, Neutral = 1:1, Cationic = 1:16 (C:D)) at 1 mg/ml for 72 hours; measured by Calcein AM assay. G) Viability of NPC treated with coacervates loaded with saporin at different formulations (Anionic = 4:1, Neutral = 1:1, Cationic = 1:16 (C:D)) at 5 mg/ml for 72 hours; measured by Calcein AM assay. H) DLS sizing data of coacervates in media loaded with 100 nM saporin at different formulations (Anionic = 4:1, Neutral = 1:1, Cationic = 1:16 (C:D)) at 1 mg/ml and 5 mg/ml. (B, C) Graphs show mean ± SEM with n=8 with ordinary one-way ANOVA with Tukey’s multiple comparisons test. (B) Comparisons made to 100 E/F. (C) Comparisons made to 0 E/F. (A, E, H) Graphs show mean ± SEM with n=5-7 with ordinary one-way ANOVA with Tukey’s multiple comparisons test within concentration groups to control or solution, and 2way ANOVA with Tukey’s multiple comparisons test across concentration groups. Symbols directly above bars represent comparisons to control. (F, G) Graphs show mean ± SEM with n=6-7 with 2way ANOVA with Tukey’s multiple comparisons test. Symbols indicate comparisons made to solution at equivalent saporin concentration. Not significant (ns), * P < 0.05, ** P < 0.01, *** P < 0.001, **** P < 0.0001.

Next, we wanted to understand how coacervates interact with cultured cells. Using fluorescein-labeled coacervates to visualize coacervate-cell interactions, we noted that many of the micro-sized coacervates that form at the 1 mg/ml concentration seemed to readily adhere to both neural cell types after 24 hours of incubation (Figure S18, S19). Adhesion of coacervates to cells was a surprising observation since we anticipated that a net neutral, trehalose-based surface would be non-fouling [56]. To further investigate this, we probed whether a surface layer of bulk coacervates promoted cellular attachment by coating non-treated suspension polystyrene plates with 2 mg/ml of coacervates or constituent oligomers, culturing cells directly on top of the thin residual material layer and evaluating the proportion of adhered cells resistant to wash-off after 24 hours. For NPC, the coacervate layer or adsorbed oligomer surface significantly increased the proportion of attached cells by almost four-fold compared to polystyrene controls with over 90% of cells remaining adhered (Figure S18D). Cultured astrocytes showed a similar capacity to adhere to coacervates and individual oligomers, although polyanionic oligomers showed slightly reduced cell attachment (Figure S19B).

Peptide- and pyrene-based coacervate systems have effectively delivered proteins intracellularly via an endocytic pathway [57–59], so we next evaluated the intracellular protein delivery capacity of trehalose-based coacervates. A net cationic charge is a conserved feature of intracellular delivery carriers, such as cell penetrating peptides and engineered lipid or polymer-based vehicles, that enables them to engage negatively charged cell membranes and facilitate endosomal escape [60–62]. To test whether surface charge on coacervates may alter their intracellular delivery capacity, we prepared 3 different formulations: (i) Neutral coacervates where C and D functionalized oligomers were added with stoichiometric equivalency; (ii) Anionic coacervates generated through an oligomer C:D ratio of 4:1, and (iii) Cationic coacervates generated through an oligomer C:D ratio of 1:16. These specific C:D ratios were selected because they provided the greatest net excess of anionic or cationic charge while still being able to generate microscopically detectable coacervates (Figure 3D). Cationic coacervates had detectable positive charge, whereas neutral coacervates did not have any measurable cationic charge by TNS assay (Figure S20). We prepared the three differentially charged coacervates at two concentrations for cell experiments: (i) 1 mg/ml to replicate the aforementioned cell viability experiments, and (ii) 5 mg/ml to provide a higher concentration of coacervates that may alter cell uptake. All coacervates formulated without protein cargo at 1 mg/ml evoked no detectable cytotoxicity, with cell viability no different to the untreated control group after 3 days of incubation. When NPC were exposed to coacervates at 5 mg/ml, there was a small but significant loss in cell viability (∼30%) for only the cationic coacervate treated group (Figure 6A).

To evaluate intracellular delivery outcomes for the coacervate formulations with different surface charges, we loaded coacervates with FITC-BSA and applied them to cultured NPC in serum free media for 3 days at the two different concentrations (1 mg/ml and 5 mg/ml). Without a suitable carrier, BSA is not extensively transported into the cell cytoplasm [63] and, since it is well tolerated by *in vitro* cultures, it serves as a suitable model protein to evaluate intracellular delivery. All coacervate formulations showed high FITC-BSA encapsulation efficiencies (>70% for all formulations) even when maintaining net anionic or cationic charge, which was attributed to the use of the relatively high LRs of 100 for 1 mg/ml and 500 for 5 mg/ml. After three days of incubation, cationic coacervates promoted significant intracellular delivery of FITC-BSA resulting in dissemination of protein throughout the cell cytoplasm for both concentrations (Figure 6D,E). We also detected increased intracellular accumulation of FITC-BSA in cells using neutral coacervates, but this was only significantly increased above background levels for the 5 mg/ml concentration. There was no detectable delivery of FITC-BSA to the cell cytoplasm when loaded in anionic coacervates for both concentrations (Figure 6D,E).

To test for functional intracellular protein delivery by the differently charged coacervate formulations, we next loaded the coacervates with saporin. Saporin is a ribosome inactivating protein that causes cell cytotoxicity when introduced in its intact form into the cell cytoplasm However, saporin has limited cell membrane permeability, so without a carrier to effectively transport saporin into the cell cytoplasm, it has no detectable cytotoxicity within a defined concentration range [57, 64]. Saporin has an isoelectric point of ∼9.5, and thus has a slight net positive charge at physiologic pH. We encapsulated saporin into anionic, neutral, and cationic coacervates at 3 different solution concentrations (1, 10, and 100 nM) and incubated the saporin-loaded coacervates with NPC at 1 mg/ml and 5 mg/ml for 3 days prior to evaluation of cell viability. Given the relative low amount of saporin needed to induced cytotoxicity, the LRs for saporin at the 1 mg/ml coacervate concentration were 35000, 3500, and 350 for the 1, 10 and, 100 nM saporin concentrations respectively, while 5 mg/ml coacervates had LRs that were 5 times higher. These very high loading ratios ensure near total encapsulation of saporin into the coacervate phase for all formulations. Applying the 1 mg/ml coacervate concentrations to NPC for 3 days resulted in no detectable cytotoxicity for any coacervate type compared to the saporin solution control at 1 and 10 nM saporin loading, but at 100 nM both the neutral and cationic formulations caused measurable and significant cell death (Figure 6F). By contrast the anionic coacervates showed no cytotoxicity at the 100 nM saporin concentration for the same coacervate concentration. At 5 mg/ml coacervate concentrations, 1 nM saporin loaded into cationic coacervates had a pronounced cytotoxic effect (∼60% cell loss) which was greater than what was observed for the unloaded cationic coacervates at this same coacervate concentration (Figure 6A,G). Saporin loaded at 1 nM in neutral coacervates had moderate cytotoxicity (∼30% cell loss) at 5 mg/ml whereas anionic coacervates showed no loss of cell viability under the same conditions. As the concentration of saporin increased in the 5 mg/ml coacervate concentrations, all formulations, irrespective of charge imbalance, showed some measurable induced cytotoxicity suggestive of functional saporin intracellular delivery (Figure 6G). However, for all tested saporin concentrations, the cationic coacervates had a comparatively greater cytotoxic effect suggesting that more effective protein delivery was achieved by cationic coacervates.

Collectively, these data show that intracellular protein delivery by coacervates is concentration, incubation time, and surface charge dependent (Figure 6, S18, S19). Ultimately, coacervates formulated with excess cationic charge promoted the most effective intracellular protein delivery and achieved widespread dispersal of protein cargo throughout the cell cytoplasm. However, these results do not rule out potential contributions of coacervate size to intracellular delivery, since formulations prepared with excess cationic charge tended to generate larger coacervate droplets (1-2 μm) compared to the equivalent formulations with excess anionic characteristics (<1 μm) (Figure 6H). However, the size dependent effects, if they exist, are likely to be within a narrow range, since larger coacervate droplets and bulk coacervate substrates are not readily internalized by cells.

### Trehalose-based coacervates permit local bioactive protein delivery to the mouse striatum

The favorable cytocompatibility and bioactive protein release properties of trehalose-based coacervates motivated us to test coacervate-based delivery *in vivo* in the mouse CNS. Rheological evaluations revealed that bulk coacervates are viscoelastic but readily thin upon application of high strain (Figure S21A), indicating that we could easily draw up coacervates into narrow glass micropipettes for infusion into the mouse brain (Figure S21B). We injected coacervates into the caudate putamen region of mouse striatum, which is a readily accessible brain region that contains diverse neural parenchymal elements. To evaluate delivery outcomes, we used a standardized CNS FBR immunohistochemistry (IHC)-based framework that we have validated extensively [24, 32]. As part of this assay, we loaded biotinylated dextran amine (10 kDa) (BDA), a non-bioactive model macromolecule, into coacervates to track coacervate-dependent CNS molecular delivery outcomes by IHC. We evaluated coacervate-dependent delivery of BDA in two forms: (i) as coacervate bulk (100% v/v phase-separated material), or (ii) as diluted coacervate droplets suspended within a BDA containing solution at 10% v/v phase-separated material. Both bulk and suspended coacervates were formulated at a LR of 100 and 1:1 ratio bC2.00 to bD2.00. To compare coacervate delivery, we employed BDA-only solutions as well as a BDA loaded methyl cellulose (MC)-based hydrogels as benchmarking controls [24, 32]. Detailed quantitative IHC analysis on brain tissue sections recovered from cohorts of treated mice revealed notable differences in neural tissue biodistribution between the four different treatments at 7 days after injection (Figure 7A-F). Infused bulk coacervates could be readily detected at 24 hours after injection by BDA staining as micro-sized spherical droplets, with coacervates dispersing throughout a narrow region of neural parenchyma in the striatum that was no more than approximately 380 µm wide with individual coacervates intermingling within beds of NeuN-positive neurons (Figure S22A,C). Bulk coacervate injections caused minimal disruption of neural tissue that is otherwise observed for bulk hydrogel injections into striatum [24, 32] (Figure 7B,G,I), with essentially no DAPI-negative regions detected for coacervate-treated brains at 24 hours after injection (Figure S22B). This result is characteristic of tissue dispersal of coacervates rather than deposition of large volumes of material that would otherwise cause local tissue displacement around the material. Evaluation of infused BDA solution at 7 days after injection revealed extensive uptake of BDA into neurons and axons throughout the striatum and the cortex as well as transport of BDA along large blood vessels resulting in extensive accumulation of BDA in perivascular spaces and at the meninges (Figure 7A,C,D, S23). By contrast, bulk coacervate-based delivery resulted in reduced tissue penetration of BDA such that the 50^th^ percentile of detected BDA was constrained to within a radial distance of less than 75 µm from the injection epicenter which was 26% of the BDA tissue penetration observed for both the BDA solution and MC hydrogels (Figure 7D). Bulk coacervate treated mice showed no detectable accumulation of BDA along perivascular spaces or at the meninges (Figure S23). Suspended coacervates (10% v/v) increased the striatal penetration of BDA compared to bulk coacervates such that it was no different to BDA solutions, although suspended coacervates had notably less cortical and meningeal delivery compared to BDA solutions (Figure 7C-F, S23). Constrained BDA delivery by bulk coacervates was correlated with significantly less total BDA accumulation in neural tissue compared to BDA solution, but it was equivalent to that detected for MC hydrogels (Figure 7E,F). This decreased BDA accumulation is consistent with a reduction in BDA diffusion and cell uptake throughout the neuropil occurring acutely, which is typical of delayed release systems [24]. BDA released locally from coacervate formulations was taken up by NeuN-positive neurons but was not detected in Iba-1 positive microglia (Figure S24).

**Figure 7.**
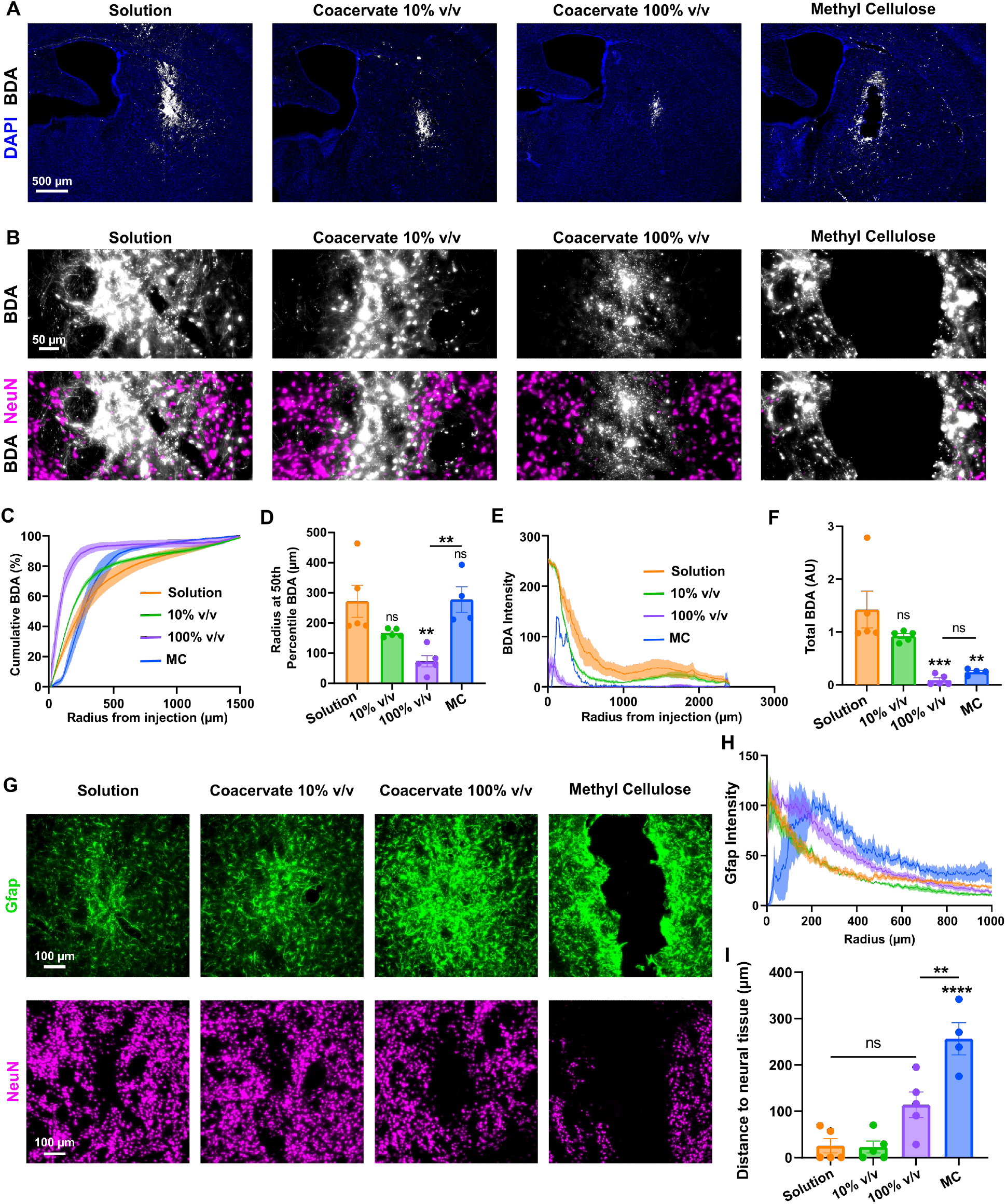
BDA-loaded coacervates injected into healthy mouse striatum. A) Survey images showing biotinylated dextran amine (BDA) dispersion through brain with BDA-loaded solution, suspended (10% v/v) and bulk (100% v/v) coacervate, and methyl cellulose (MC) gel injections into the striatum. B) Detailed images showing BDA deposit at the injection site and proximity to nearby neurons with BDA-loaded solution, 10% v/v and 100% v/v coacervate, and methyl cellulose gel injections. C) Cumulative percent BDA delivered to neural tissue over 1.5 mm radial distance from injection site. D) Radial distance from injection at which 50% of total cumulative BDA is dispersed. E) Radial intensity profile for BDA indicating the radial dispersion of BDA from striatal injection site. F) Total BDA in ipsilateral striatum and cortex. G) Images of reactive astrocyte response (Gfap) and neural cell (NeuN) displacement at injection site. H) Radial intensity profile of Gfap-positive astrocytes evaluated from the striatal injection epicenter. I) Radial distance from injection epicenter to the start of NeuN-positive neural tissue. All graphs show mean ± SEM with n=4-5. Ordinary one-way ANOVA with Tukey’s multiple comparisons test. Not significant (ns), ** P < 0.01, *** P < 0.001, **** P < 0.0001.

Locally injected biomaterials that evoke severe FBRs, such as cationic sulfonium or chitosan gels [24], promote the formation of non-neural cell compartments as a result of localized recruitment and persistence of blood-borne innate immune cells at the biomaterial-tissue interface [24]. Neither suspended or bulk coacervate treatments induced formation of any non-neural compartment and there was minimal peripherally-derived immune cell recruitment, such that there was no difference in the overall numbers of Ly6b2-positive neutrophils and Cd13-positive myeloid lineage cells (e.g. macrophages, monocytes) for coacervates compared to the solution control at 7 days post-injection (Figure S25A-C). Bulk coacervates evoked an elevated and more diffuse astrocyte reactivity response, measured by increased Gfap expression, compared to either suspended coacervates or solution controls (Figure 7G,H, S26). However, despite the increased astrocyte reactivity response caused by bulk coacervate injections, there was no difference in the overall magnitude of this response compared to MC hydrogels (Figure S27). Notably, coacervates did not provoke the formation of new glial limitans-like borders or cause any significant depletion of NeuN-positive neural tissue which is otherwise associated with MC hydrogel treatment (Figure 7G-I). Bulk coacervates moderately increased expression of vimentin in local astrocytes at the injection site compared to solution-only controls consistent with increased astrocyte reactivity (Figure S28A,B). We did not detect any Ki67-positive astrocytes in coacervate-treated or solution groups at 7 days after injection but there were notably fewer Sox9-positive astrocytes at the injection epicenter of coacervate treated mice (Figure S28C). Bulk coacervate treatment promoted a small but significant localized elevation in Iba-1-positive cells at the injection epicenter which was correlated with a small decrease in the number of P2ry12-positive microglia compared to solution controls (Figure S25D-F). Downregulation of P2ry12 and upregulation of Iba-1 suggests that bulk coacervates cause a small but detectable local activation of microglia at the injection site.

To assess bioactive protein delivery to the CNS using coacervates, we applied a neurotrophin delivery model that we have used previously in the evaluation of hydrogel-based delivery systems [24]. This model leverages the established biological effect of the neurotrophin, NGF, on basal forebrain cholinergic neurons. These neurons are sensitive to extracellular NGF concentrations such that they atrophy when deprived of NGF and hypertrophy when exposed to exogenous NGF. Measurable changes in striatal cholinergic neurons following injections of soluble NGF or delivery of NGF from biomaterials can be readily detected by choline acetyltransferase (ChAT) IHC staining and subsequent analysis of increased number and size of these cells as well as increased density of cholinergic axon networks [24]. Recombinant human neurotrophin, NGF (pI = 9.5) in PBS, was loaded into coacervates prepared with a modest excess of polyanion bC2.00 (C:D = 1.25:1) at a LR of 100 that resulted in an encapsulation efficiency of 60% with a concentrating factor of 6 (Figure 8A). NGF loaded coacervates were delivered as a bulk 100% v/v formulation via stereotactic injection into the mouse striatum (Figure 8B). Brains were evaluated at 7 days after injection by IHC and showed no differences in the total numbers of NeuN-positive neurons within the ipsilateral striatum across the NGF coacervate, NGF solution, and saline control groups (Figure S29), suggesting no significant neural damage was induced by coacervate injection. Coacervates effectively delivered bioactive NGF, causing a significant 78% increase in ChAT intensity and approximately 35% increase in neuron size in the ipsilateral striatum compared to the cells on the contralateral side, whereas saline treated controls showed no changes in these parameters (Figure 8C-E, S29). The extent of the NGF-induced changes to ChAT-positive neurons in the ipsilateral striatum promoted by coacervate-based NGF delivery were not significantly different to that evoked by NGF solution (Figure 8D,E). However, coacervate-based delivery resulted in a more focal treatment effect, such that increases in ChAT intensity were isolated to a narrower 700 µm radial distance from the injection epicenter for coacervates whereas the NGF solution increased ChAT intensity throughout the entire ipsilateral striatum (Figure 8F) as well as in the contralateral striatum (Figure 8G). Cumulative representations of ChAT intensity in the ipsilateral striatum reveal that coacervates and solution have a similar effect close to the injection but that coacervates begin to have a reduced effect around 500 µm from the injection epicenter (Figure 8H). Cumulative ChAT intensity in the contralateral striatum further illustrates the extensive distribution of NGF solution to the mouse brain whereas coacervates minimize that effect (Figure 8I). These data show that trehalose-based coacervates are effective vehicles for precisely delivering bioactive proteins to the neuropil.

**Figure 8.**
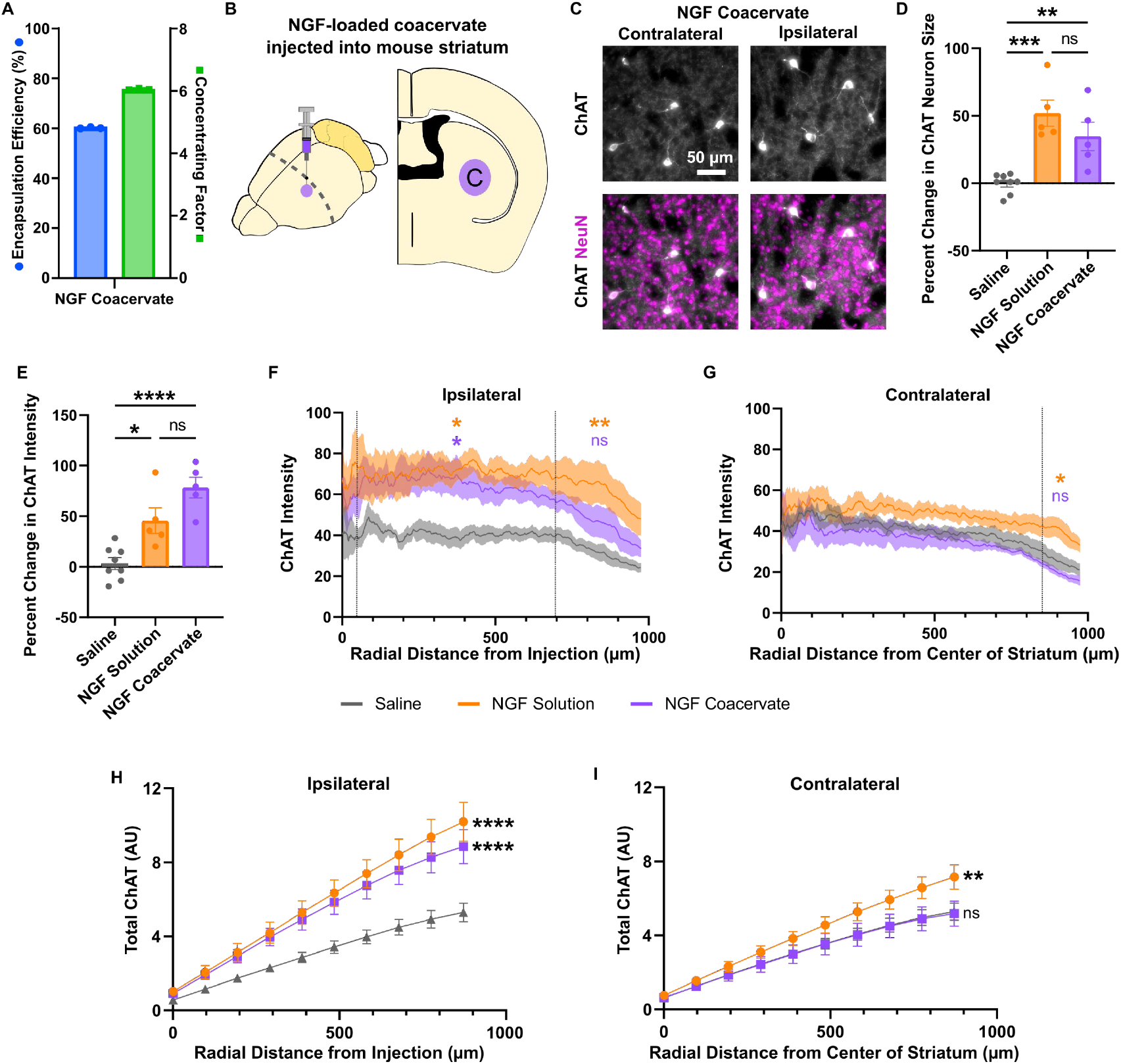
Local delivery of NGF to ChAT-positive neurons in mouse striatum using coacervates. A) Encapsulation efficiency and concentrating factor of NGF coacervates formed with 1 mg/ml NGF solution at LR = 100, C:D = 1.25:1 in PBS; measured by ELISA. B) Schematic of NGF coacervate injection into healthy mouse striatum. C) Detailed images of choline acetyltransferase (ChAT)-positive neurons among neural tissue (NeuN) with nerve growth factor (NGF) coacervate delivery in ipsilateral and contralateral striatum. D) Quantification of percent change in size of ChAT-positive neurons from ipsilateral to contralateral striatum with saline, NGF solution, and NGF coacervate injections. Ordinary one-way ANOVA with Tukey’s multiple comparisons test. E) Quantification of percent change in intensity of ChAT staining from ipsilateral to contralateral striatum with saline, NGF solution, and NGF coacervate injections. Ordinary one-way ANOVA with Tukey’s multiple comparisons test. F) Radial intensity of ChAT staining from injection site. G) Radial intensity of ChAT staining from center of contralateral striatum. H) Cumulative ChAT intensity radially from injection site. 2way ANOVA with Tukey’s multiple comparisons test. Comparisons shown on graph are from final point and compared to saline. I) Cumulative ChAT intensity radially from center of contralateral striatum. 2way ANOVA with Tukey’s multiple comparisons test. Comparisons shown on graph are from final point and compared to saline. (F, G) 2way ANOVA with Tukey’s multiple comparisons test performed on every 50^th^ pixel in radial angle profile – sections indicate where minimum P value is consistent. All comparisons made to saline. All graphs show mean ± SEM with n=5-8. * P < 0.05, ** P < 0.01, *** P < 0.001, **** P < 0.0001.

## Discussion

Protein-based therapies hold promise for improving CNS disorder outcomes, but translation has been hampered by ineffective delivery. Biomaterial-based systems can improve the spatiotemporal control of local protein delivery to the CNS, although new bioengineering solutions are needed to address the limitations of currently available tools, particularly for loading and long-term stability of bioactive cargo as well as the extent of neuropil disruption that is caused by injection or implantation of these systems. Here, we address these issues by developing a trehalose-based complex coacervate platform that shows facile protein loading, tunable protein release, cytocompatibility, regulation of intra- and extra-cellular protein delivery, and a capacity to be dispersed within neural tissue with minimal induced FBR. Our findings demonstrate proof-of-principle that trehalose-based coacervates can be used to effectively deliver bioactive proteins to cultured neural cells *in vitro* and locally to CNS tissue *in vivo* and have broad implications for: (i) using the trehalose-based coacervate platform as a versatile delivery vehicle and biotechnology tool, and (ii) applications of the coacervate-based therapies to treat CNS disorders.

Here we show that trehalose-based coacervates can load a variety of model proteins with a capacity to concentrate proteins directly from solutions while maintaining protein solubility and structural stability. There is growing interest in using coacervates to address persistent and ubiquitous challenges facing the delivery, formulation, manufacture, and supply chain management of therapeutic biomolecules such as growth factors, antibodies, and viral vectors [65–67]. When applied to such therapeutics, coacervates formed using synthetic polypeptides or charged polysaccharides have already improved the capacity to prepare injectable, high concentration monoclonal antibody formulations [67], enhance the delivery of growth factors to stimulate angiogenesis [43], and confer biomolecule thermostability improvements to address cold chain challenges [66]. However, these existing coacervate technologies make use of linear polyelectrolytes that are not biodegradable and whose electrostatic interactions with biomolecules cannot be easily regulated, which poses problems for controlled protein release applications. The branched oligomer synthesis scheme used in the trehalose-based coacervate platform enables facile control of both trehalose content and charge density, and resulting oligomers are amenable to biodegradation by ester hydrolysis. Thus, by having control over oligomer-biomolecule charge interactions and the resorption of the coacervates, this trehalose-based system represents an important advance in coacervate technologies for drug delivery applications. The unique hydrogen bonding characteristics of trehalose confer protein stabilization properties superior to all other types of saccharides or small molecule excipients [30, 68]. Multivalent incorporation of trehalose and the enhanced chain homogeneity provided by the step-growth Michael addition reaction mechanism used here to form the branched oligomers has the potential to further improve upon the stabilization capacity of both free trehalose and other previously developed coacervate systems [27]. Given the modular nature of trehalose-based coacervates, important next steps include dissecting the formulation space with different ionizable end-caps and multifunctional thiol molecules to tune coacervate properties such as electrostatic strength and branching structure.

While the focus of this study was on protein delivery to the CNS, this work opens exciting new research opportunities to extend the trehalose-based coacervate platform into a variety of other applications. For example, formulating coacervates with other molecular cargo such as small molecules or viral vectors could extend the utility of the coacervates as a general delivery tool. Mixing of differently loaded coacervates into a single bulk coacervate structure may enable precise temporal control over the release of multiple different drug cargos. Because of their substantial protein concentrating capacity and tunable properties, trehalose-based coacervates may also be a useful tool for protein purification processes where enrichment of specific proteins from dilute solutions could be achieved. Additionally, by being able to effectively concentrate biomolecules, trehalose-based coacervates could serve important new roles as catalytic sites for different chemical reactions.

The unique capacity to precisely tune the formulation of trehalose-based coacervates to encourage either extracellular retention or intracellular uptake of coacervates and its cargo affords unique application opportunities for protein delivery that are not possible with other currently available coacervate systems. By formulating coacervates to bind to but not be internalized by cells *in vitro*, trehalose-based coacervates could be used as a practical bioprinting tool to grow cell sheets or organoids that could then be readily transferred or molded into distinct structures by leveraging the malleable, viscoelastic properties of the coacervates. The distinct capacity for extracellular remaining trehalose-based coacervates to sequester and stabilize growth factors from local environments, as demonstrated by our studies here, may also further enhance such *in vitro* tissue engineering efforts.

Developing safe and effective ways to direct intracellular delivery of therapeutic proteins affords a path towards new therapies for a multitude of diseases [69, 70]. Here we showed that formulating coacervates with an excess of polycationic trehalose-based oligomers enables functional intracellular protein delivery to cultured neural cells with minimal effect on cell viability. These results inspire exciting future intracellular protein delivery possibilities for trehalose-based coacervates including using them to deliver exogenous transcription factors or ribonucleoprotein complexes to regulate cell fate and functions that could enable improved wound repair outcomes or precise corrections of cell dysfunctions to address degenerative or neoplastic diseases. While other coacervate systems have also demonstrated an intracellular delivery capacity [57, 58], the cellular uptake pathways used by coacervates are still poorly understood and improvements to delivery efficacy are required for clinical translation. For trehalose-based coacervates, it is possible that trehalose content, coacervate size, or protein corona induced by sequestering proteins from the local environment may also alter intracellular delivery. Future studies should focus on dissecting these mechanisms and precisely tuning charge imbalance and/or coacervate droplet size to optimize intracellular delivery outcomes. Independent of protein delivery, effective intracellular accumulation of trehalose-based coacervates alone may also prove to be useful in the manufacture of clinical cell therapies, since introduction of low concentrations of intracellular trehalose has afforded enhanced cryoprotection during processing [71, 72].

Trehalose-based coacervates also represent an important advance over shear thinning hydrogel systems for *in vivo* CNS neural parenchyma drug delivery. Previously, we have shown that hydrogel systems are useful tools for delivering growth factors to CNS injury lesion environments during the acute stage of injury [3, 4], but when applied to healthy neural parenchyma, hydrogels disrupt large volumes of neural tissue at the precise site where bioactive protein is to be delivered [24]. Here we demonstrated that trehalose-based coacervates can be loaded with higher concentrations of therapeutic proteins and are readily dispersed into discrete volumes of neural parenchyma to deliver bioactive drugs to neurons and glia without causing any of the extensive neural tissue displacement or inducing a substantial foreign body response, as seen with hydrogel injections. The proof-of-principle *in vivo* studies conducted in this work provide a strong foundation on which to explore numerous potential applications of the trehalose-based coacervate technology across CNS disorders in future studies. For example, trehalose-based coacervates will be useful for improving the delivery of chemoattracting factors for guiding stimulated axon regeneration after CNS injury, since we currently rely upon shear thinning hydrogels or viral vectors to deliver such factors [3, 4]. Use of coacervates in CNS injury applications could also be extended to augment wound repair functions of astrocytes by delivering exogenous, or sequestering endogenous, growth cues that enhance astrocyte proliferation and migration [73]. Coacervates may also be useful for enhancing the bioactivity of delivered enzymes to effectively debride fibrotic lesions that pose an obstacle for axon regeneration [74] or afford precise remodeling of peri-neuronal nets to promote local neural circuit plasticity [75]. Furthermore, since trehalose-based coacervates can be formulated to bind to but not be internalized by neural cells, they have the potential to serve as new carriers for exogenous grafts. In such an application, their sustained delivery properties can be used to provide local growth cues, their protein sequestering properties can be used to concentrate endogenous supportive cues, and their viscoelastic and shear-thinning properties can provide protection to cells during *in vivo* infusion, while still enabling necessary dispersion of transplanted cells within the grafting site.

In conclusion, this work describes the development of new trehalose-based coacervates and proof-of-principle studies demonstrating their capacity to serve as effective controlled release protein delivery systems in the CNS. The results from this study lay the foundation for further investigation of coacervate-based delivery of proteins and other biomolecules in models of CNS disease to direct specific therapeutic outcomes.

## Experimental procedures

### Resource availability

#### Lead contact

Further information and requests for resources and materials should be directed to and will be fulfilled by the lead contact, Timothy M. O’Shea.

### Materials

Trimethylolpropane tris(3-mercaptopropionate) (TMPTMP) (381489), 2-Carboxyethyl acrylate (55234), 2-(Dimethylamino)ethyl acrylate (330957), 3-Sulfopropyl acrylate potassium salt (251631), 2-(Acryloyloxy)ethyl trimethylammonium chloride solution (496146), 3,3′,5,5′-Tetramethylbenzidine (TMB) Liquid Substrate System for ELISA (T0440), Sodium hydroxide 10M solution (72068), Hydrochloric acid solution (6N) (1430071000), 5,5′-Dithiobis(2-nitrobenzoic acid) (D218200), Lysozyme (L6876), Human IgG-FITC (F9636), Fluorescein isothiocyanate isomer I (F7250), Peroxidase from horseradish (77332), Fluorescein isothiocyanate–dextran (average molecular weights 10 kDa (FD10S) 70 kDa (FD70S), 150 kDa (46946)), Rhodamine B isothiocyanate–Dextran (R8881), Fluorescein o-acrylate (568856), Albumin-fluorescein isothiocyanate conjugate (A9771), Saporin (S9896), and Super-DHB matrix (50862), were purchased from Sigma-Aldrich. Lysozyme was FITC-labelled by reacting fluorescein isothiocyanate with lysozyme at a 1:10 molar ratio in 0.1 M borate buffer (pH 8.5) for 4 hours at 4°C. The reaction was then dialyzed against deionized (DI) H_2_O for 48 hours at 4°C protected from light using a 3.5 kDa cutoff dialysis bag. The product was filtered, lyophilized, and stored at 4°C for use. Trehalose dihydrate (T0331), 2-(Dimethylamino)ethyl acrylate (A1235), 4-methoxyphenol (MEHQ) (M0123), N,N-Diisopropylethylamine (D1599), Ceric Ammonium Molybdate Solution (C1794), and 2,2,2-Trifluoroethyl acrylate (TFEA) (A1152) were purchased from TCI Chemicals. Celite 545 (AC349675000), 4A Molecular Sieves (AAL054540B), 2-Methyl-2-butanol (2M2B) (AAA183040F), Bovine Serum Albumin (BSA) (BP9706100), Agarose (IB70040), alpha-Cyano-4-hydroxycinnamic acid (J67635.EXK), and activated alumina (Brockman I, basic) (6002042) were purchased from Thermo Scientific. Novozym 435 was received as a gift from Novozymes. Disposable 40g HP Silica Gold Cartridges were purchased from Teledyne Isco. All solvents (methanol, ethyl acetate, acetone, dimethyl sulfoxide (DMSO), N,N-Dimethylformamide (DMF)) were ACS grade and purchased from Fisher Scientific.

### Synthesis of trehalose-based oligomers

Trehalose diacrylate was synthesized by reacting 8 molar excess of TFEA with lyophilized trehalose dihydrate in the presence of CALB at 50°C for 1 week. The reaction was solubilized in dry 2M2B with 2% DMSO and included 4A molecular sieves (4AMS) as well as MEHQ to prevent radical acrylate homopolymerization. The crude product was filtered to remove CALB and 4AMS and concentrated *in vacuo* using rotary evaporation. The product was precipitated into ethyl acetate, loaded onto Celite 545, and purified via flash chromatography using the ISCO CombiFlash system using a 40g silica cartridge and ethyl acetate and 80:20 methanol:deionized H_2_O as the solvent mobile phases. Purified trehalose diacrylate was identified using thin layer chromatography (TLC) developed using CAM staining and collected fractions were dried *in vacuo* using rotary evaporation. A small quantity of MEHQ at approximately 40 mg per gram of reaction was included in the product to prevent radical acrylate homopolymerization during purification. The product was stored at 4°C under desiccant until use.

TMPTMP and all liquid acrylate end-caps were passed through basic alumina columns to remove the MEHQ inhibitor prior to use in oligomer synthesis reactions. Stoichiometrically defined amounts of TMPTMP, trehalose diacrylate, and mono-acrylate end-cap required for desired oligomer branching functionality (f_b_) were solubilized at 10 wt% in DMF or 75/25 DMF/water. Typical reactions were prepared at 100-500 mg scale, catalyzed by 1/3 molar equivalent to thiols of DIEA and allowed to proceed for approximately 12 hours stirring at room temperature. Reaction progression was monitored using TLC that was developed using Ellman’s stain to confirm the consumption of thiols in the reaction. Reactions were concentrated by rotary evaporation to remove DIEA and DMF. The crude oligomer product was solubilized in DI H_2_O aided by appropriate acidification or alkalization achieved by adding one molar equivalent of HCl or NaOH to end-cap. Residual DMF and DIEA was removed via liquid-liquid extraction with ethyl acetate and trace ethyl acetate was removed *in vacuo* by rotary evaporation. The resulting water-soluble oligomer was lyophilized and stored at 4°C until use.

### Characterization of trehalose-based oligomers

^1^H NMR spectra of trehalose diacrylate and synthesized oligomers were collected on an Agilent 500 MHz VNMRS spectrometer with a Varian ultra shielded magnet at concentrations of approximately 10 mg/mL in DMSO-d6. Hydrogen chemical shifts are expressed in parts per million (ppm) relative to the residual proton solvent resonance: DMSO-d6 δ2.50 or δ3.33. Number Molecular weight of branched oligomer was calculated using NMR. Fourier Transform Infrared Spectroscopy (FTIR) was performed using a Nicolet 4700 FTIR-ATR between the wavenumber region 4000-400 cm^-1^ (64 scans, 0.241 cm^-1^ resolution). Zeta potential measurement of constitutively charged oligomers solubilized at 10 mg/ml in water was performed using a Brookhaven NanoBrook Particle Size Analyzer. Solubility of oligomers was assessed using a series of pH adjusted buffers containing 10 mM HEPES, 10 mM MES, 10 mM sodium acetate, and 140 mM sodium chloride with pH values ranging from 3-8 in a 96-well plate with absorbance measured at 450 nm on the Molecular Devices SpectraMax i3X Microplate Detection Platform. pKa measurements of ionizable oligomers was performed using 2-(ptoluidino)-naphthalene-6-sulfonic acid (TNS) assay. For TNS assay, oligomers were solubilized in pH adjusted buffers at a concentration of 4 mg/ml prior to addition of TNS at a concentration of 2 µM. Sample fluorescence was measured in a 96-well plate using the same plate reader (excitation/emission 321/445 nm). A sigmoidal fit analysis was applied to the fluorescence data to determine the pKa value. MALDI-TOF data was collected using a Bruker autoflex speed instrument. Oligomer samples were prepared at 10 mg/ml in a mixture of methanol and water. DHB matrix was prepared at 25 mg/ml in 50/50 methanol/water. Matrix and oligomer were mixed at a ratio of 20:1 and dried onto the target. Negative and positive linear methods were used according to the oligomer end-cap charge. FlexAnalysis software was used to obtain mass lists from collected spectra.

### Formulation of trehalose-based coacervates

To form coacervates, oligomers were solubilized in DI water at 500 mg/ml and diluted into appropriate buffers at concentrations ranging from 1-100 mg/ml. Phosphate buffered saline with pH values ranging from 6-8 were used across coacervate characterization experiments. Polycation and polyanion oligomers were combined at charge equivalency or at defined molar ratios to form coacervates. Coacervates loaded with protein cargo were formulated with a desired loading ratio (LR, oligomer mass/protein mass (mg/mg)) by combining the protein of interest prepared in a solution at 1-10 mg/ml in phosphate buffered saline with the polyionic oligomer of opposite charge (the appropriate oligomer was chosen based on the known isoelectric point for the protein). The oligomer with the same charge as the protein was then incorporated at the desired molar ratio to the protein-oligomer solution to generate coacervates.

### Characterization of trehalose-based coacervates

Coacervate formulations were characterized by turbidity measurements that were performed in a 96-well plate with absorbance measured at 450 nm on a Molecular Devices SpectraMax i3X Microplate Detection Platform. Coacervate phase contrast images were acquired on Fisherbrand Inverted Microscope with Hayear camera attachment. Fluorescence imaging of FITC-labeled protein-loaded coacervates and fluorescein-labeled coacervates were obtained using an Olympus IX83 microscope. Rheological measurements of bulk coacervates were taken on a TA Instruments DHR-2 Rheometer, at 25°C with an 8 mm plate geometry. Coacervate nanodroplet diameter and polydispersity upon dilution and during disassembly *in vitro* were acquired using dynamic light scattering on a Brookhaven NanoBrook Particle Size Analyzer. Degradation of coacervates by hydrolysis was assessed by monitoring the emergence of a carboxylate (1550–1610 cm^−1^) stretch signal by FTIR. Degradation and disassembly of coacervates was monitored by quantifying fluorescence using a Molecular Devices SpectraMax i3X Microplate Detection Platform. Fluorescence recovery after photobleaching (FRAP) was conducted on an Olympus IX83 widefield microscope. Fluorescein-labeled bulk coacervates were imaged on a glass coverslip before photobleaching and then exposed to light generated by a LED Light Source (IX3-IL) passed through a 488 nm filter and 20X objective for approximately 3 minutes. The bulk coacervate was then imaged at the same location periodically over the course of 1 hour. NIH Image J software was used to quantify change in fluorescence intensity.

### Characterization of biomacromolecule encapsulation and release *in vitro* from coacervates

Coacervates were loaded with model biomacromolecules including FITC-dextrans, FITC-Lysozyme, FITC-BSA or FITC-IgG. Encapsulation efficiency of model biomacromolecules in coacervates was determined by centrifuging the formed coacervates into a pellet and then evaluating the depletion of protein from the resultant coacervate supernatant using microplate-based fluorescence detection (excitation/emission 488/520 nm). Concentrating factor was calculated by measuring the total volume of the resultant coacervate pellet and accounting for the total protein encapsulated based on the above encapsulation efficiency measurements. For protein release, studies coacervates were incubated in 200 µL of PBS at 37°C with the incubation solution replaced daily. Protein concentrations in the recovered incubation solutions were read daily using microplate-based fluorescence detection. To benchmark protein release, eight-sample serially diluted standard curves were generated for each protein to convert the microplate measured fluorescence intensity into a protein concentration value. To model retention of coacervates and release of biomacromolecules in *in vitro* neural tissue-like environments, cargo loaded coacervates were injected into a 0.6 wt% agarose gel phantom brain[54]. Agarose phantoms were incubated in a PBS solution and the concentration of model biomacromolecules released from the agarose phantoms was characterized by microplate-based fluorescence detection. Bioactive protein recovery from coacervates was performed using HRP as a model enzyme. Active HRP in recovered incubation solutions was measured by developing with tetramethylbenzidine ELISA substrate for 5 minutes followed by adding 100 µL of 2N HCl solution to stop the enzymatic reaction. Absorbance was read at 450 nm on microplate reader and compared against a HRP solution standard. All plate reader assays were performed on a Molecular Devices SpectraMax i3X Microplate Detection Platform. Agarose gel images were obtained by placing gels with injected solution or coacervate on a coverslip and imaging using IX83 Olympus epifluorescence microscope at discrete timepoints.

### *In vitro* cell culture assays and immunocytochemistry

Cytocompatibility of coacervates and individual oligomers *in vitro* with mouse neural progenitor cells (NPC) and differentiated astrocytes was assessed using Calcein AM cell viability assay. NPC were generated and cultured in neural expansion media supplement with EGF and FGF growth factors using standard methods described in detail previously by our team [76]. Astrocytes were derived by treating NPC with media containing 1% FBS for 2-4 days as we have done previously [55, 76]. Cells were seeded on tissue culture treated 96-well plates at 50,000 cells per well and treated 24 or 48 hours later with coacervates suspended in media. Cell viability was assessed at 24 hours post-treatment by microplate-based fluorescence detection of Calcein AM (Cat# 354217, Corning) which was prepared at a molarity of 2µM. Calcein AM solution was incubated with cells for 30min and read on a Molecular Devices SpectraMax i3X Microplate Detection Platform using an emission/excitation of 488/520 nm. Results were normalized with untreated control cells representing 100% viability and 80% methanol treated (dead) cells representing 0% viability. To assess coacervate-mediated growth factor depletion and supplementation of media, NPC were plated in a 96-well plate at 50,000 cells/well in standard media conditions and allowed to adhere overnight. Then, concentrated coacervates were formed at high concentration in PBS and the suspension was diluted into a volume of growth factor-containing media. The media was then spun down and the supernatant media was collected and added to cells. The coacervate pellets were then resuspended in media without added growth factor and added to cells. Cell number was assessed after 4 days using the same Calcein AM assay methods outlined above. To assess intracellular protein delivery, NPC were plated in a 96 well plate at 50,000 cells/well in standard media conditions and allowed to adhere overnight. Then, saporin was diluted into media at various concentrations and individual oligomers were added to the media. The resulting coacervate-suspension was added to cells. After 3 days of treatment, cell viability was measured using the same Calcein AM assay methods outlined above. To assess cell attachment to bulk coacervates, individual wells of a non-treated polystyrene 96-well plate were coated with 2 mg/ml coacervate or individual oligomer solutions followed by addition of 50,000 cells per well. After 24 hours, non-adhered floating cells were transferred to a tissue culture treated 96-well plate. After an additional 24 hours of culture, both plates were treated with Calcein AM and cell numbers were evaluated by microplate fluorescence. All plate reader assays were performed on a Molecular Devices SpectraMax i3X Microplate Detection Platform. Cell interactions with coacervates were evaluated using fluorescein-labeled coacervates or coacervates loaded with FITC-BSA. NPC or astrocytes were seeded on gelatin coated glass coverslips and then treated with coacervates for 4 hours or 24 hours. Cell coverslips were fixed with 4% paraformaldehyde (prepared from 32% PFA Aqueous Solution (Cat# 15714, EMS)) with cell morphology and nucleus visualized using Acti-stain 555 phalloidin (1:2000, Cytoskeleton, PHDH1A) and DAPI (2ng/ml; Molecular probes) respectively. Coverslips were mounted onto microscope slides using Prolong Gold antifade reagent (Invitrogen, Cat# P36934) and imaged using an Olympus IX83 Widefield epifluorescence microscope.

### Surgical procedures

All surgical procedures were approved by the BU IACUC (Protocol number: PROTO20200045) and conducted within a designated surgical facility. All procedures were performed on C57/BL6 female mice (Cat#000664, JAX) that were aged 8-12 weeks at the time of craniotomy. Mice were positioned into a rodent stereotaxic apparatus using ear bars (David Kopf, Tujunga, CA) and maintained under general anesthesia achieved through inhalation of isoflurane (1-3%) in oxygen-enriched air. Using the aid of a stereo microscope, a small rectangular shaped craniotomy was performed over the left coronal suture using a high-speed surgical drill. A small flap of bone encompassing sections of the frontal and parietal bone was removed to expose the surface of the brain prior to injection. Bulk coacervates (100% v/v), suspended coacervates (10% v/v in PBS), BDA solution (0.4 wt%), NGF solution (2 mg/ml), or saline (1xPBS) were loaded into pulled borosilicate glass micropipettes (WPI #1B100-4) that were ground to a 35° beveled tip with 150–250 μm inner diameter after formulation with sterile-filtered components. Encapsulation of BDA in coacervates yielded a concentrating factor of ∼1, ensuring the same concentration of BDA injected in solution and coacervate conditions. Glass micropipettes were mounted to the stereotaxic frame and high-pressure polyetheretherketone tubing and connectors were used to attach micropipettes to a 10 μL syringe (Hamilton, Reno, NV, #801 RN) that was mounted into a syringe pump (Pump 11 Elite, Harvard Apparatus). Coacervates or PBS containing either BDA (10 kDa—Thermofisher, #D1956) or recombinant human β-NGF (PeproTech, Cat# 450-01) were injected into the caudate putamen nucleus of the striatum at 0.15 μL/min using target coordinates relative to Bregma: +1.0 mm A/P, +2.5 mm L/M and −3.0 mm D/V. A total volume of 1 μL was used for all conditions. Once the complete volume of coacervate or solution had been injected the micropipette was allowed to dwell in the brain at the injection site for an additional 4 minutes before being slowly removed from the brain incrementally over a 2-min period. After injection, the surgical incision site was sutured closed with polyethylene sutures and the animals were allowed to recover. To benchmark results using coacervates with a bulk biomaterial, we used injections of 5 wt% methyl cellulose (MC) (Methocel A15C, [MW = 304 kDa; 27.5–31.5% substitution], Sigma #64625) hydrogels loaded with BDA as a control group in these studies.

### Transcardial perfusion and immunohistochemistry

At 24 hours or 7 days after striatal injections, mice underwent terminal anesthesia by overdose of isoflurane and were then perfused transcardially with heparinized saline and 4% paraformaldehyde (PFA) using a peristaltic pump at a rate of 7 mL/min. Approximately, 10 mL of heparinized saline and 50 mL of 4% PFA were used per animal. Brains were dissected and post-fixed in 4% PFA for 6-8 hours followed by cryoprotection in 30% sucrose with 0.01% sodium azide in Tris-Buffered Saline (TBS) for at least 3 days at 4°C prior to cryosectioning. Coronal brain sections (40 µm thick) were cut using a cryostat (Leica CM1950 Cryostat). Tissue sections were stored in 96well plates immersed with 1X TBS and 0.01% sodium azide solution at 4°C. For immunohistochemistry (IHC), tissue sections were stained using a free-floating staining protocol outlined in detail previously[24, 55]. Following 1N HCl antigen retrieval and serial washes, the tissue sections were blocked and permeabilized using 1X TBS containing 5% Normal donkey serum (NDS, Jackson ImmunoResearch Laboratories) and 0.5% Triton x-100 for one hour. Tissue sections were then stained with primary antibodies overnight in 1X TBS containing 0.5% Triton x-100. The following primary antibodies were used in this study: anti-Rabbit NeuN (Abcam, ab177487, 1:1000); anti-Rat Gfap (Invitrogen, 13-0300, 1:1000); anti-Guinea pig Iba-1 (Synaptic Systems, 234 004, 1:500); anti-Rabbit P2ry12 (Ana Spec, AS-55043A, 1:800); anti-Goat Cd13 (R&D systems, AF2335, 1:500); anti-Rat Ly6b2 (BioRad, MCA771G, 1:200); anti-Goat ChAT (MilliporeSigma, AB144P, 1:500). Tissue sections were washed three times and incubated with appropriate secondary antibodies diluted at 1:250 in 1X TBS containing 0.5% Triton x-100 and 5% NDS for two hours. All secondary antibodies were affinity purified whole IgG(H + L) purchased from Jackson ImmunoResearch Laboratories, with donkey host and target specified by the primary antibody. BDA was visualized using streptavidin-HRP plus Tyr-Cy3 (PerkinElmer). Cell nuclei were counter stained with 4’,6’-diamidino-2-phenylindole dihydrochloride (DAPI; 2 ng/ml; Molecular Probes) prior to mounting onto glass slides and then cover slipped with ProLong Gold (Invitrogen, Cat# P36934) anti-fade reagent. Stained sections were imaged using epifluorescence and deconvolution epifluorescence microscopy on an Olympus IX83 microscope.

### Quantitative analysis of immunohistochemistry

Immunohistochemical staining intensity quantification for the various antibody markers or BDA was performed on tiled images prepared on the Olympus IX83 epifluorescence microscope that were taken at a standardized exposure time across groups and using the raw/uncorrected intensity settings. Quantification of total antibody staining or BDA accumulated into tissue was performed using NIH Image J (1.51) software and the Radial Profile Angle plugin as we have done before[24]. Total values for IHC stainings or detected BDA were determined by taking the integral (area under the curve) of the radial intensity profile. Bioactive NGF delivery was assessed by characterizing the relative change in size of ChAT- and NeuN-positive cholinergic neurons and ChAT-positive neuropil on the ipsilateral striatum treated with NGF solution or coacervate compared to the contralateral striatum using the analyze particles function in NIH Image J (1.51) as done previously[24].

### Statistical analysis

Statistical evaluations were conducted by one-way or 2way ANOVA with post hoc independent pairwise analysis via Tukey’s or Sidak’s multiple comparisons test where appropriate using Prism 10 (GraphPad Software Inc, San Diego, CA). Across all statistical tests, significance was defined as p-value <0.05. All graphs show mean values plus or minus standard error of the means (S.E.M.) as well as individual values overlaid as dot plots.

### Data availability statement

All data generated for this study are included in the main and supplementary figures. For all quantitative figures, the results of statistical tests are provided with the paper. Files of source data of individual values and other data that support the findings of this study are available on reasonable request from the corresponding author.

## Acknowledgements

This research is supported by the Biomedical Engineering Core Facilities at Boston University. A special thanks to Xin Brown and the Biointerface Technologies (BIT) core, as well as the Micro and Nano Imaging (MNI) core at BU. This research was also supported by the BU Chemistry Department Chemical Instrumentation Center (CIC) and staff. We are grateful to the National Science Foundation for the purchase of the NMR (CHE0619339) and MALDI-TOF mass spectrometer (CHE1337811) used in this work. This work was supported by internal Boston University start-up funds (T.M.O) and the Wings for Life Spinal Cord Research Foundation (T.M.O).

## References

[1] R.G. Thorne, W.H. Frey, Delivery of Neurotrophic Factors to the Central Nervous System, Clinical Pharmacokinetics 40(12) (2001) 907–946.

[2] E. Nance, S.H. Pun, R. Saigal, D.L. Sellers, Drug delivery to the central nervous system, Nature Reviews Materials 7(4) (2022) 314–331.

[3] M.A. Anderson, J.E. Burda, Y. Ren, Y. Ao, T.M. O’Shea, R. Kawaguchi, G. Coppola, B.S. Khakh, T.J. Deming, M.V. Sofroniew, Astrocyte scar formation aids central nervous system axon regeneration, Nature 532 (2016) 195.

[4] M.A. Anderson, T.M. O’Shea, J.E. Burda, Y. Ao, S.L. Barlatey, A.M. Bernstein, J.H. Kim, N.D. James, A. Rogers, B. Kato, A.L. Wollenberg, R. Kawaguchi, G. Coppola, C. Wang, T.J. Deming, Z. He, G. Courtine, M.V. Sofroniew, Required growth facilitators propel axon regeneration across complete spinal cord injury, Nature 561(7723) (2018) 396–400.

[5] M.V. Sofroniew, C.L. Howe, W.C. Mobley, Nerve Growth Factor Signaling, Neuroprotection, and Neural Repair, Annu Rev Neurosci 24(1) (2001) 1217–1281.

[6] T. Kobayashi, H. Ahlenius, P. Thored, R. Kobayashi, Z. Kokaia, O. Lindvall, Intracerebral infusion of glial cell line-derived neurotrophic factor promotes striatal neurogenesis after stroke in adult rats, Stroke 37(9) (2006) 2361–7.

[7] J.M. Obermeyer, A. Tuladhar, S.L. Payne, E. Ho, C.M. Morshead, M.S. Shoichet, Local Delivery of Brain-Derived Neurotrophic Factor Enables Behavioral Recovery and Tissue Repair in Stroke-Injured Rats, Tissue Engineering Part A 25(15-16) (2019) 1175–1187.

[8] S.S. Kang, M.P. Keasey, S.A. Arnold, R. Reid, J. Geralds, T. Hagg, Endogenous CNTF mediates stroke-induced adult CNS neurogenesis in mice, Neurobiology of disease 49 (2013) 68–78.

[9] J. Houlton, N. Abumaria, S.F.R. Hinkley, A.N. Clarkson, Therapeutic Potential of Neurotrophins for Repair After Brain Injury: A Helping Hand From Biomaterials, Front Neurosci 13(790) (2019).

[10] J.W. Squair, M. Milano, A. de Coucy, M. Gautier, M.A. Skinnider, N.D. James, N. Cho, A. Lasne, C. Kathe, T.H. Hutson, S. Ceto, L. Baud, K. Galan, V. Aureli, A. Laskaratos, Q. Barraud, T.J. Deming, R.E. Kohman, B.L. Schneider, Z. He, J. Bloch, M.V. Sofroniew, G. Courtine, M.A. Anderson, Recovery of walking after paralysis by regenerating characterized neurons to their natural target region, Science (New York, N.Y.) 381(6664) (2023) 1338–1345.

[11] A. Eslamboli, B. Georgievska, R.M. Ridley, H.F. Baker, N. Muzyczka, C. Burger, R.J. Mandel, L. Annett, D. Kirik, Continuous low-level glial cell line-derived neurotrophic factor delivery using recombinant adeno-associated viral vectors provides neuroprotection and induces behavioral recovery in a primate model of Parkinson’s disease, The Journal of neuroscience : the official journal of the Society for Neuroscience 25(4) (2005) 769–77.

[12] M.H. Tuszynski, Intraparenchymal NGF infusions rescue degenerating cholinergic neurons, Cell Transplant 9(5) (2000) 629–36.

[13] A. Whone, M. Luz, M. Boca, M. Woolley, L. Mooney, S. Dharia, J. Broadfoot, D. Cronin, C. Schroers, N.U. Barua, L. Longpre, C.L. Barclay, C. Boiko, G.A. Johnson, H.C. Fibiger, R. Harrison, O. Lewis, G. Pritchard, M. Howell, C. Irving, D. Johnson, S. Kinch, C. Marshall, A.D. Lawrence, S. Blinder, V. Sossi, A.J. Stoessl, P. Skinner, E. Mohr, S.S. Gill, Randomized trial of intermittent intraputamenal glial cell line-derived neurotrophic factor in Parkinson’s disease, Brain 142(3) (2019) 512–525.

[14] G. García-Alías, S. Barkhuysen, M. Buckle, J.W. Fawcett, Chondroitinase ABC treatment opens a window of opportunity for task-specific rehabilitation, Nature Neuroscience 12(9) (2009) 1145–1151.

[15] W.B.J. Cafferty, E.J. Bradbury, M. Lidierth, M. Jones, P.J. Duffy, S. Pezet, S.B. McMahon, Chondroitinase ABC-Mediated Plasticity of Spinal Sensory Function, The Journal of Neuroscience 28(46) (2008) 11998–12009.

[16] A.D. Zurn, H.R. Widmer, P. Aebischer, Sustained delivery of GDNF: towards a treatment for Parkinson’s disease, Brain Research Reviews 36(2) (2001) 222–229.

[17] G.M. Thomsen, M. Alkaslasi, J.P. Vit, G. Lawless, M. Godoy, G. Gowing, O. Shelest, C.N. Svendsen, Systemic injection of AAV9-GDNF provides modest functional improvements in the SOD1(G93A) ALS rat but has adverse side effects, Gene Ther 24(4) (2017) 245–252.

[18] A. Arvidsson, D. Kirik, C. Lundberg, R.J. Mandel, G. Andsberg, Z. Kokaia, O. Lindvall, Elevated GDNF levels following viral vector-mediated gene transfer can increase neuronal death after stroke in rats, Neurobiology of Disease 14(3) (2003) 542–556.

[19] M. Luz, E. Mohr, H.C. Fibiger, GDNF-induced cerebellar toxicity: A brief review, NeuroToxicology 52 (2016) 46–56.

[20] S.J. Chan, C. Love, M. Spector, S.M. Cool, V. Nurcombe, E.H. Lo, Endogenous regeneration: Engineering growth factors for stroke, Neurochemistry International 107 (2017) 57–65.

[21] C.J. Kearney, D.J. Mooney, Macroscale delivery systems for molecular and cellular payloads, Nature Materials 12 (2013) 1004.

[22] D.J. Cook, C. Nguyen, H.N. Chun, I. L Llorente, A.S. Chiu, M. Machnicki, T.I. Zarembinski, S.T. Carmichael, Hydrogel-delivered brain-derived neurotrophic factor promotes tissue repair and recovery after stroke, Journal of Cerebral Blood Flow & Metabolism 37(3) (2016) 1030–1045.

[23] C.P.J. Hunt, V. Penna, C.W. Gantner, N. Moriarty, Y. Wang, S. Franks, C.M. Ermine, I.R. de Luzy, C. Pavan, B.M. Long, R.J. Williams, L.H. Thompson, D.R. Nisbet, C.L. Parish, Tissue Programmed Hydrogels Functionalized with GDNF Improve Human Neural Grafts in Parkinson’s Disease, Advanced Functional Materials n/a(n/a) (2021) 2105301.

[24] T.M. O’Shea, A.L. Wollenberg, J.H. Kim, Y. Ao, T.J. Deming, M.V. Sofroniew, Foreign body responses in central nervous system mimic natural wound responses and alter biomaterial functions, Nature Communications 11(6203) (2020).

[25] J. Lee, J.H. Ko, K.M. Mansfield, P.C. Nauka, E. Bat, H.D. Maynard, Glucose-Responsive Trehalose Hydrogel for Insulin Stabilization and Delivery, Macromolecular Bioscience 18(5) (2018) 1700372.

[26] J. Lee, E.-W. Lin, U.Y. Lau, J.L. Hedrick, E. Bat, H.D. Maynard, Trehalose Glycopolymers as Excipients for Protein Stabilization, Biomacromolecules 14(8) (2013) 2561–2569.

[27] T.M. O’Shea, M.J. Webber, A.A. Aimetti, R. Langer, Covalent Incorporation of Trehalose within Hydrogels for Enhanced Long-Term Functional Stability and Controlled Release of Biomacromolecules, Advanced Healthcare Materials 4(12) (2015) 1802–1812.

[28] D. Vinciguerra, M.B. Gelb, H.D. Maynard, Synthesis and Application of Trehalose Materials, JACS Au 2(7) (2022) 1561–1587.

[29] J.H. Crowe, L.M. Crowe, Preservation of mammalian cells-learning nature’s tricks, Nat Biotechnol 18(2) (2000) 145–6.

[30] N.K. Jain, I. Roy, Effect of trehalose on protein structure, Protein Sci 18(1) (2009) 24–36.

[31] S.S. Buchanan, S.A. Gross, J.P. Acker, M. Toner, J.F. Carpenter, D.W. Pyatt, Cryopreservation of stem cells using trehalose: evaluation of the method using a human hematopoietic cell line, Stem Cells Dev 13(3) (2004) 295–305.

[32] E.M. DuBois, H.O. Adewumi, P.R. O’Connor, J.E. Labovitz, T.M. O’Shea, Trehalose-Guanosine Glycopolymer Hydrogels Direct Adaptive Glia Responses in CNS Injury, Advanced Materials (2023) 2211774.

[33] M.J. Mitchell, M.M. Billingsley, R.M. Haley, M.E. Wechsler, N.A. Peppas, R. Langer, Engineering precision nanoparticles for drug delivery, Nature Reviews Drug Discovery 20(2) (2021) 101–124.

[34] E.A. Nance, G.F. Woodworth, K.A. Sailor, T.-Y. Shih, Q. Xu, G. Swaminathan, D. Xiang, C. Eberhart, J. Hanes, A Dense Poly(Ethylene Glycol) Coating Improves Penetration of Large Polymeric Nanoparticles Within Brain Tissue, Sci Transl Med 4(149) (2012) 149ra119–149ra119.

[35] M. van de Weert, W.E. Hennink, W. Jiskoot, Protein Instability in Poly(Lactic-co-Glycolic Acid) Microparticles, Pharmaceutical Research 17(10) (2000) 1159–1167.

[36] S.P. Schwendeman, R.B. Shah, B.A. Bailey, A.S. Schwendeman, Injectable controlled release depots for large molecules, Journal of Controlled Release 190 (2014) 240–253.

[37] E. Astoricchio, C. Alfano, L. Rajendran, P.A. Temussi, A. Pastore, The Wide World of Coacervates: From the Sea to Neurodegeneration, Trends in Biochemical Sciences 45(8) (2020) 706–717.

[38] N.R. Johnson, Y. Wang, Coacervate delivery systems for proteins and small molecule drugs, Expert Opin Drug Deliv 11(12) (2014) 1829–32.

[39] S. Lindhoud, M.M.A.E. Claessens, Accumulation of small protein molecules in a macroscopic complex coacervate, Soft Matter 12(2) (2016) 408–413.

[40] N.A. Zervoudis, A.C. Obermeyer, The effects of protein charge patterning on complex coacervation, Soft Matter 17(27) (2021) 6637–6645.

[41] K.A. Black, D. Priftis, S.L. Perry, J. Yip, W.Y. Byun, M. Tirrell, Protein Encapsulation via Polypeptide Complex Coacervation, ACS Macro Letters 3(10) (2014) 1088–1091.

[42] Bradley W. Davis, William M. Aumiller, N. Hashemian, S. An, A. Armaou, Christine D. Keating, Colocalization and Sequential Enzyme Activity in Aqueous Biphasic Systems: Experiments and Modeling, Biophys J 109(10) (2015) 2182–2194.

[43] H. Chu, J. Gao, C.-W. Chen, J. Huard, Y. Wang, Injectable fibroblast growth factor-2 coacervate for persistent angiogenesis, Proceedings of the National Academy of Sciences 108(33) (2011) 13444–13449.

[44] S. Sen, J.E. Puskas, Green Polymer Chemistry: Enzyme Catalysis for Polymer Functionalization, Molecules 20(5) (2015) 9358–9379.

[45] G.B. Perin, M.I. Felisberti, Enzymatic synthesis and structural characterization of methacryloyl-D-fructose- and methacryloyl-D-glucose-based monomers and poly(methacryloyl-D-fructose)-based hydrogels, Journal of Chemical Technology & Biotechnology 93(6) (2018) 1694–1704.

[46] H. Chen, J. Kong, Hyperbranched polymers from A2 + B3 strategy: recent advances in description and control of fine topology, Polymer Chemistry 7(22) (2016) 3643–3663.

[47] S. Unal, T.E. Long, Highly Branched Poly(ether ester)s via Cyclization-Free Melt Condensation of A2 Oligomers and B3 Monomers, Macromolecules 39(8) (2006) 2788–2793.

[48] C.E. Sing, S.L. Perry, Recent progress in the science of complex coacervation, Soft Matter 16(12) (2020) 2885–2914.

[49] H. Lv, S. Zhang, B. Wang, S. Cui, J. Yan, Toxicity of cationic lipids and cationic polymers in gene delivery, Journal of Controlled Release 114(1) (2006) 100–109.

[50] A.M. Jörgensen, R. Wibel, A. Bernkop-Schnürch, Biodegradable Cationic and Ionizable Cationic Lipids: A Roadmap for Safer Pharmaceutical Excipients, Small n/a(n/a) (2023) 2206968.

[51] N. Kamiya, A.M. Klibanov, Controling the rate of protein release from polyelectrolyte complexes, Biotechnology and Bioengineering 82(5) (2003) 590–594.

[52] C. Zhang, P. Mastorakos, M. Sobral, S. Berry, E. Song, E. Nance, C.G. Eberhart, J. Hanes, J.S. Suk, Strategies to enhance the distribution of nanotherapeutics in the brain, Journal of Controlled Release 267 (2017) 232–239.

[53] D.J. Wolak, R.G. Thorne, Diffusion of Macromolecules in the Brain: Implications for Drug Delivery, Mol Pharm 10(5) (2013) 1492–1504.

[54] Z.J. Chen, G.T. Gillies, W.C. Broaddus, S.S. Prabhu, H. Fillmore, R.M. Mitchell, F.D. Corwin, P.P. Fatouros, A realistic brain tissue phantom for intraparenchymal infusion studies, J Neurosurg 101(2) (2004) 314–22.

[55] H.O. Adewumi, G.I. Berniac, E.A. McCarthy, T.M. O’Shea, Ischemic and hemorrhagic stroke lesion environments differentially alter the glia repair potential of neural progenitor cell and immature astrocyte grafts, Experimental Neurology 374 (2024) 114692.

[56] Q. Zhang, H.R. Phillips, A. Purchel, J.K. Hexum, T.M. Reineke, Sustainable and Degradable Epoxy Resins from Trehalose, Cyclodextrin, and Soybean Oil Yield Tunable Mechanical Performance and Cell Adhesion, ACS Sustainable Chemistry & Engineering 6(11) (2018) 14967–14978.

[57] Y. Sun, S.Y. Lau, Z.W. Lim, S.C. Chang, F. Ghadessy, A. Partridge, A. Miserez, Phase-separating peptides for direct cytosolic delivery and redox-activated release of macromolecular therapeutics, Nature Chemistry 14(3) (2022) 274–283.

[58] T. Iwata, H. Hirose, K. Sakamoto, Y. Hirai, J.V.V. Arafiles, M. Akishiba, M. Imanishi, S. Futaki, Liquid Droplet Formation and Facile Cytosolic Translocation of IgG in the Presence of Attenuated Cationic Amphiphilic Lytic Peptides, Angewandte Chemie International Edition 60(36) (2021) 19804–19812.

[59] Y. Bao, H. Chen, Z. Xu, J. Gao, L. Jiang, J. Xia, Photo-Responsive Phase-Separating Fluorescent Molecules for Intracellular Protein Delivery, Angewandte Chemie International Edition 62(42) (2023) e202307045.

[60] J.M. Horn, A.C. Obermeyer, Genetic and Covalent Protein Modification Strategies to Facilitate Intracellular Delivery, Biomacromolecules 22(12) (2021) 4883–4904.

[61] A. Vedadghavami, C. Zhang, A.G. Bajpayee, Overcoming negatively charged tissue barriers: Drug delivery using cationic peptides and proteins, Nano Today 34 (2020) 100898.

[62] B.C. Evans, R.B. Fletcher, K.V. Kilchrist, E.A. Dailing, A.J. Mukalel, J.M. Colazo, M. Oliver, J. Cheung-Flynn, C.M. Brophy, J.W. Tierney, J.S. Isenberg, K.D. Hankenson, K. Ghimire, C. Lander, C.A. Gersbach, C.L. Duvall, An anionic, endosome-escaping polymer to potentiate intracellular delivery of cationic peptides, biomacromolecules, and nanoparticles, Nature Communications 10(1) (2019) 5012.

[63] A. Tabernero, A. Velasco, B. Granda, E.M. Lavado, J.M. Medina, Transcytosis of albumin in astrocytes activates the sterol regulatory element-binding protein-1, which promotes the synthesis of the neurotrophic factor oleic acid, J Biol Chem 277(6) (2002) 4240–6.

[64] Y. Rui, D.R. Wilson, J. Choi, M. Varanasi, K. Sanders, J. Karlsson, M. Lim, J.J. Green, Carboxylated branched poly(β-amino ester) nanoparticles enable robust cytosolic protein delivery and CRISPR-Cas9 gene editing, Science Advances 5(12) (2019) eaay3255.

[65] W. Jiskoot, A. Hawe, T. Menzen, D.B. Volkin, D.J.A. Crommelin, Ongoing Challenges to Develop High Concentration Monoclonal Antibody-based Formulations for Subcutaneous Administration: Quo Vadis?, Journal of Pharmaceutical Sciences 111(4) (2022) 861–867.

[66] X. Mi, W.C. Blocher McTigue, P.U. Joshi, M.K. Bunker, C.L. Heldt, S.L. Perry, Thermostabilization of viruses via complex coacervation, Biomater Sci 8(24) (2020) 7082–7092.

[67] J.E. Bramham, S.A. Davies, A. Podmore, A.P. Golovanov, Stability of a high-concentration monoclonal antibody solution produced by liquid-liquid phase separation, MAbs 13(1) (2021) 1940666.

[68] C. Colaço, S. Sen, M. Thangavelu, S. Pinder, B. Roser, Extraordinary Stability of Enzymes Dried in Trehalose: Simplified Molecular Biology, Bio/Technology 10(9) (1992) 1007–1011.

[69] R. Goswami, T. Jeon, H. Nagaraj, S. Zhai, V.M. Rotello, Accessing Intracellular Targets through Nanocarrier-Mediated Cytosolic Protein Delivery, Trends in Pharmacological Sciences 41(10) (2020) 743–754.

[70] D. Morshedi Rad, M. Alsadat Rad, S. Razavi Bazaz, N. Kashaninejad, D. Jin, M. Ebrahimi Warkiani, A Comprehensive Review on Intracellular Delivery, Advanced Materials 33(13) (2021) 2005363.

[71] A. Eroglu, M.J. Russo, R. Bieganski, A. Fowler, S. Cheley, H. Bayley, M. Toner, Intracellular trehalose improves the survival of cryopreserved mammalian cells, Nature Biotechnology 18(2) (2000) 163–167.

[72] S. Stewart, X. He, Intracellular Delivery of Trehalose for Cell Banking, Langmuir 35(23) (2019) 7414–7422.

[73] T.M. O’Shea, Y. Ao, S. Wang, Y. Ren, A. Cheng, R. Kawaguchi, V. Swarup, M.V. Sofroniew, Border-forming wound repair astrocytes, bioRxiv (2023) 2023.08.25.554857.

[74] K. Zukor, S. Belin, C. Wang, N. Keelan, X. Wang, Z. He, Short hairpin RNA against PTEN enhances regenerative growth of corticospinal tract axons after spinal cord injury, The Journal of neuroscience : the official journal of the Society for Neuroscience 33(39) (2013) 15350–15361.

[75] B.A. Sorg, S. Berretta, J.M. Blacktop, J.W. Fawcett, H. Kitagawa, J.C.F. Kwok, M. Miquel, Casting a Wide Net: Role of Perineuronal Nets in Neural Plasticity, The Journal of Neuroscience 36(45) (2016) 11459–11468.

[76] T.M. O’Shea, Y. Ao, S. Wang, A.L. Wollenberg, J.H. Kim, R.A. Ramos Espinoza, A. Czechanski, L.G. Reinholdt, T.J. Deming, M.V. Sofroniew, Lesion environments direct transplanted neural progenitors towards a wound repair astroglial phenotype in mice, Nature Communications 13(1) (2022) 5702.

