## Supplementary Information for "Trehalose-based coacervates for local bioactive protein delivery to the central nervous system"

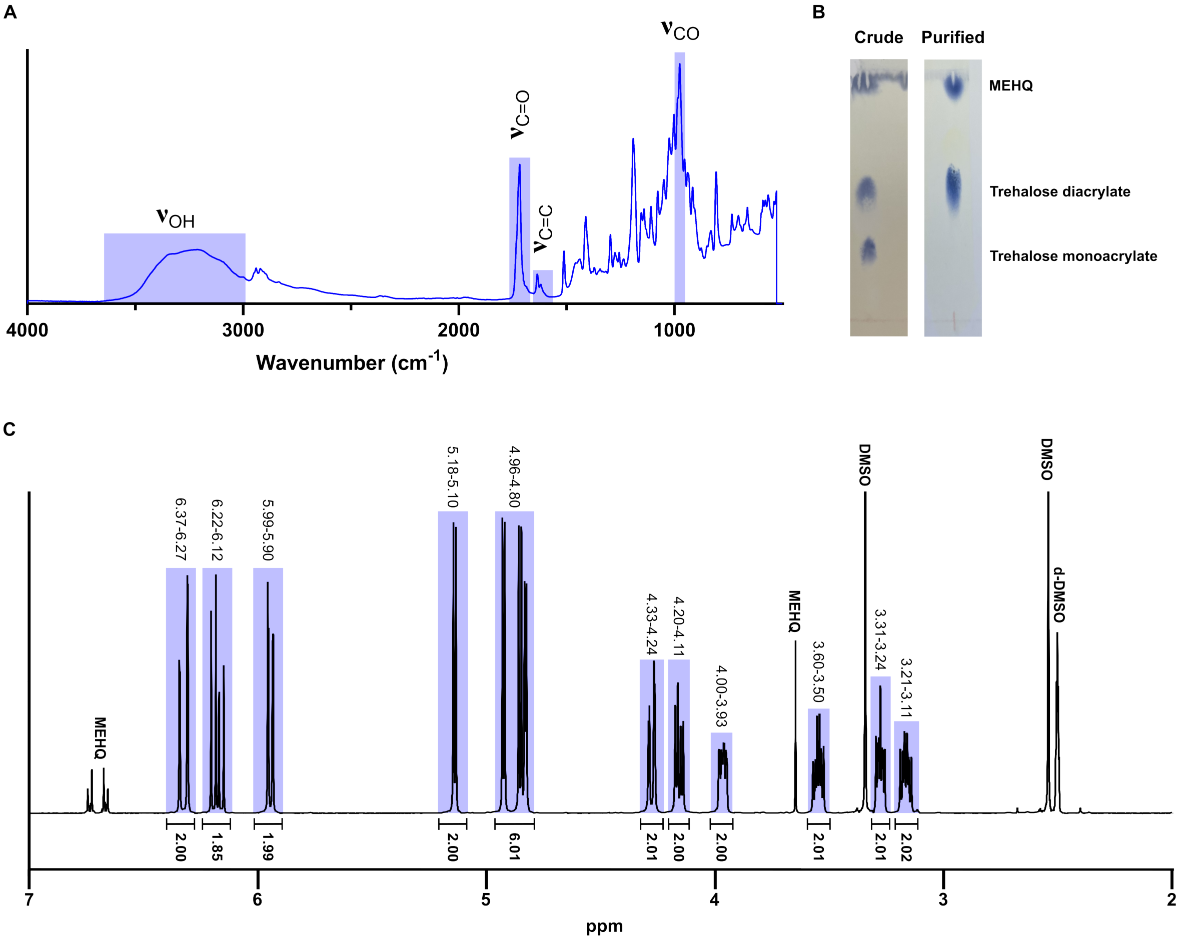

**Supplementary Figure 1**. Trehalose diacrylate characterization.
A) FTIR of purified trehalose diacrylate with characteristic peaks highlighted.
B) TLC with CAM stain of crude reaction and flash chromatography purified trehalose diacrylate using 17:4:1 ethyl acetate:methanol:water as the mobile phase.
C) $^{1}$H NMR spectrum of purified trehalose diacrylate in dDMSO. Spectrum is referenced to dDMSO solvent peak and integrations normalized to 6.27-6.37 ppm peak.

$$f_b = \frac{2 * A2 + 3 * B3 + 1 * AR}{A2 + B3 + AR}$$

**Molar ratio of monomers**

| Oligomer | fb | A2 | B3 | AR | AR Species |
|---|---|---|---|---|---|
| bS1.71 | 1.71 | 0.5 | 1 | 2 | Sulfonate |
| bS2.00 | 2 | 1 | 1 | 1 | Sulfonate |
| bA1.71 | 1.71 | 0.5 | 1 | 2 | Quaternary amine |
| bA2.00 | 2 | 1 | 1 | 1 | Quaternary amine |
| bA2.29 | 2.29 | 1.375 | 1 | 0.25 | Quaternary amine |
| bC1.71 | 1.71 | 0.5 | 1 | 2 | Carboxylic acid |
| bC2.00 | 2 | 1 | 1 | 1 | Carboxylic acid |
| bD1.71 | 1.71 | 0.5 | 1 | 2 | Tertiary amine |
| bD2.00 | 2 | 1 | 1 | 1 | Tertiary amine |
| bD2.29 | 2.29 | 1.375 | 1 | 0.25 | Tertiary amine |

**Supplementary Figure 2.** Equation for branching functionality ($f_b$) and molar ratio of monomers for different oligomers.

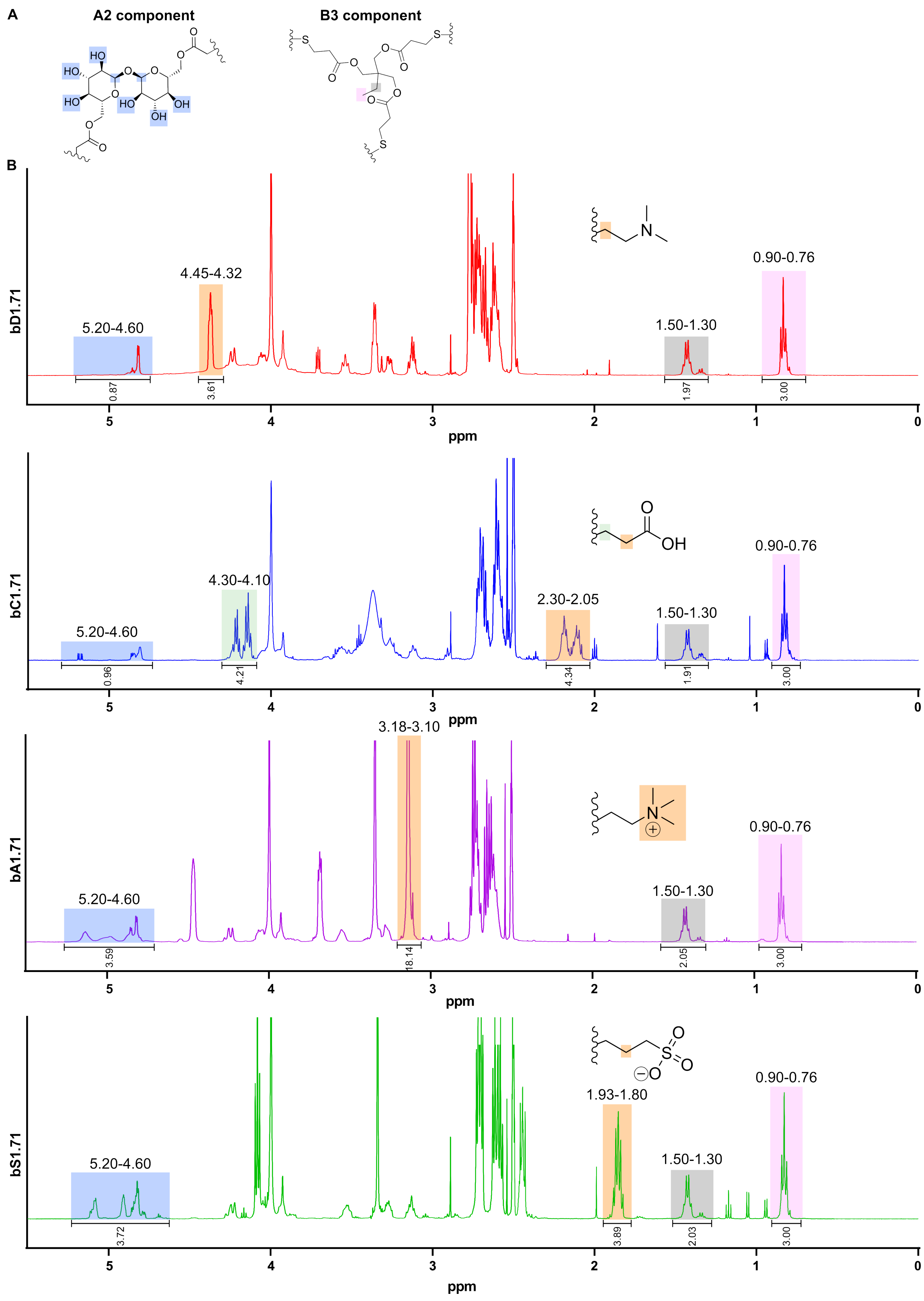

A
A2 component
B3 component
B
bD1.71
5.20-4.60
4.45-4.32
1.50-1.30
0.90-0.76
0.87
3.61
1.97
3.00
ppm
bC1.71
5.20-4.60
4.30-4.10
2.30-2.05
1.50-1.30
0.90-0.76
0.96
4.21
4.34
1.91
3.00
ppm
bA1.71
3.18-3.10
5.20-4.60
1.50-1.30
0.90-0.76
3.59
18.14
2.05
3.00
ppm
bS1.71
1.93-1.80
5.20-4.60
1.50-1.30
0.90-0.76
3.72
3.89
2.03
3.00
ppm

**Supplementary Figure 3.** $^1$H NMR spectra of branched oligomers with each end-cap at $f_b$ = 1.71 in dDMSO. All spectra referenced to dDMSO solvent peak and integrations normalized to B3 CH3 at 0.76-0.9 ppm.
A) A2 and B3 component structure with characteristic peak protons highlighted.
B) $^1$H NMR spectra for bD1.71, bC1.71, bA1.71 and bS1.71 with end-cap structure and characteristic peak protons highlighted.

| | B3 CH3 Integration | # Protons | A2 Integration | # Protons | AR Integration | # Protons | B3 Int/Prot | A2 Int/Prot | AR Int/Prot | Ex-perimental branching | Theoretical branching | Error (%) |
|---|---|---|---|---|---|---|---|---|---|---|---|---|
| Theoretical | | | | | | | **1** | **0.5** | **2** | | **1.71** | |
| bS1.71 | 3 | 3 | 3.72 | 8 | 3.89 | 2 | 1 | 0.47 | 1.95 | 1.723 | 1.71 | 0.753 |
| bC1.71 | 3 | 3 | 0.96 | 8 | 4.34 | 2 | 1 | 0.12 | 2.17 | 1.644 | 1.71 | 3.838 |
| bA1.71 | 3 | 3 | 3.59 | 8 | 18.14 | 9 | 1 | 0.45 | 2.02 | 1.707 | 1.71 | 0.184 |
| bD1.71 | 3 | 3 | 0.87 | 8 | 3.61 | 2 | 1 | 0.11 | 1.81 | 1.724 | 1.71 | 0.803 |

**Supplementary Figure 4.** Table with experimental integration values of different chemical groups allows for estimation of achieved branching and molecular weight of synthesized oligomers.

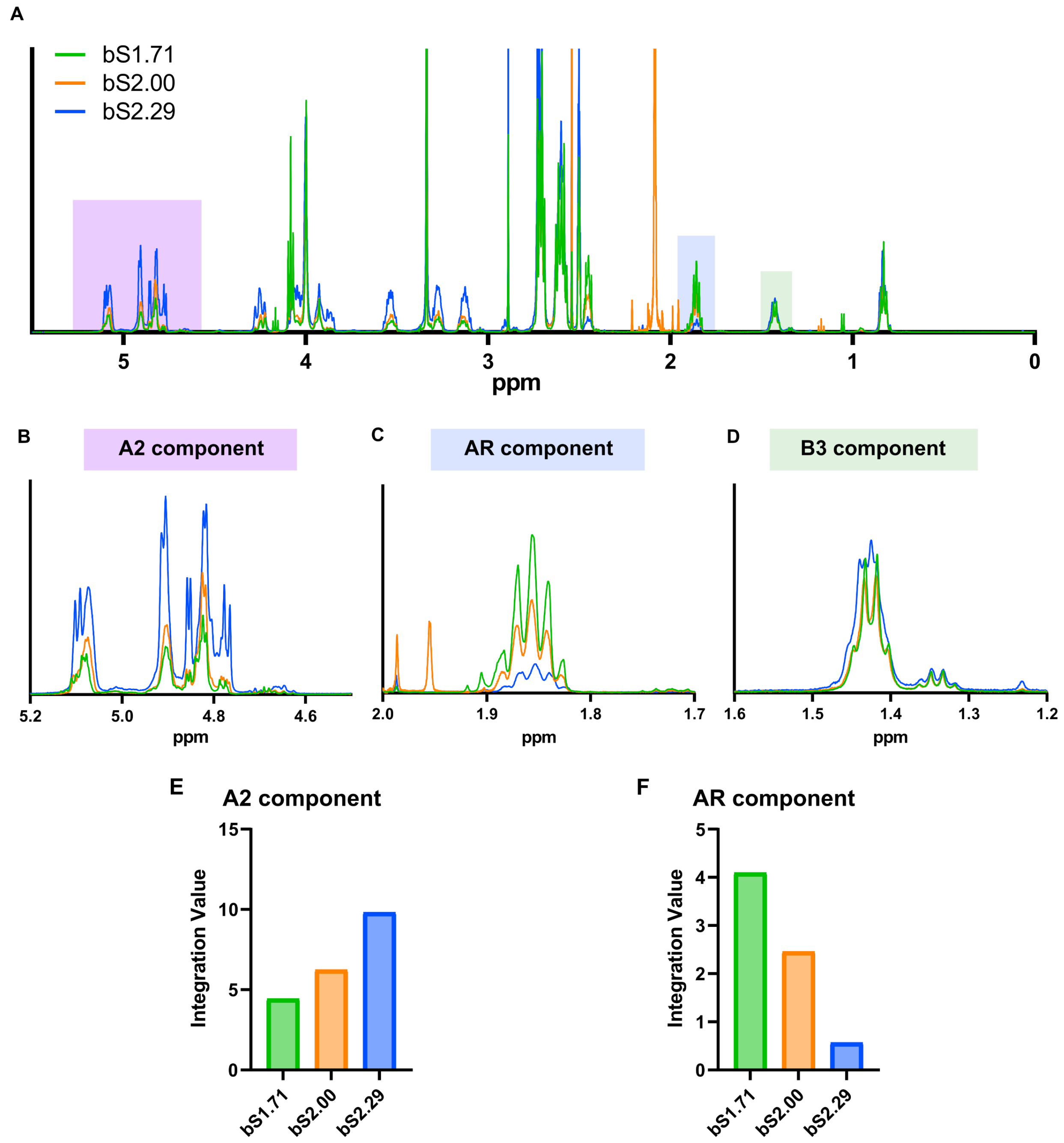

**Supplementary Figure 5.** $^{1}$H NMR spectra of differently branched bS oligomers.
A) Overlaid spectra of 3 different branching functionalities with characteristic regions highlighted. All spectra are referenced to dDMSO solvent peak and normalized to integration of B3 CH3 at 0.8 ppm.
B-D) A2, AR, and B3 regions of NMR spectra, respectively.
E, F) Integration values for differently branched oligomers of A2 and AR regions, respectively.

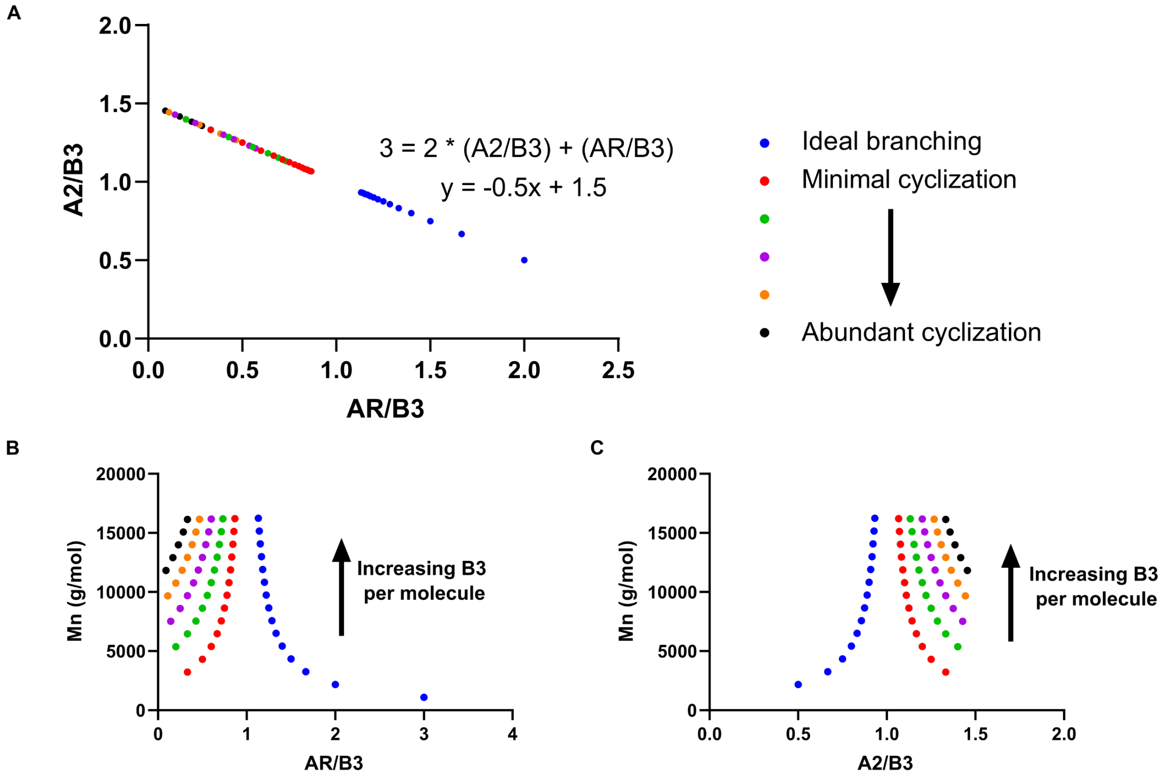

D

| | | bS1.71 | bS2.00 | bS2.29 |
|---|---|---|---|---|
| Raw integration | B3 | 2.03 | 2.04 | 2.02 |
| | AR | 4.1 | 2.46 | 0.57 |
| | A2 | 4.44 | 6.24 | 9.82 |
| Normalized to proton count | B3 | 1.015 | 1.02 | 1.01 |
| | AR | 2.05 | 1.23 | 0.285 |
| | A2 | 0.555 | 0.78 | 1.2275 |
| Experimental | AR/B3 | 2.02 | 1.21 | 0.28 |
| Theoretical (Ideal) | A2/B3 | 0.49 | 0.90 | 1.36 |
| Assuming ideal or least cyclization | Mn (g/mol) | 2177 | 10827 | 11838 |
| Higher cylization | Mn (g/mol) | - | - | 15067 |

**Supplementary Figure 6.** In the A2+B3+AR reaction scheme, the ideal branching is defined such that for n B3 monomers, there are n-1 A2 monomers. We assume full thiol and acrylate consumption. Cyclization increases as, for n B3 monomers, there are n or more A2 monomers.
A) There is a linear relationship between the molar ratios of AR/B3 and A2/B3. Different amounts of cyclization range different ratios, with multiple points of the same color indicating different numbers of B3 monomers per molecule.
B, C) As the number of B3 monomers per molecule increases, the AR/B3 and A2/B3 ratios approach 1. For AR/B3, increased cyclization shifts the ratio to the left, indicating less AR end-cap density, while the opposite is true for increased cyclization on the A2/B3 relationship.

D) We use the ratio of monomers determined from NMR to estimate the Mn for differently branched oligomers. Given the experimental AR/B3 ratio, we devised the ideal A2/B3 ratio such that all thiols and acrylates are consumed (following the linear relationship in A). For bS1.71 and bS2.00, with the ratio of A2/B3 < 1, we can assume ideal branching which allows us to determine Mn using our ideal branching curve – 2.2 kDa and 10.8 kDa respectively. For bS2.29, with the A2/B3 ratio > 1, we can devise a range of Mn based on its ratios that fit a number of differently cyclized molecules – approximately 11.8-15.1 kDa.

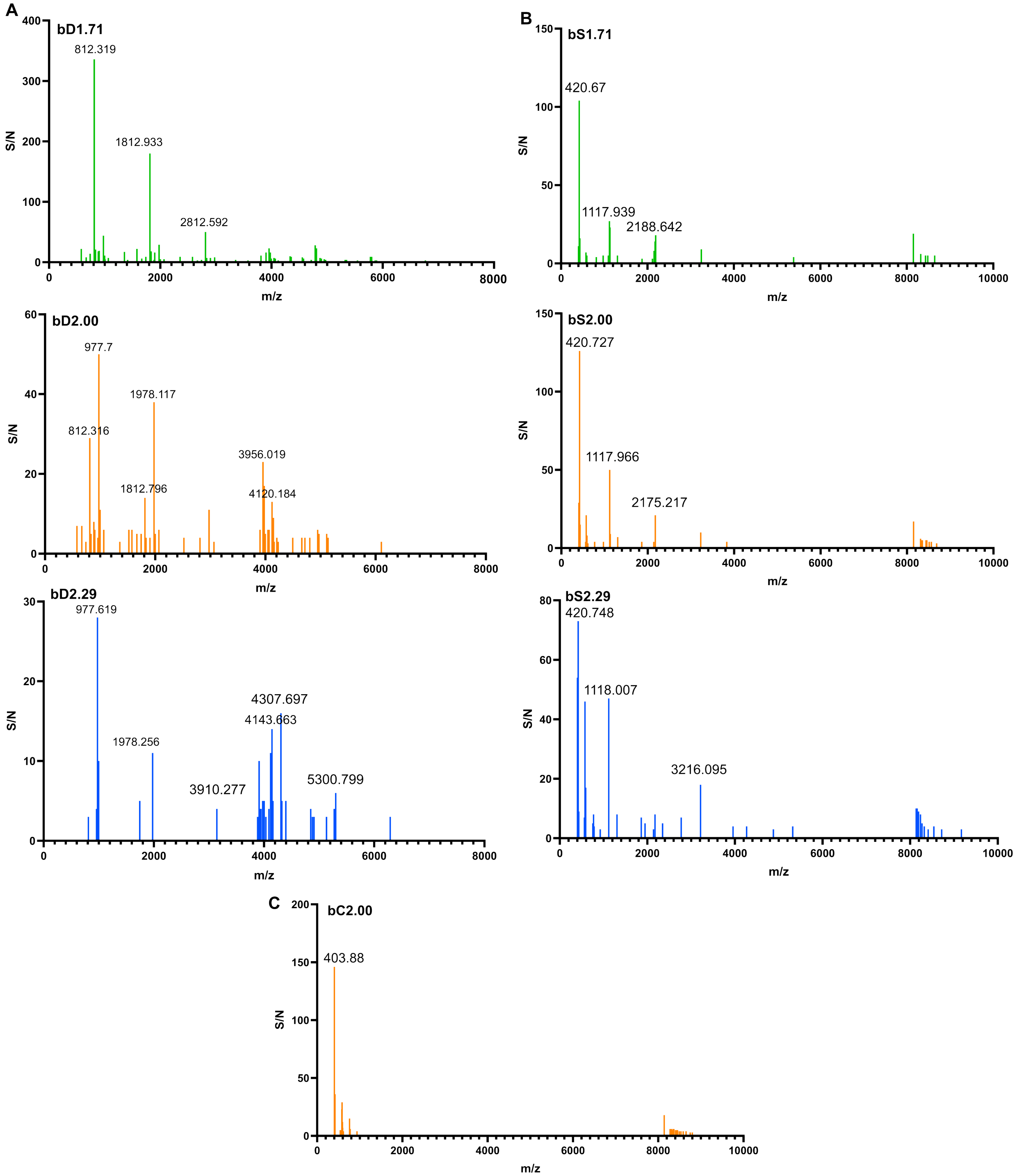

**Supplementary Figure 7.** MALDI-TOF measurements of branched oligomers.

A) Spectra of bD oligomers at 3 different branching ratios. For bD1.71, 2812 m/z represents a structure composed of a B3:A2:AR ratio = 3:2:5. For bD2.00, 3956 m/z represents a structure composed of B3:A2:AR = 4:4:4. For bD2.29, 4143 m/z represents an oligomer structure of B3:A2:AR = 4:5.5:1 - 71 Da (a segment of the DMAEA end-cap).

B) Spectra of bS oligomers at 3 different branching ratios. The constitutive charge of the bS

oligomers causes a shift towards low m/z.
C) Spectrum of bC2.00 oligomer. The carboxylic acid end-cap group did not ionize by MALDI-TOF.

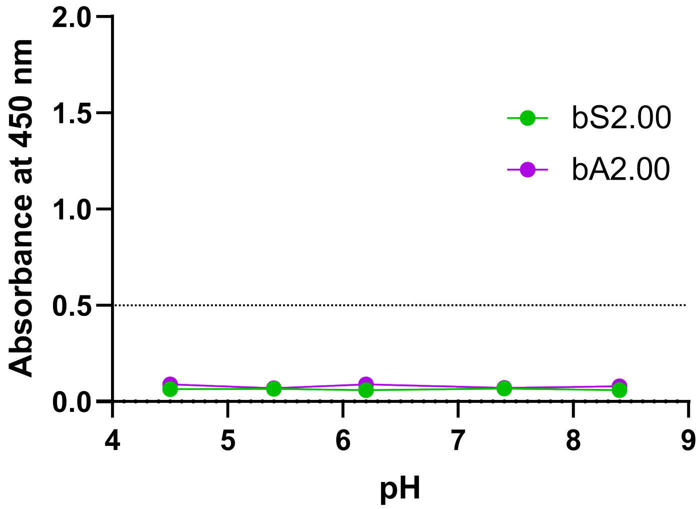

**Supplementary Figure 8.** pH dependent changes in solubility of constitutively charged oligomers (10 mg/ml). Threshold Abs450 = 0.5 set at point of visible turbidity. Graph shows mean ± SEM with n=3.

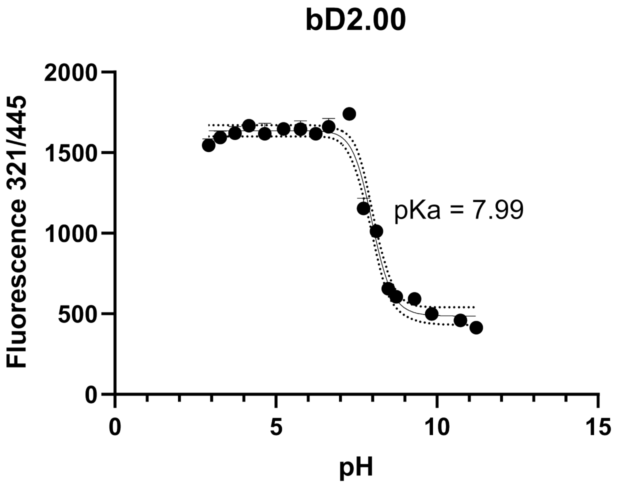

**Supplementary Figure 9.** pKa determination using TNS assay of bD2.00 oligomer at 4 mg/ml. Graph shows mean ± SEM with n=3. Points were fit to a 4PL sigmoidal curve and pKa was determined to be 7.99.

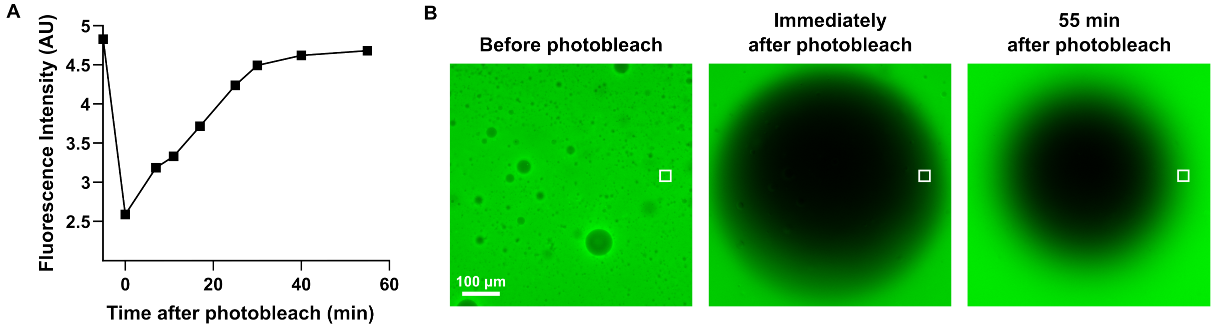

**Supplementary Figure 10.** Fluorescence recovery after photobleaching (FRAP) of fluorescein-labeled coacervates.
A) FITC intensity of bulk coacervate at 250 µm from center of photobleached spot over 1 hour (area of measure represented by white box in B).
B) Representative images of fluorescence recovery over 1 hour of a ~600 µm diameter photobleached area of coacervate.

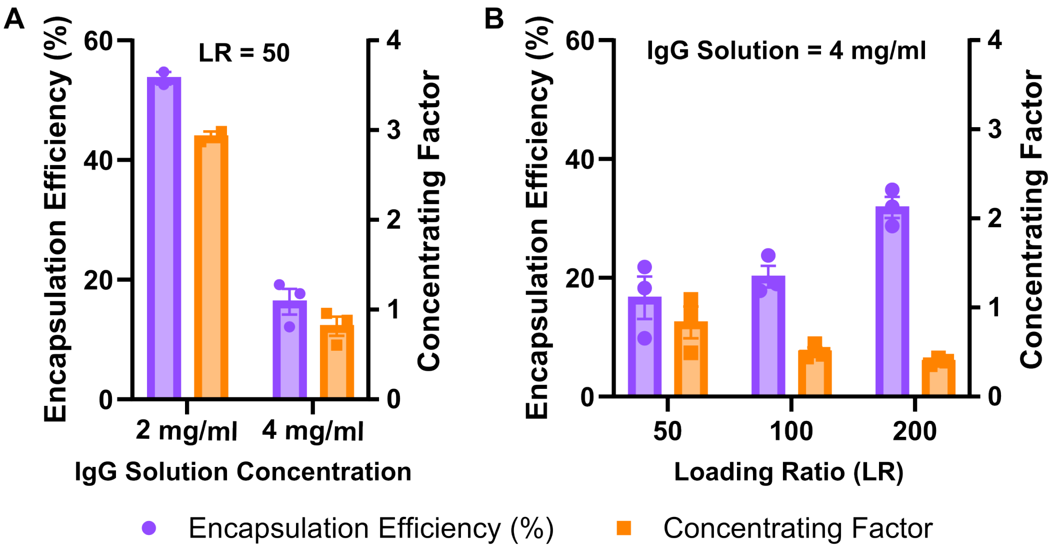

**Supplementary Figure 11.** IgG encapsulation and concentrating factor.
A) FITC-IgG encapsulation efficiency and concentrating factor at different protein solution concentrations with the same coacervate formulation (LR = 50, C:D = 1:1).
B) FITC-IgG encapsulation efficiency and concentrating factor at different loading ratios (C:D = 1:1) with the same protein solution concentration (4 mg/ml).

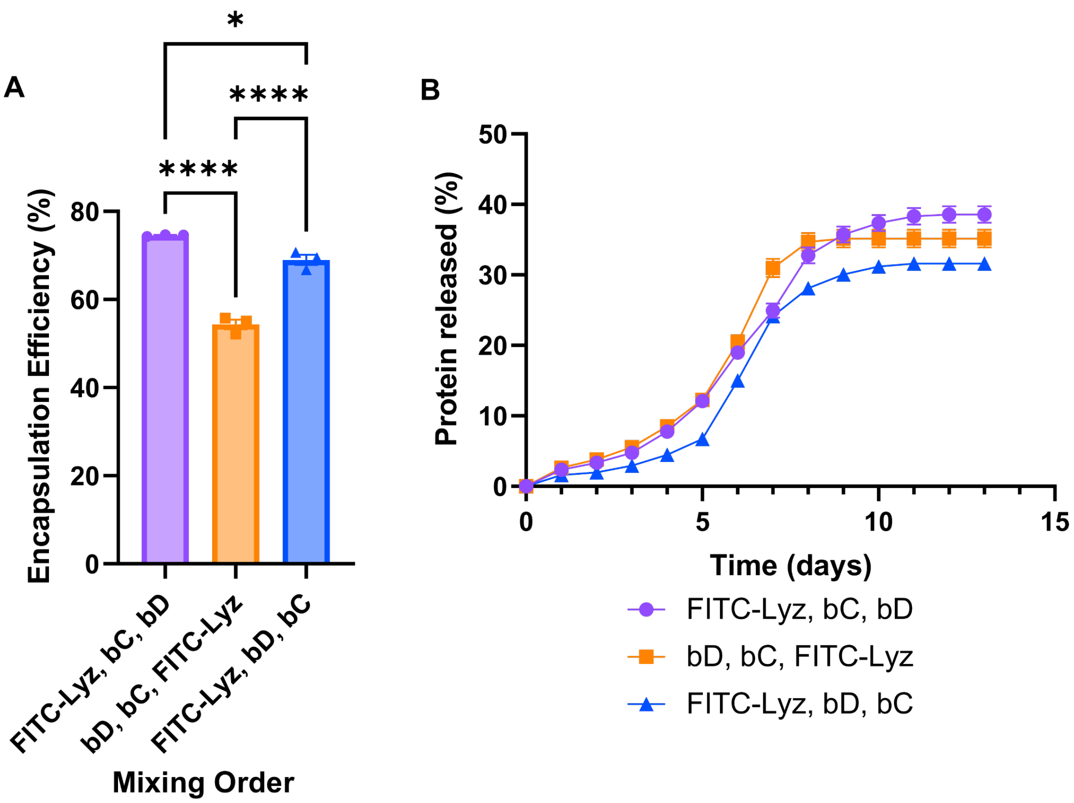

**Supplementary Figure 12.** Order of incorporating protein and positively and negatively charged oligomers affects encapsulation and release profiles. Formulation was set at loading ratio (LR) = 50 with 20 mg of total oligomer, 0.4 mg of FITC-Lyz and C:D = 1.25:1.

A) Encapsulation efficiency of FITC-Lyz with different mixing orders. Ordinary one-way ANOVA with Tukey's multiple comparisons test, * $P < 0.05$, **** $P < 0.0001$.

B) Release profiles of FITC-Lyz over 2 weeks with different mixing orders.

Graphs show mean ± SEM with n=3.

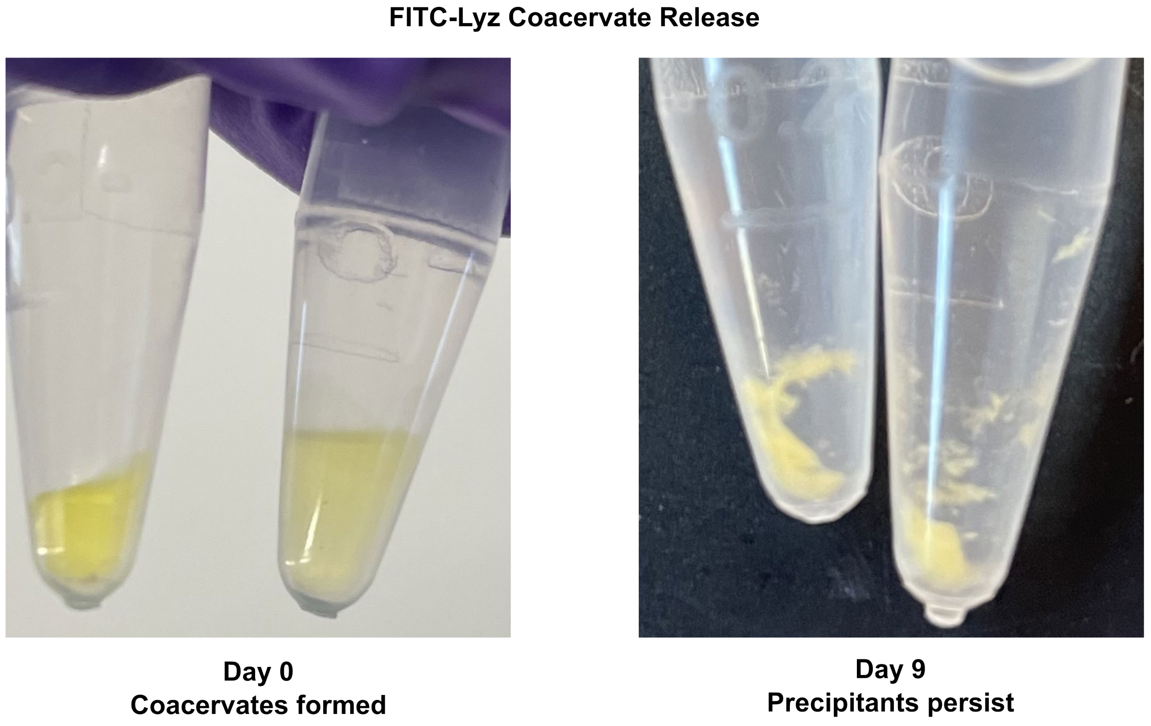

**Supplementary Figure 13.** Images of FITC-Lyz coacervates upon formation and after 9 days of release shows transition from liquid phase to solid precipitate phase.

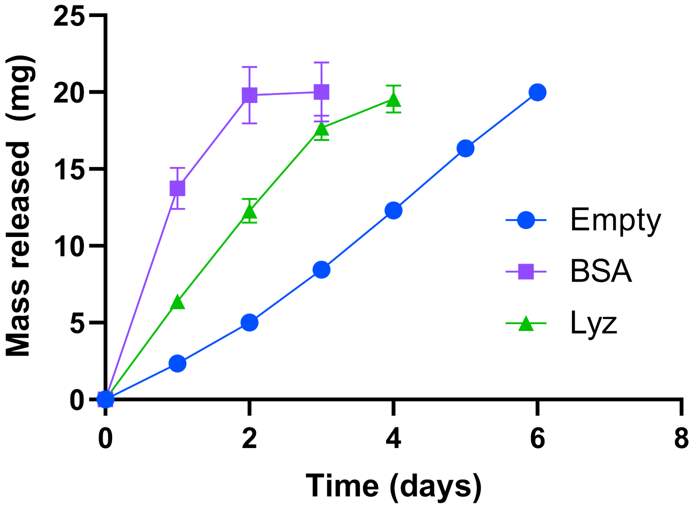

**Supplementary Figure 14.** Comparison of mass release of fluorescein labeled coacervate when empty vs. when loaded with unlabeled BSA or Lyz. All formulations are 20 mg of total oligomer with LR = 50 for protein coacervates and C:D = 1:1. Graph shows mean ± SEM with n=3.

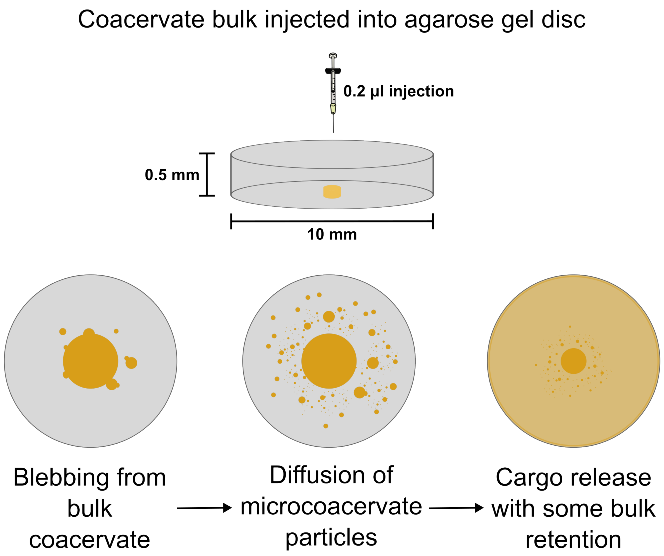

**Supplementary Figure 15.** Schematic illustrating injection and diffusion of rhodamine-dextran loaded coacervate into 0.6 wt% agarose gel phantom brain disc.

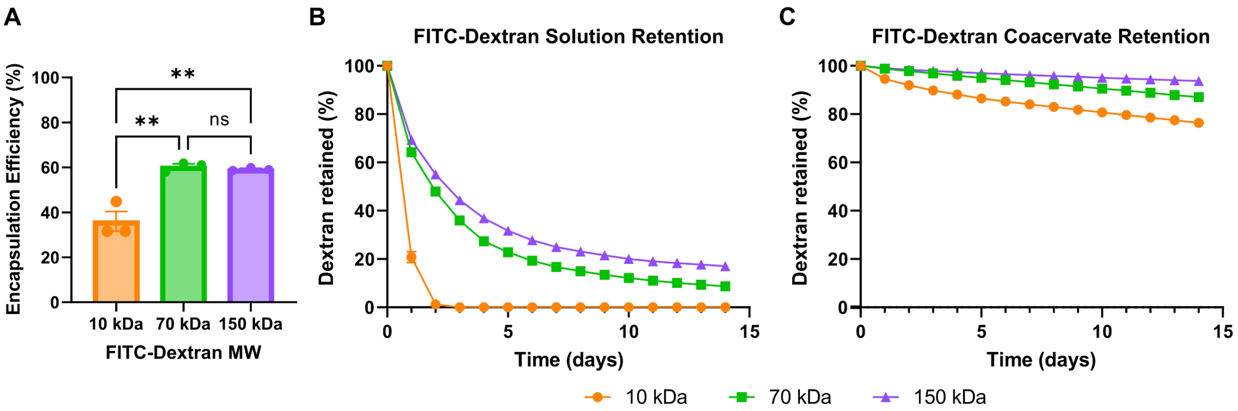

**Supplementary Figure 16.** FITC-dextran solution or coacervates were suspended in 0.6 wt% agarose gel and submerged PBS supernatant. Retention was measured as amount of FITC-dextrans not released into supernatant after 14 days with daily replacement of PBS and fluorescent read of daily collected supernatant.

A) Encapsulation efficiency of different MW FITC-dextrans in the same coacervate formulation (LR = 50, total oligomer mass = 25 mg, C:D = 1:1). Ordinary one-way ANOVA with Tukey's multiple comparisons test, ** $P < 0.01$.

B) FITC-dextrans of different MW retention in agarose gel by solution.

C) FITC-dextrans of different MW retention in agarose gel by coacervate.

Graphs show mean ± SEM with n=3.

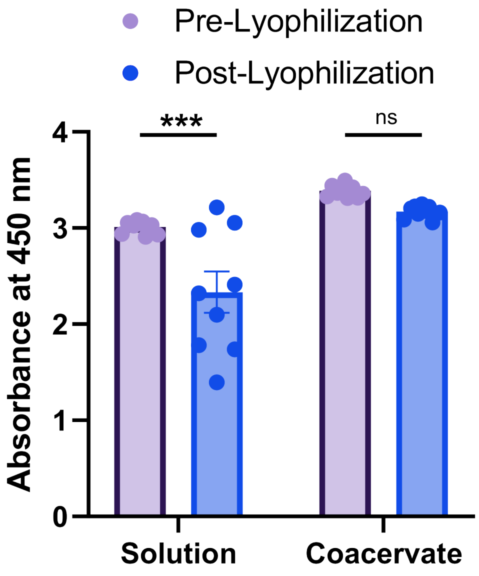

**Supplementary Figure 17.** Comparison of HRP bioactivity as measured by absorbance at 450 nm before and after lyophilization and resuspension either in coacervate or solution. 2way ANOVA with Tukey's multiple comparisons test, *** $P < 0.001$. Graph mean ± SEM with n=9.

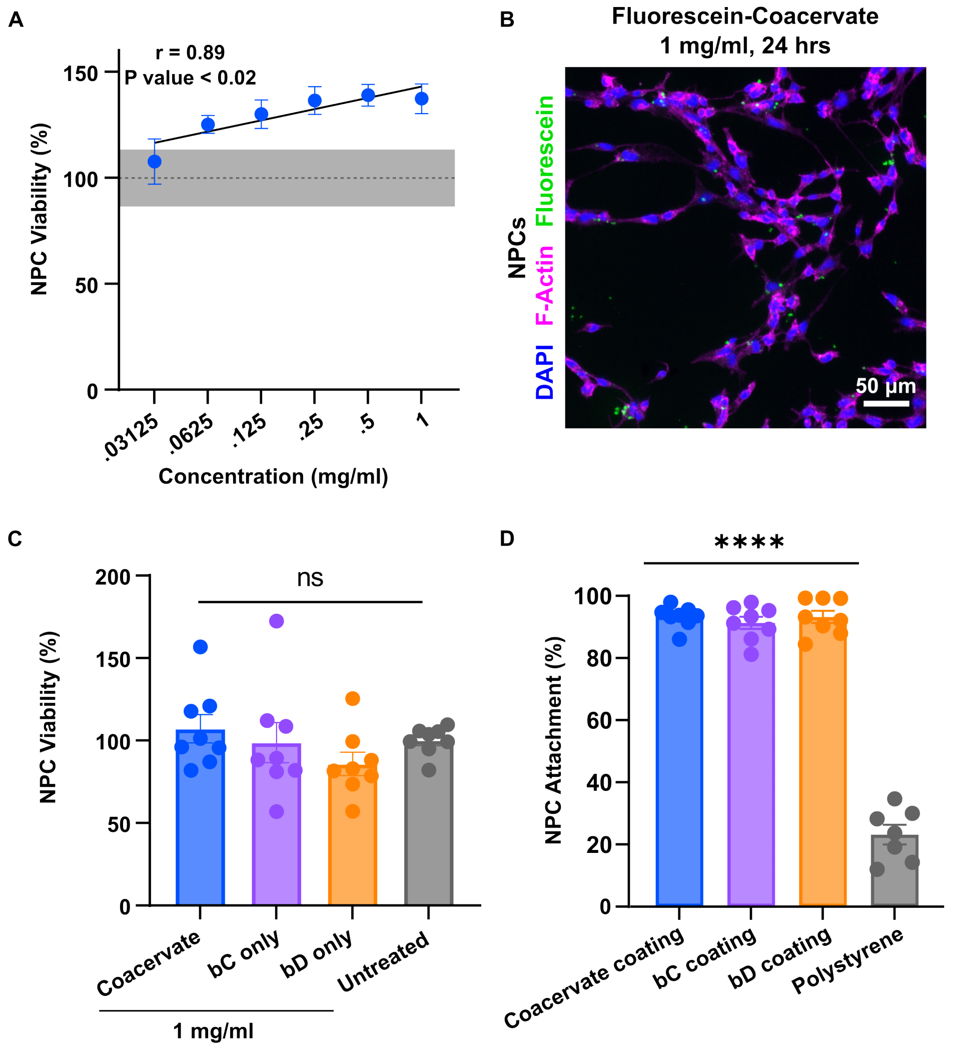

**Supplementary Figure 18.** NPC interactions with coacervates.
A) Dose response of neural progenitor cells (NPC) to neutral coacervates after 24 hours of treatment measured by Calcein AM assay. Graph shows mean ± SEM for n=6 with linear correlation fit to $\log_2$(coacervate concentration). Pearson r = 0.89, P = 0.017. Dotted line and shading represent control mean ± SEM.
B) Image of NPC treated with fluorescein-coacervates (1 mg/ml) for 24 hours. Stained with DAPI and phalloidin.
C) Viability of neural progenitor cells (NPC) with coacervates and individual oligomers all at 1 mg/ml; measured by Calcein AM assay.
D) NPC cell attachment to coacervate and individual oligomers (2 mg/ml) compared to untreated (polystyrene) and benchmarked against tissue culture treated plastic, measured by Calcein AM assay. Comparisons made to polystyrene control.
(C, D) Graphs show mean ± SEM with n=7-8 with ordinary one-way ANOVA with Tukey's multiple comparisons test. Not significant (ns), **** $P < 0.0001$.

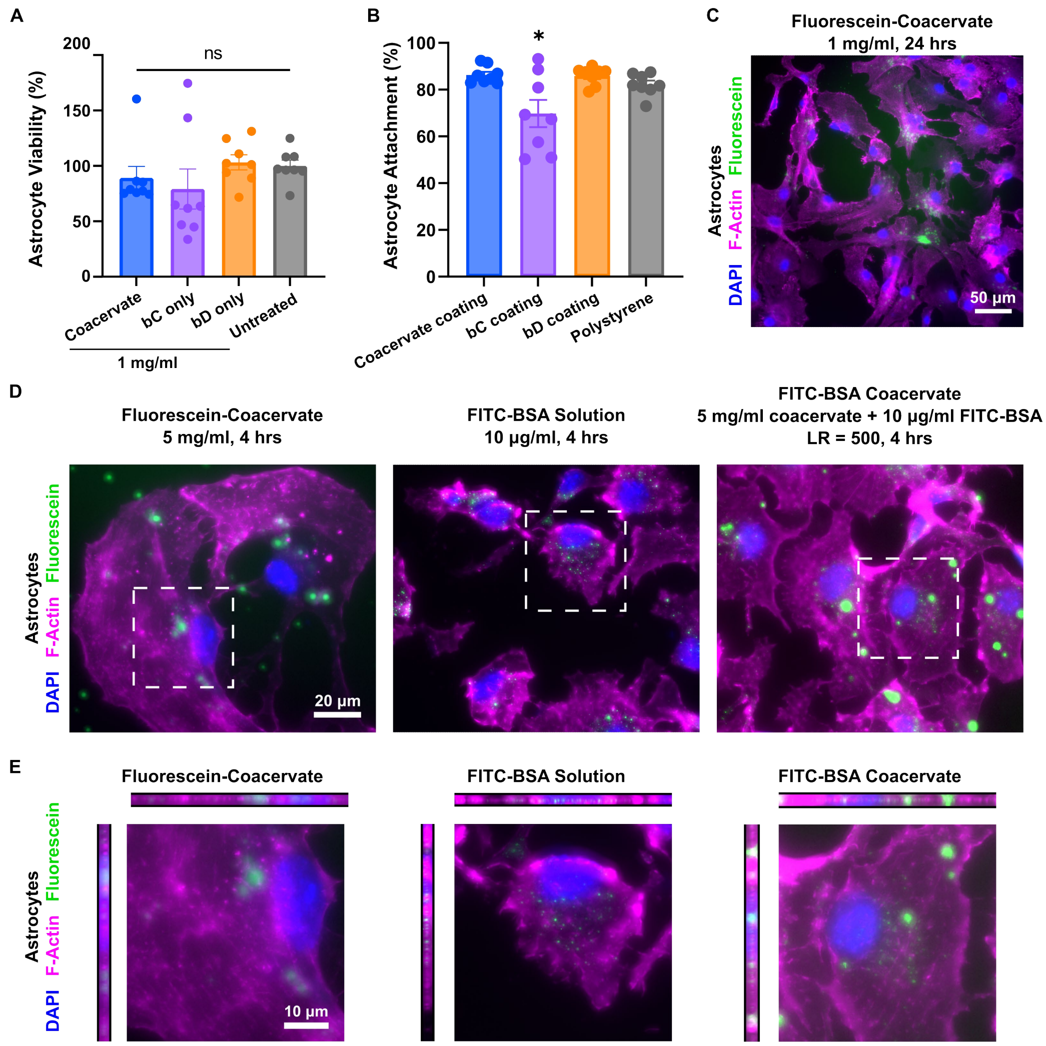

**Supplementary Figure 19.** Astrocyte interactions with coacervates.
A) Viability of astrocytes with coacervates and individual oligomers all at 1 mg/ml; measured by Calcein AM assay.
B) Astrocyte cell attachment to coacervate and individual oligomers (2 mg/ml) compared to untreated (polystyrene) and benchmarked against tissue culture treated plastic, measured by Calcein AM assay. Comparison made to polystyrene control.
C) Image of astrocytes treated with fluorescein-coacervates (1 mg/ml) for 24 hours. Stained with DAPI and phalloidin.
D) Images of astrocytes treated with fluorescein-coacervates (5 mg/ml), FITC-BSA solution (10 µg/ml), and FITC-BSA coacervates (10 µg/ml FITC-BSA, LR = 500) for 4 hours. Stained with DAPI and phalloidin. White boxes represent area of orthogonal projection in E. Cultured astrocytes treated with FITC-BSA loaded coacervates for 4 or 24 hours did not increase the intracellular BSA concentration within the cell cytoplasm compared to FITC-BSA solution controls and micro-sized coacervates persisted on the surface of cells throughout the 24-hour period of evaluation. We suspect that the lack of intracellular delivery or endocytosis of coacervates to astrocytes could be in part the result of an insufficient period of incubation or

direct binding of cell culture media constituents, such as serum proteins, to the surface of coacervate droplets, but future studies will be required to rigorously dissect this hypothesis.
E) Orthogonal projections of astrocytes treated with either fluorescein-labelled coacervates, FITC-BSA solution of FITC-BSA loaded coacervates.
(A, B) Graphs show mean ± SEM with n=7-8 with ordinary one-way ANOVA with Tukey's multiple comparisons test. Not significant (ns), * $P < 0.05$.

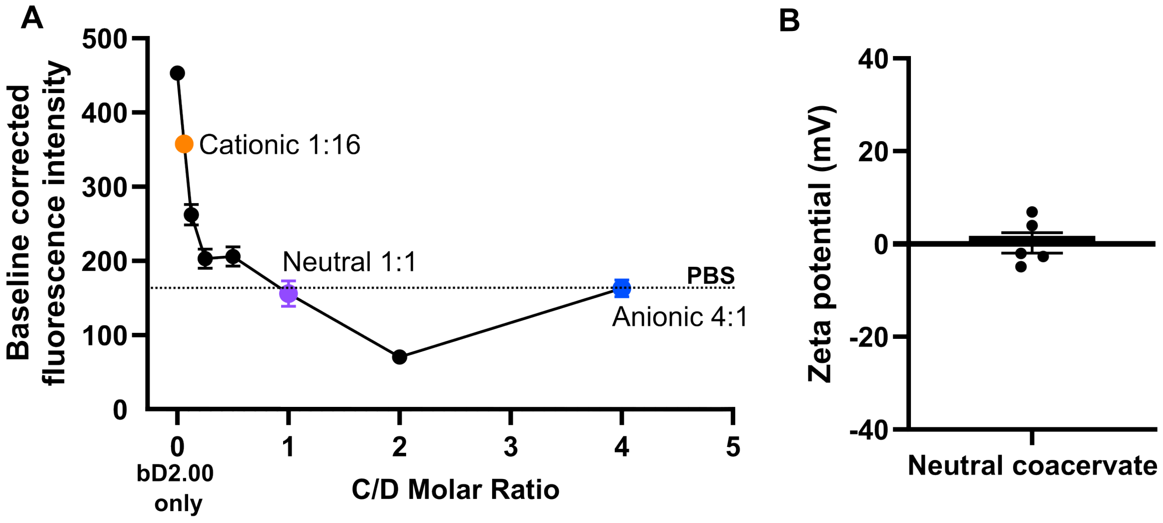

**Supplementary Figure 20.** Surface charge of coacervates.
A) Excess positive surface charge on charge-imbalanced coacervates measured by TNS assay. Dotted line indicates PBS baseline fluorescence. Colored dots indicate formulations used in *in vitro* studies.
B) Zeta potential measurement of neutral (C:D = 1:1) coacervate.

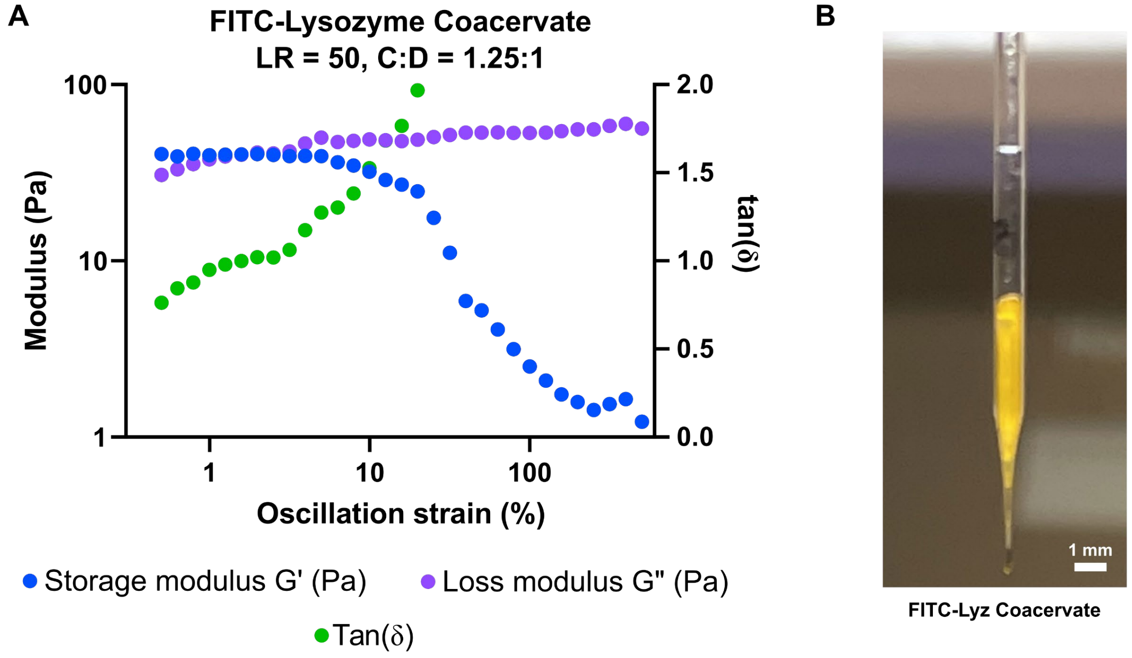

**Supplementary Figure 21.** Protein loaded coacervates are readily loaded into micropipette for injection.

A) Dynamic rheological oscillatory strain sweep measurements for FITC-Lyz coacervate showing storage modulus (G'), loss modulus (G"), and tan(δ).

B) FITC-Lyz coacervate loaded into glass micropipette (outer diameter = 1 mm, inner diameter = 0.5 mm).

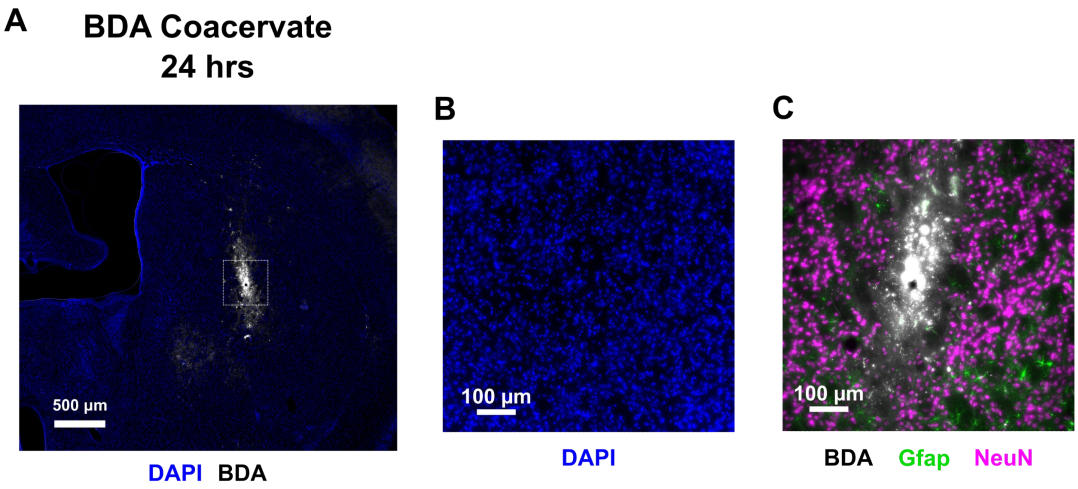

**Supplementary Figure 22.** 100% v/v biotinylated dextran amine (BDA) coacervate injection at 24 hours in healthy mouse striatum.

A) Low magnification image of injection into healthy mouse striatum.

B) High magnification image of DAPI stain at injection site.

C) High magnification image showing surrounding neural tissue (neurons = NeuN, astrocytes = Gfap) around injection.

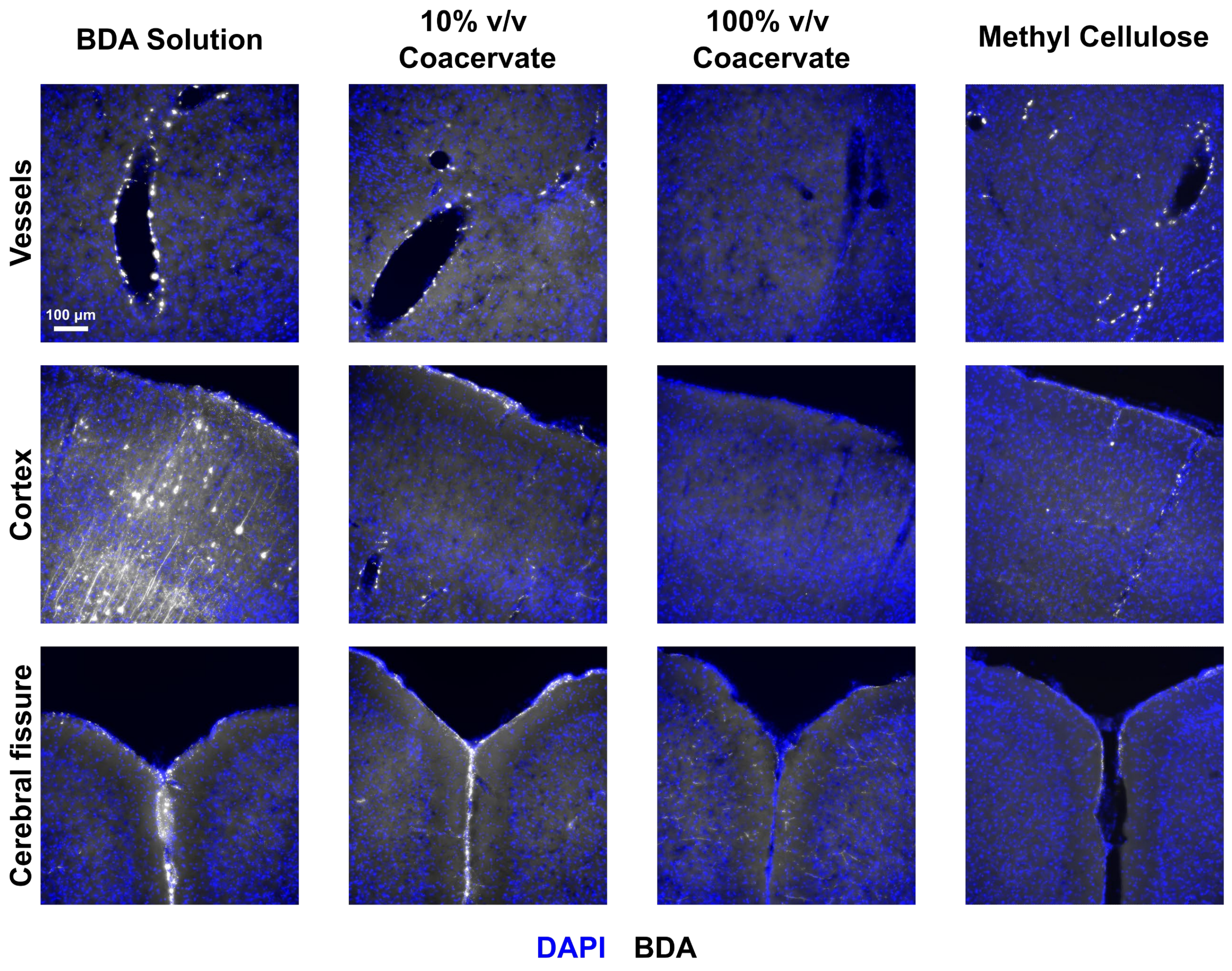

**Supplementary Figure 23.** Images of different regions of mouse brain showing BDA distribution in perivascular spaces, cortex, and meninges with different BDA carriers.

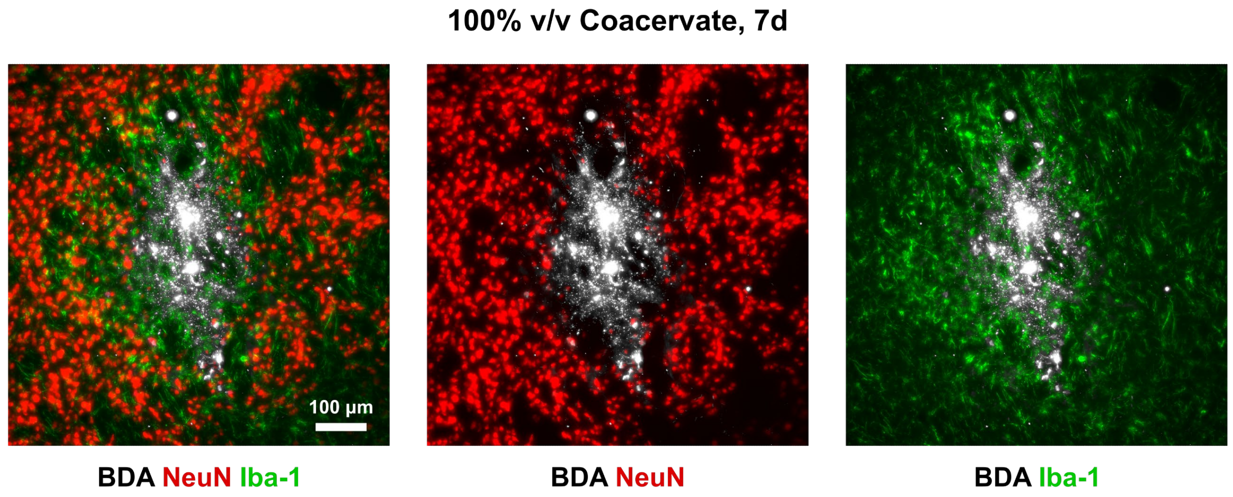

**Supplementary Figure 24.** Images of injection site of 100% v/v BDA coacervate into healthy mouse striatum at 7d and uptake by neurons (NeuN) and microglia (Iba-1).

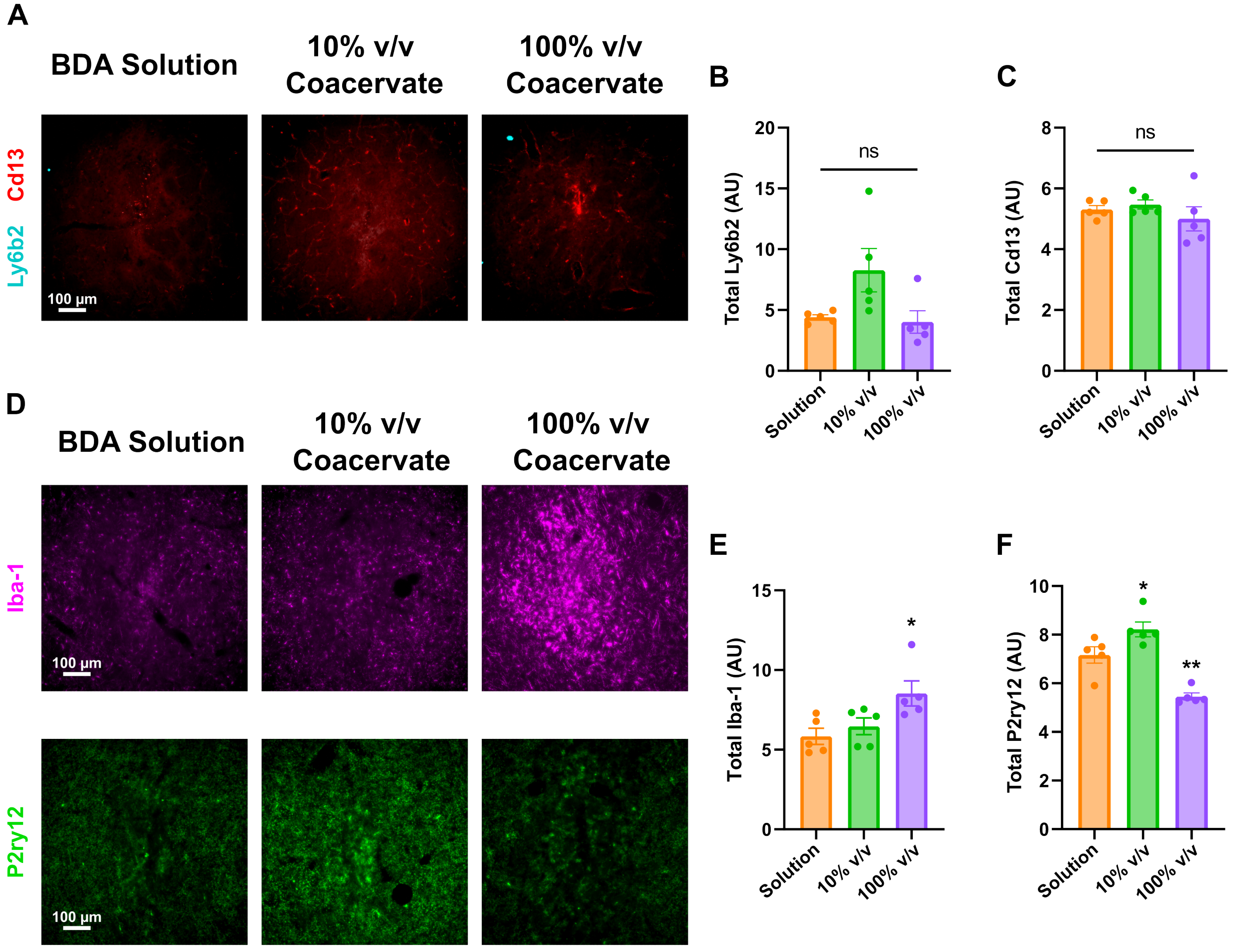

**Supplementary Figure 25.** Images and quantification of immune response at 7d to BDA coacervates.

A) Images of injection center in healthy mouse striatum 7d after injection of BDA solution, 10% v/v coacervate and 100% v/v coacervate stained for neutrophils (Ly6b2) and myeloid lineage cells (Cd13).

B) Quantification of total Ly6b2 within 1000 μm radius of injection.

C) Quantification of total Cd13 within 1000 μm radius of injection.

D) Microglia response at injection center.

E) Quantification of total Iba-1 within 1000 μm radius of injection.

F) Quantification of total P2ry12 within 1000 μm radius of injection.

All statistical tests are ordinary one-way ANOVA with Tukey's multiple comparisons test, * $P < 0.05$, ** $P < 0.01$. Graphs show mean ± SEM with n=5.

### 100% v/v Coacervate, 7d

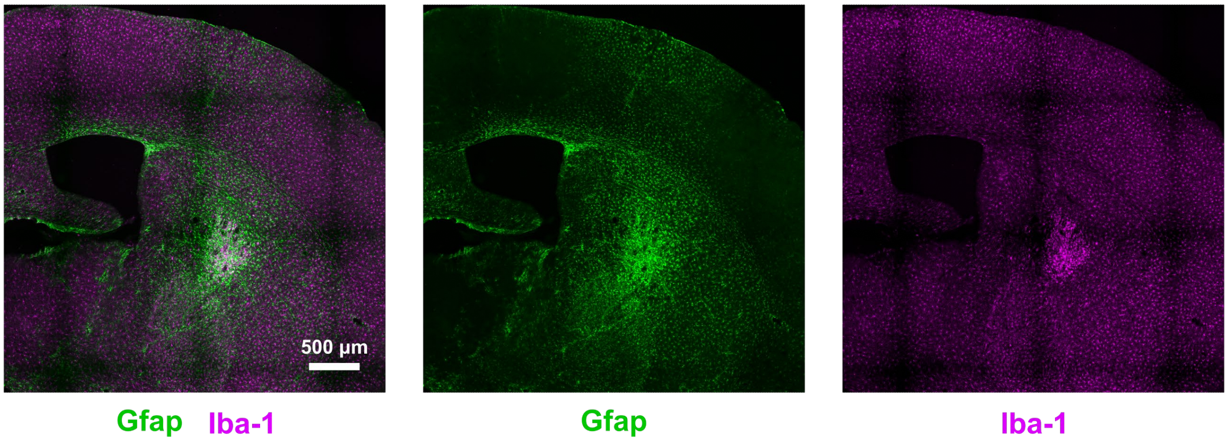

**Supplementary Figure 26.** Images of injection site of 100% v/v BDA coacervate into healthy mouse striatum at 7d and infiltration by astrocytes (Gfap) and microglia (Iba-1).

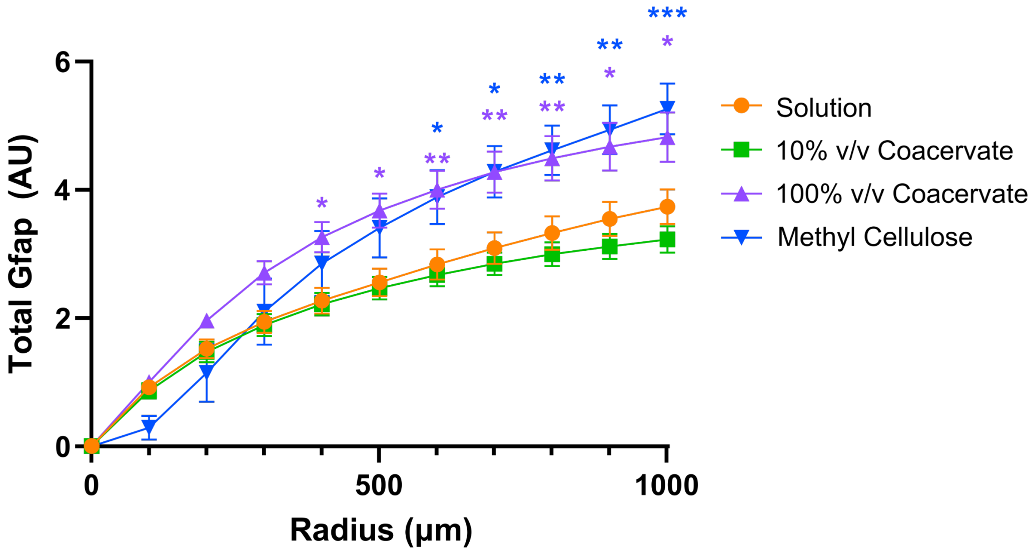

**Supplementary Figure 27.** Cumulative Gfap expression from injection center to 1000 µm radius in healthy mouse striatum at 7d after injection. 2way ANOVA with Tukey's multiple comparisons test, * $P < 0.05$, ** $P < 0.01$, *** $P < 0.001$. Graph shows mean ± SEM with n=4-5.

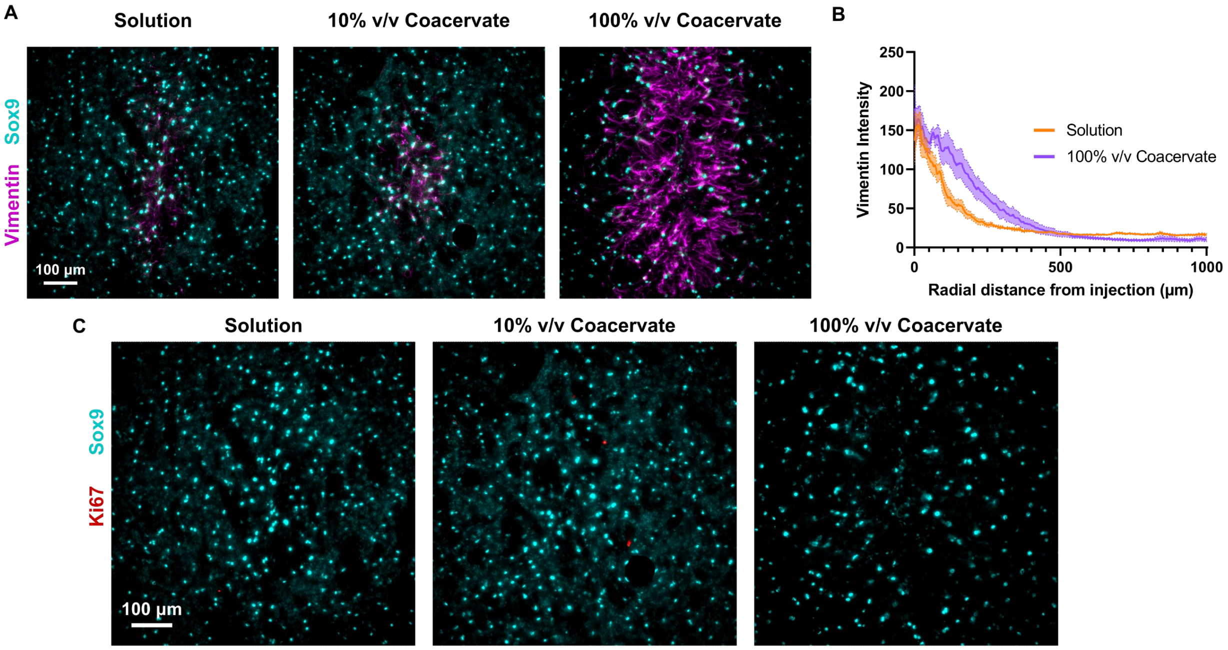

**Supplementary Figure 28.** Images and quantification of astrocyte reactivity and proliferation.
A) Images of astrocyte response at injection site of solution, 10% v/v coacervate and 100% v/v coacervate at 7d.
B) Radial angle profile of Vimentin staining of solution and 100% v/v coacervate from injection center. Graph shows mean ± SEM with n=5.
C) Images of injection site show little to no Ki67+ astrocytes at 7d in any condition.

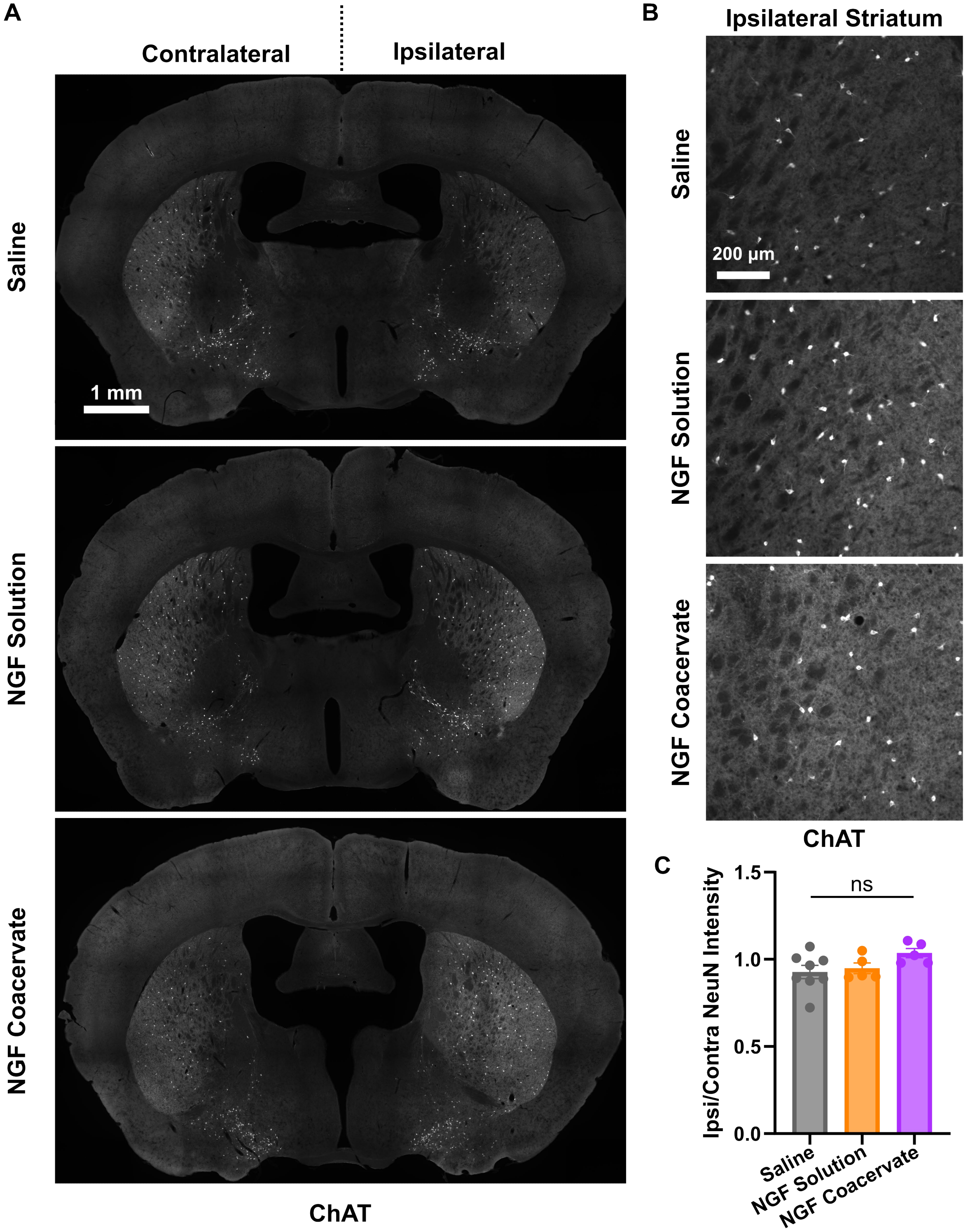

**Supplementary Figure 29.** NGF solution and coacervates increase ChAT neuron size and intensity without causing neural damage.
A) Low magnification images of whole section choline acetyltransferase (ChAT) staining and detailed images of ipsilateral striatum for saline, NGF solution, and NGF coacervate.

B) Higher magnification image of striatal ChAT neurons.
C) Quantification of change in NeuN intensity from the ipsilateral to contralateral sides of the brain. Graph shows mean ± SEM with n=5-8 with ordinary one-way ANOVA with Tukey's multiple comparisons test. Not significant (ns).
